# Contrasting microhaplotype diversity responses of saproxylic beetles to forest management

**DOI:** 10.64898/2026.11.25.769104

**Authors:** Nermeen R. Amer, Rama S. Krovi, Alicja Wierzbicka, Tomasz Jaworski, Adrian Smolis, Radosław Plewa, Marcin Kadej, Maria Oczkowicz, Łukasz Kajtoch

## Abstract

Genetic responses to forest management may vary substantially among saproxylic beetles because species differ in habitat requirements, dispersal and ecological specialization. Here, we evaluated whether microhaplotypes reconstructed from Double Digest Restriction-site Associated DNA sequencing (ddRADseq) loci provide additional locus-level genetic information and reveal species-specific associations with forest protection, management and habitat characteristics across 12 saproxylic beetle species sampled in eight forest complexes in Poland. Phased combinations of linked Single Nucleotide Polymorphism (SNPs) within RAD loci were analysed as multi-allelic microhaplotypes. Observed microhaplotype diversity was compared with analytical expectations under linkage equilibrium to assess how diversity differed from that expected if constituent SNP positions were independent. Observed microhaplotype diversity was consistently lower than expected under linkage equilibrium across all 12 species, with the magnitude of the difference varying among species. Patterns of population differentiation inferred from SNP-based F_ST_ and microhaplotype-based Φ_ST_ were significantly concordant in all species, although the strength of correspondence varied considerably among taxa. Individual microhaplotype homozygosity (F_HAP_) showed heterogeneous associations with site protection and management categories, with no consistent directional response across species. Sensitivity analyses restricted to the NE Poland detected no significant management associations after correction for multiple testing. Associations between site-level microhaplotype diversity and habitat characteristics also varied among species. Species traits showed suggestive associations with management-related changes in F_HAP_, although none remained statistically significant after correction for multiple-testing. Overall, microhaplotypes retain linked within-locus genetic variation and broadly recover SNP-based patterns of population differentiation, while associations between forest context and genetic variation remain strongly species-dependent.

## Introduction

Restriction-site associated DNA sequencing (RADseq) is a reduced-representation genomic approach that repeatedly samples homologous genomic regions associated with restriction sites, allowing large numbers of polymorphic loci to be discovered and genotyped across individuals (Baird et al., 2008; Andrews et al., 2016). ddRADseq uses two restriction enzymes together with fragment-size selection to target a reproducible subset of genomic fragments and has become widely used in population genomics of non-model organisms (Peterson et al., 2012; Andrews et al., 2016). Importantly, a RAD locus is a short sequenced genomic fragment rather than intrinsically a single-SNP marker and may therefore contain several linked polymorphic sites. When these sites occur within the same sequenced region, their allelic combinations can be retained as haplotypes instead of representing each variant independently.

This feature makes multi-SNP RAD loci suitable for a microhaplotype representation. Microhaplotypes are short genomic regions containing closely linked polymorphic sites whose phased combinations define multiallelic haplotypes (Kidd et al., 2014; Kidd & Speed, 2015). Whereas individual SNPs are generally biallelic, combinations of linked SNPs can generate several alleles within a single microhaplotype locus. Baetscher et al. (2018) demonstrated that retaining these phased combinations can preserve substantially more information from short-read sequence data than reducing the same regions to individual SNPs. However, increased allelic resolution does not necessarily translate into improved performance for every downstream analysis. Its value depends on the genetic statistic or biological question being investigated.

The expected information gain should be particularly pronounced at loci containing several polymorphic sites because increasing numbers of linked variants can generate increasing numbers of haplotypic combinations. Microhaplotypes should therefore be viewed not as replacements for SNPs, but as a complementary representation that retains additional within-locus allelic information already present in sequence data.

Although microhaplotypes have been developed predominantly for forensic genetics, relationship inference and population assignment (Kidd et al., 2014; Kidd & Speed, 2015; Baetscher et al., 2018), their application to ecological and conservation-genomic questions remains comparatively limited. Reconstructing microhaplotypes from existing RADseq datasets may be particularly valuable because it can recover an additional layer of genetic information without requiring new sequencing. Nevertheless, whether this representation changes ecological interpretations or provides information beyond that obtained from conventional SNP-based analyses must be evaluated empirically rather than inferred from increased haplotype diversity alone.

This approach is potentially useful in multispecies systems such as saproxylic beetles, where species differ strongly in resource specialization, dispersal and responses to forest structure. Deadwood amount, diversity and microclimatic heterogeneity influence saproxylic assemblages (Seibold et al., 2016; Seibold et al., 2017), while even threatened species occupying the same forest landscape can depend on different deadwood resources (Jaworski et al., 2019). Such ecological heterogeneity provides a strong basis for expecting species-specific rather than uniform genetic responses to forest management.

Forest management is an important driver of biodiversity change because it modifies habitat structure, resource availability and continuity, microclimatic conditions, and landscape connectivity (Oettel & Lapin 2021). These effects are particularly relevant to saproxylic insects, which depend during at least part of their life cycle on dead or dying trees, wood-inhabiting fungi, or other organisms associated with these substrates. Saproxylic insects contribute to decomposition, nutrient cycling and forest food webs, while many species are sensitive to changes in forest management and habitat quality (Grove, 2002). Forest management can reduce availability of old trees and deadwood and alter the spatial and temporal continuity of these resources. However, deadwood is highly heterogeneous, varying with respect to tree species, dimensions, decay stage, moisture, fungal colonization and microclimate, and different components of this resource support highly diversified saproxylic assemblages (Seibold et al., 2016). Consequently, even ecologically similar saproxylic species may respond differently to the same management regime because of differences in their habitat and resource requirements (Jaworski et al., 2019).

Changes in habitat amount, quality and connectivity can ultimately influence genetic diversity and population structure. Habitat fragmentation may reduce population sizes and limit dispersal, thus increasing genetic drift and potentially reducing within-population diversity while increasing differentiation among populations. However, these responses depend strongly on species ecology, dispersal capacity, habitat specialization and demographic history. Such variation is evident among saproxylic beetles. For example, *Bolitophagus reticulatus* (Linnaeus, 1767) (Tenebrionidae) showed substantially greater genetic differentiation in a fragmented landscape than in continuous oldgrowth forest (Knutsen et al., 2000), while landscape-genetic analyses of *Elater ferrugineus* Linnaeus, 1758 (Elateridae) demonstrated the importance of habitat connectivity for dispersal among fragmented populations (Oleksa et al., 2015). More broadly, genetic patterns among European saproxylic beetles reflect both contemporary landscape characteristics and historical processes, including Pleistocene range dynamics and postglacial expansion (Krovi et al., 2025). Genetic responses to contemporary forest management should therefore be considered against species-specific ecological and historical backgrounds.

Our previous genetic analysis of 12 saproxylic beetle species demonstrated this complexity using genome-wide SNPs derived from double-digest restriction-site associated DNA sequencing (ddRADseq). Geographic location and associated historical population structure were the predominant determinants of genomic differentiation across Polish forests, whereas forest management had a secondary and species-dependent influence (Amer et al., 2026). Related SNP-based investigations using this genetic dataset have also examined the roles of structural heterogeneity, species-specific dispersal and ecological characteristics in shaping genetic diversity across saproxylic beetles (Krovi et al., 2026a,b). These results showed that contemporary habitat modification acts upon genetic variation already structured by spatial and historical processes. However, reducing sequence data to individual Single Nucleotide Polymorphism (SNPs) does not necessarily retain all genetic information contained within the original RAD loci, particularly when several linked polymorphisms occur within the same sequenced fragment.

Here, we use ddRADseq data from 12 saproxylic beetle species sampled across managed and primeval forests in Poland, retaining linked polymorphisms within RAD loci as microhaplotypes. The present study builds on previously generated ddRADseq data (Amer et al., 2026; Krovi et al., 2026a,b) by introducing a complementary microhaplotype-based approach, providing an additional perspective on genetic variation and enabling the investigation of questions that differ from those addressed in the previous SNP-based analyses. We first compared observed microhaplotype diversity with the diversity expected under linkage equilibrium, assessing how their difference varied with the number of polymorphic SNP positions per RAD locus. We then compare patterns of population differentiation estimated from SNPs and microhaplotypes, examine associations between individual microhaplotype homozygosity and site protection status, investigate relationships between site-level microhaplotype diversity and forest characteristics, and explore whether management-associated differences in homozygosity vary with species traits.

We evaluated five hypotheses. **H1:** Observed microhaplotype diversity at multi-SNP RAD loci is lower than the expected diversity under linkage equilibrium, reflecting non-random associations among polymorphic SNP positions within RAD loci. The magnitude of this deviation varies among species and with the number of polymorphic SNPs per locus. **H2:** SNP-based and microhaplotype-based estimates of population differentiation are expected to display broadly concordant patterns among sampling sites, although the strength of correspondence may vary among species because the two representations capture genetic variation differently. We evaluate concordance in relative differentiation patterns rather than assuming that the numerical values of the two statistics are directly equivalent. **H3:** individual microhaplotype homozygosity (F_HAP_) differs between protected and unprotected sites, as well as between primeval and managed sites with the magnitude and direction of this difference varying among species. Because protection status and forest-management regime are associated with other aspects of forest history and geographic location, these comparisons were interpreted as associations with site conditions rather than as isolated causal effects of protection or management. **H4:** Site-level microhaplotype diversity is associated with forest-management regime, time since the most recent logging event, and structural habitat characteristics such as deadwood diversity, total volume of deadwood, total volume of living trees, tree diversity, and abundance of veteran trees, with the strength and direction of these relationships differing among species. Because management regimes and habitat characteristics may be unevenly distributed across geographic regions, these analyses account for geographic structure, where supported by the sampling design and use sensitivity analyses to assess the robustness of management-related patterns. **H5:** Management-associated changes in microhaplotype homozygosity vary among species according to ecological and conservation-related species traits, including conservation status (protected/unprotected), occurrence (common/rare), trophy (fungivorous/invertivorous/xylovorous) and dispersal ability (high/moderate). Because these comparisons use only 12 species and the species-level traits may not be independent, H5 is treated as an exploratory analysis intended to identify patterns for future investigation rather than to establish general trait-based mechanisms.

## Materials and Methods

### Study sites

Sampling was conducted across eight forest complexes distributed throughout Poland, encompassing lowland, upland and mountainous regions and covering a broad geographic range of the focal saproxylic beetle species (Figure 1; Table S1). Sampling targeted forests with contrasting management histories and intensities, with the aim of capturing populations exposed to varying environmental conditions, particularly differences in forest naturalness and deadwood microhabitat availability. The sampled landscapes included relatively natural and protected forests, including Białowieża Forest (BF), the Holy Cross Mountains (HM) and the Carpathians (CM), as well as predominantly commercial forest complexes, including Augustów (AF), Knyszyn (KF), Silesian (SF), Oder (OF) and Barycz forests (DB) (Figure 1). Within these landscapes, 412 sampling sites represented variation in forest management and protection status, ranging from strictly protected or old-growth stands to actively managed stands.

**Figure 1.**
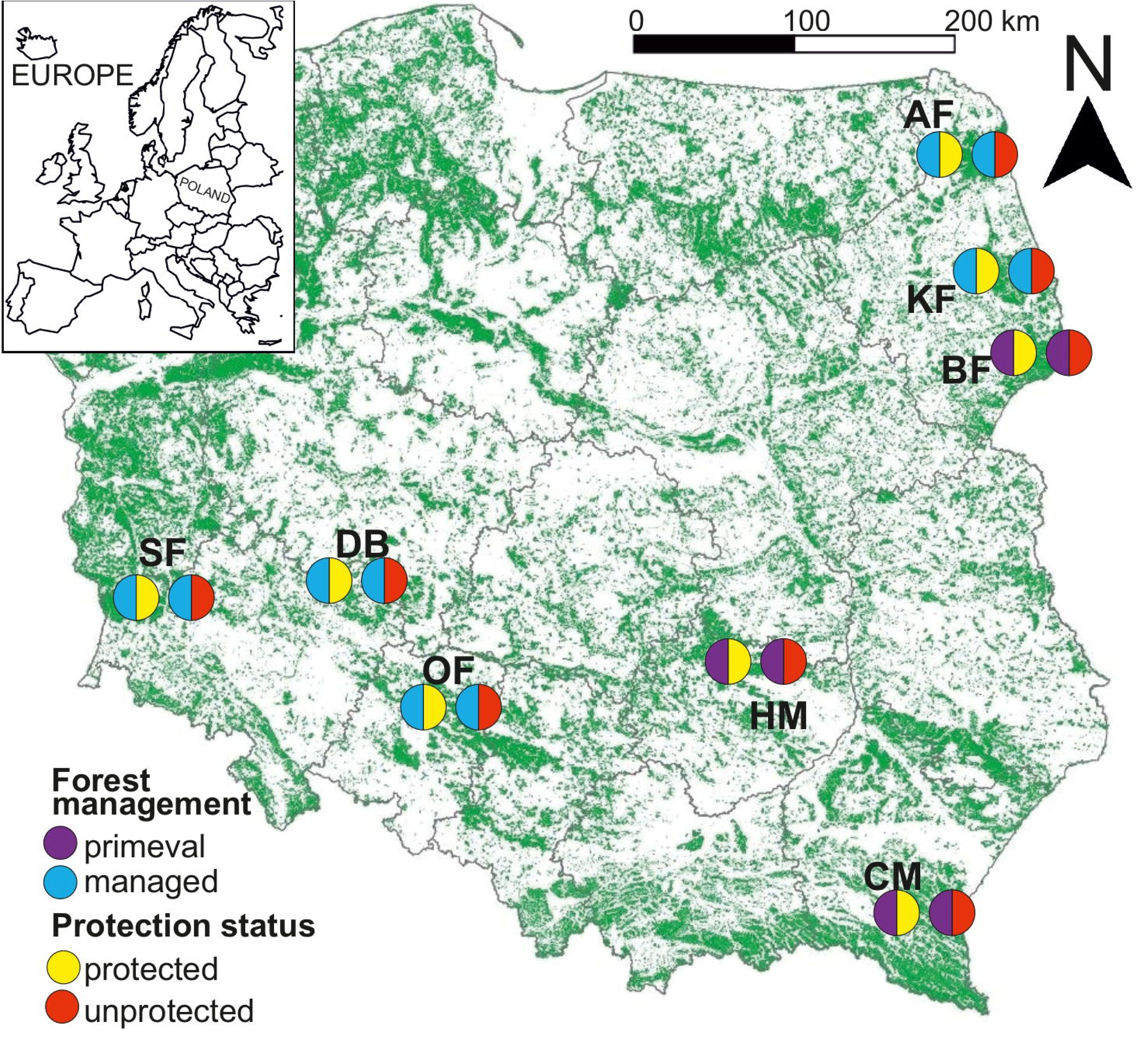
Geographic distribution of the eight studied forest areas in Poland. Representative sampling sites are illustrated using split-circle symbols. The left half indicates forest-management category (primeval, purple; managed, blue), while the right half indicates protection status (protected, yellow; unprotected, red). BF – Białowieża Forest, KF – Knyszyn Forest, AF – Augustów Forest, HM – Holy Cross Mts, CM – Carpathian Mts, OF – Oder Forest, DB – Barych Forest, SF – Silesia Forest. For clarity, only representative sites are shown; the complete set of sampling sites is provided in Table S1.

For the present study, sampling sites were classified according to the site-management variables relevant to the microhaplotype analyses (Table S1). In commercial forests, beetles were sampled in two types of sites: typical logged stands, generally characterized by limited deadwood resources, and nature reserves comprising old-growth forest patches that resemble natural forests but are limited in area and isolated within a matrix of commercial woodland. In primeval forests, sampling was conducted in strictly protected stands, such as national parks and nature reserves, and in stands without formal protection, where logging intensity is substantially lower than in commercial forests.

For the analyses, we considered forest-management category and protection status as two separate classification systems. For comparisons of forest-management categories, all sites within commercial forests were grouped as “managed”, whereas all sites within primeval forests were grouped as “primeval”. For comparisons of protection status, nature reserves within commercial forests and strictly protected sites within primeval forests were classified as “protected”, whereas logged sites within commercial forests and unprotected sites within primeval forests were classified as “unprotected”. Thus, the two classifications represent different aspects of the sampling context and should not be considered interchangeable (Figure 1).

Geographic coordinates were recorded for all 412 sampling localities and used to characterize the spatial distribution of populations and link genetic data with site-level environmental and management information (Table S1). Environmental characteristics were assessed within a 500 m² circular plot (radius 12.6 m) established at each sampling site. Six variables were considered: time (years) since the most recent logging or protection designation (Latest_logging), deadwood diversity (Dw_diversity), total deadwood volume (V_total), total volume of living trees (Wood_volume), tree diversity (Tree_diversity), and the abundance of veteran trees (veterans). Time since the most recent logging or intensive management activity was expressed based on the Polish Forest Data Bank (BDL; https://www.bdl.lasy.gov.pl/portal). While for protected sites, time since the most recent logging was calculated as the number of years since the designation of protection, using the relevant designation dates. Total deadwood volume was calculated from the volume of coarse deadwood recorded within each plot, excluding twigs and roots. Deadwood diversity was defined as the number of tree species represented among the deadwood. Total volume of living trees and tree diversity (tree species richness) were extracted from BDL forest inventory records. Veteran trees were defined as living monumental trees with advanced microhabitat features, including cavities, fungal fruiting bodies and large dead branches. They were identified using diameter-at-breast-height thresholds of >100 cm for oak and beech and >80 cm for other tree species (Table S1). Deadwood volume (Dw_diversity) and living wood volume (Wood volume) are reported in m^3^/ha; veteran-tree abundance (veterans) and number of tree species (Tree diversity) are per 500-m² plot; time since logging (Latest Logging) are in years. Because sampling sites differed in geographic location and forest-management history, management categories were not fully independent of geographic region. In particular, primeval and managed sites were unevenly distributed among regions (Figure 1). Accordingly, management-related associations were interpreted in the context of the spatial sampling design rather than as isolated causal effects of forest management.

### Beetle sampling

Beetles were sampled in 2023 and 2024 (c. 90% in 2023) using standardized complementary collection approaches tailored to their ecology. These included direct inspection of microhabitats beneath the bark of dying and dead trees, examination of decaying wood and wood-associated fungi, and funnel or barrier traps equipped with pheromone lures for selected taxa (see additional sampling details in Amer et al. 2026). Where available, approximately five individuals per species were sampled at each site for genetic analyses (Table S1).

Twelve saproxylic beetle species were selected for genetic analyses based on their ecological characteristics, conservation relevance and availability of sufficient samples across multiple localities: *Bolitophagus reticulatus* (Linnaeus, 1767) (Tenebrionidae) (Br, n=185), *Boros schneideri* (Panzer, 1796) (Boridae) (Bs, n=154), *Cucujus cinnaberinus* (Scopoli, 1763) (Cucujidae) (Cc, n=190), *Elater ferrugineus* (Linnaeus, 1758) (Elateridae) (Ef, n=84), *Ips acuminatus* (Gyllenhal, 1827) (Curculionidae) (Ia, n=125), *Ips typographus* (Linnaeus, 1758) (It, n=201), *Neomida haemorrhoidalis* (Fabricius, 1787) (Tenebrionidae) (Nh, n=199), *Phaenops cyanea* (Fabricius, 1775) (Buprestidae) (Pc, n=140), *Rhagium inquisitor* (Linnaeus, 1758) (Cerambycidae) (Ri, n=164), *Stenurella melanura* (Linnaeus, 1758) (Cerambycidae) (Sm, n=197), *Schizotus pectinicornis* (Linnaeus, 1758) (Pyrochroidae) (Sp, n=164), and *Thanasimus formicarius* (Linnaeus, 1758) (Cleridae) (Tf, n=178). Only one individual per species was selected from each deadwood log, reducing the likelihood of including siblings collected from the same log. Together, these species represent contrasting ecological characteristics, including differences in distribution, conservation status (*Bs*, *Cc*, and *Ef* are protected species, additionally *Nh* is red-listed, but not protected), trophy and dispersal abilities (Table S2).

Specimens were preserved in 95-99% ethanol and stored at -20°C until DNA extraction. Species identification was performed by experienced coleopterologists within the research team.

### DNA extraction, ddRAD library preparation and sequencing

DNA was extracted separately from each specimen using the Sherlock AX kit (A&A Biotechnology) following the manufacturer’s recommendations. Prior to extraction, specimens were rinsed with distilled water to minimize potential external contamination. The amount of tissue used depended on specimen size: whole individuals were used for small-bodied species, whereas selected body parts were used for larger beetles and larvae (Table 1). For *E. ferrugineus*, a single leg was sampled from adult specimens in accordance with conservation permit requirements and individuals were subsequently released.

**Table 1.** Sampling and RAD-sequencing summary for the 12 saproxylic beetle species included in the study. Number of sampled individuals (N Ind.); number of sampling sites (N Sites); number of retained RAD loci (N loci); total number of unique haplotypes (N Haps); total number of polymorphic nucleotide positions across retained RAD loci (N SNPs). Tissue codes indicate head and thorax (*), leg (**), or whole body (no asterisk). For a full list of geographic coordinates and regional classifications for each site, see Table S1 in the Supporting Information.

| **Species** | **Collection<br>stage** | **N<br>Ind.** | **N<br>Sites** | **N<br>loci** | **N<br>Haps** | **N<br>SNPs** |
|---|---|---|---|---|---|---|
| *Neomida haemorrhoidalis* | adults | 199 | 40 | 5304 | 38005 | 12467 |
| *Boros schneideri* | larvae* | 154 | 31 | 3004 | 23792 | 7736 |
| *Cucujus cinnaberinus* | larvae* | 190 | 39 | 1872 | 11523 | 3645 |
| *Elater ferrugineus* | adults** | 84 | 21 | 5514 | 27440 | 13575 |
| *Bolitophagus reticulatus* | adults | 185 | 38 | 1457 | 9615 | 2941 |
| *Stenurella melanura* | adults | 197 | 42 | 3229 | 38005 | 13708 |
| *Schizotus pectinicornis* | larvae | 164 | 35 | 1306 | 9397 | 3142 |
| *Thanasimus formicarius* | adults* | 178 | 37 | 521 | 7982 | 3008 |
| *Ips acuminatus* | adults | 125 | 26 | 1703 | 12911 | 4625 |
| *Ips typographus* | adults | 201 | 41 | 688 | 6676 | 1851 |
| *Rhagium inquisitor* | larvae* | 164 | 34 | 988 | 9982 | 3469 |
| *Phaenops cyanea* | adults* | 140 | 28 | 10771 | 81464 | 31781 |

Double-digest restriction-site-associated DNA sequencing (ddRADseq) libraries were prepared following Bayona-Vásquez et al. (2019). DNA concentrations were standardized to 20 ng/μL prior to library preparation. Individual samples were identified using iTru fusion primers with internal tags(iTru_NheI_R1_stub_A – H and iTru_EcoRI_R2_RC_stub_1-12), followed by restriction digestion, adapter ligation and pooling. Libraries were purified using AMPure XP magnetic beads (Beckman Coulter), amplified using indexed i5 and i7 primers, and quantified using Qubit. Fragment-size distributions were assessed using TapeStation, and fragments of approximately 400-600 bp were size-selected using Pippin Prep (Sage Science).

Libraries were normalized, pooled and sequenced by Admera Health (New Jersey, USA) using an Illumina NovaSeq X Plus platform with paired-end 150-bp sequencing, generating approximately four million reads per individual.

### Sequence processing and SNP dataset

Raw sequencing reads were initially assessed using FastQC v0.11.9 (Andrews 2010), with summary reports generated using MultiQC (Ewels et al. 2016). Reads were demultiplexed and quality filtered using process_radtags in Stacks v2.68 (Rochette et al. 2019). Reads containing low-quality bases, uncalled bases or incomplete restriction-site overhangs were excluded. RAD loci were assembled de novo separately for each species using the Stacks pipeline, with paired-end assembly enabled (denovo_map.pl --paired) and a maximum mismatch distance of two nucleotides between stacks within individuals (-M 2). During catalogue construction, a maximum mismatch distance of two nucleotides between loci from different individuals was used (-n 2). SNP genotypes were initially called using the Stacks populations module. Loci were retained if they were present in at least two populations (-p 2) and in at least 80% of individuals within each population (-r 0.80), with a minimum genotype depth of five reads (--min-gt-depth 5). The repeat-removal algorithm implemented in ustacks was also enabled to identify and remove unusually high-coverage repetitive stacks.

The resulting SNP dataset was exported in PLINK format and subjected to additional filtering. First, individuals with more than 50% missing genotypes were removed (--mind 0.50). Subsequently, loci with more than 30% missing genotypes (--geno 0.30) or a minor allele frequency below 0.03 (--maf 0.03) were excluded. A whitelist containing the retained SNP positions and their corresponding RAD-locus identifiers was then generated from the filtered PLINK map file. This whitelist was applied in a second Stacks populations run to retain the corresponding RAD loci for the microhaplotype analyses. The population map was updated to include only individuals retained after the PLINK sample-filtering step. Consequently, the final SNP and microhaplotype datasets were based on the same set of individuals and selected RAD loci.

### Microhaplotype diversity and linkage-equilibrium expectations (H1)

Phased SNP combinations within individual RAD loci were obtained from haplotype outputs generated by the Stacks *populations* module (Rochette et al., 2019). These combinations were treated as multi-allelic microhaplotypes, retaining the information contained in linked polymorphic SNP positions within each RAD locus. RAD loci containing a single polymorphic SNP were treated as single-SNP loci, whereas loci containing two or more polymorphic SNP positions were considered multi-SNP microhaplotypes. The number of polymorphic positions within each RAD locus was determined from the aligned haplotype sequences.

To assess how observed microhaplotype diversity compared with the diversity expected under independence among constituent SNP positions, observed haplotype diversity was compared with its analytical expectation under linkage equilibrium. Calculations were performed separately for each sampling population to avoid pooling individuals from different sites, which could introduce associations among SNPs resulting from population structure.

For each multi-SNP RAD locus, observed microhaplotype diversity (H_d_) was calculated from the frequencies of the observed haplotypes. Expected haplotype diversity under linkage equilibrium (H_d-LE_) was calculated analytically from the marginal allele frequencies at individual SNP positions, assuming independence among SNPs. Under this assumption, the expected frequency of a multi-SNP haplotype is the product of the frequencies of its constituent alleles across SNP positions. Expected diversity was consequently calculated as one minus the sum of squared expected haplotype frequencies (H_d-LE_ = 1 - ∑ (p-h,_LE_)^2^ ); where p-h,_LE_ is the expected frequency of haplotype h under linkage equilibrium.

The deviation from the linkage equilibrium expectation was quantified as ΔH_LE_ = H_d_ – H_d-LE_. Calculations were conducted within populations, and population-level results were subsequently summarised for each species. Differences were also examined in relation to the number of polymorphic SNP positions per RAD locus to investigate whether deviations from the linkage equilibrium expectation varied with locus-level polymorphism. Data processing, manipulation and summarisation were conducted using the tidyverse collection of packages (Wickham et al., 2019).

### Comparison of SNP- and microhaplotype-based genetic differentiation (H2)

To determine whether microhaplotypes recovered patterns of population differentiation consistent with those obtained from individual SNPs, genetic differentiation among sampling sites was quantified separately from the SNP and microhaplotype datasets.

Pairwise SNP-based F_ST_ and microhaplotype-based (Φ_ST_) matrices were generated independently for each species, following the general frameworks of Weir and Cockerham (1984) and Excoffier et al. (1992), respectively. SNP-based differentiation was obtained from the Stacks SNP F-statistic (F_ST_) outputs, whereas haplotype-based differentiation was obtained from the corresponding Stacks haplotype F-statistic phistats outputs. Sampling-site identifiers were matched between matrices, and comparisons were restricted to sites represented in both datasets. Concordance between the two measures of genetic differentiation was assessed using Mantel tests based on Spearman’s rank correlation, with 9,999 permutations (Mantel, 1967), implemented in the vegan package (Oksanen et al., 2026). For each species, the test compared the corresponding entries of the SNP-based and microhaplotype-based pairwise differentiation matrices. This rank-based approach evaluated whether pairs of sampling sites exhibiting relatively high or low differentiation based on SNPs showed corresponding patterns when differentiation was estimated from multi-allelic microhaplotypes. Spearman rank correlations were used because the objective was to evaluate correspondence in the relative differentiation among sampling sites without assuming a linear relationship or identical numerical scaling between F_ST_ and Φ_ST_.

Mantel correlation coefficients (ρ) were used to quantify the strength of correspondence between SNP- and microhaplotype-based differentiation.

The results are interpreted as concordance in the relative patterns of population differentiation, rather than evidence that F_ST_ and Φ_ST_ are numerically equivalent or that either representation provides universally superior estimates. Because pairwise differentiation values share sampling sites and are therefore not independent, the Mantel test was used to assess matrix-level correspondence rather than treating individual site pairs as independent replicates.

### Protection status and individual microhaplotype homozygosity (H3)

For each individual, microhaplotype homozygosity (F_HAP_) was calculated as the proportion of successfully scored polymorphic microhaplotype loci at which the two haplotype alleles were identical. F_HAP_ was interpreted as a measure of individual multi-locus haplotype homozygosity and not as a direct estimator of inbreeding, autozygosity or inbreeding load.

Differences in microhaplotype homozygosity between protected and unprotected sites were evaluated separately for each species using generalized linear mixed models with a beta-binomial error distribution. Rather than modelling the F_HAP_ proportion directly, the response was specified as the paired counts of homozygous and heterozygous successfully scored loci, thereby accounting for variation among individuals in the total number of genotyped loci. Site condition (protected vs unprotected) and geography (NE, SE, and SW) were included as fixed effects, and sampling site was included as a random intercept. The model structure was:

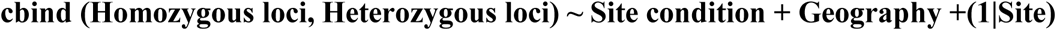

Models were fitted using the glmmTMB package in R (Brooks et al., 2017). For descriptive comparisons, site-level mean F_HAP_ values were compared between protected and unprotected sites within each species. These descriptive summaries were used to visualize differences in homozygosity between site conditions, whereas statistical inference was based on the species-specific GLMMs. Although both protected and unprotected sites were represented in all three geographic regions, their relative representation varied among regions. Therefore, a sensitivity analysis excluding geographic adjustment was conducted to assess the robustness of the estimated associations.

As a supplementary analysis, F_HAP_ was also compared between managed and primeval sites separately for each species, using the same general beta-binomial mixed-model framework. This comparison represented a broad contrast in forest context and management history rather than a direct comparison of logged and unlogged sites. Because primeval and managed sites were not represented across all geographic regions (primeval sites occurred only in NE and SE, whereas managed sites occurred only in NE and SW), sensitivity analyses were conducted both without geographic adjustment and, additionally, using only NE sites, where both management categories were represented for most species. Together, these analyses assessed the robustness of the observed patterns to geographic adjustment and geographic restriction.

For the protection status comparison, model estimates were calculated using the contrast (F_HAP_ = unprotected – protected). For the supplementary forest management comparison, descriptive differences in mean F_HAP_ were calculated as (F_HAP_ = managed – primeval). Accordingly, positive and negative values represent the direction of each respective contrast. The estimated differences are interpreted as associations between site condition and multilocus haplotype homozygosity, rather than a direct evidence of management-induced changes in inbreeding or genetic health.

Differences in geographic location, forest history, marker availability and genotype completeness may also contribute to observed patterns.

### Forest management history, habitat structure and site-level microhaplotype diversity (H4)

Site-level mean microhaplotype diversity (H_d_) was derived from the Stacks haplotype statistics and linked to forest-management and structural habitat metadata. Differences in mean H_d_ between managed and primeval sites were evaluated separately for each species, with P-values adjusted across species using the Benjamini-Hochberg FDR procedure (Benjamini & Hochberg, 1995).

Associations between site-level H_d_ and site and habitat characteristics were examined separately for each species using linear regression models. Six environmental predictors were evaluated individually: deadwood diversity (Dw_diversity), time since the most recent logging event (Latest_Logging), total deadwood volume (V_total), number of tree species (Tree_diversity), living wood volume (Wood_volume), and the abundance of veteran trees (veterans) (Table S1). For each species and predictor, a separate model was fitted with site-level mean H_d_ as the response variable and the respective environmental characteristic as the explanatory variable, with geographic region included as an adjustment factor (Mean H_d_ ∼ predictor + Geography).

Thus, each model evaluated the association with one environmental predictor while accounting for geographic differences, rather than including all environmental predictors simultaneously. A sensitivity analysis restricted to NE sites where both management categories were represented for most species was used to examine whether the observed patterns were also evident within this region. Additionally, sensitivity analyses were conducted using corresponding models without geographic adjustment (Mean H_d_ ∼ predictor) to evaluate whether the estimated associations depended on accounting for geographic region. For each species-predictor combination, the estimated regression coefficient, standard error, test statistic, confidence interval, sample size, model R², and raw P-value were extracted. Benjamini-Hochberg FDR correction was performed separately for each environmental predictor across the species analysed. These analyses were treated as exploratory associations rather than tests of causal environmental effects, and their interpretation considered the limited number of sampling sites available for some species and the potential influence of geographic structure.

### Species traits and management-associated changes in microhaplotype homozygosity (H5)

To explore whether species-level ecological characteristics were associated with interspecific variation in management-associated microhaplotype homozygosity, the difference in F_HAP_ between managed and primeval sites (ΔF_HAP_) was related to conservation status, occurrence, trophy and dispersal ability.

Because both ΔF_HAP_ and the ecological traits were defined at the species level, analyses used the 12 species as observational units rather than treating individual beetles as independent replicates. Separate univariate linear models were fitted for each trait, with conservation status, occurrence, trophy and dispersal ability treated as categorical factors. Overall trait effects were evaluated using analysis of variance (ANOVA). Each model therefore assessed the association between one species-level trait and the corresponding interspecific variation in ΔF_HAP_, without simultaneously adjusting for the other traits. Highly parameterized multivariable models were avoided because of the limited number of species (N = 12). These analyses were considered exploratory because the small number of species limits statistical power, some trait categories may be represented by few species, and ecological traits may be correlated. The results were therefore interpreted in terms of estimated patterns and uncertainty rather than as definitive evidence that particular traits determine genomic responses to forest management.

### General statistical procedures

All statistical analyses were conducted in R v4.4.1. (R Development Core Team, 2024). All visualizations have been done using the ggplot2 package (Wickham 2016). Analyses were performed separately for the 12 study species where appropriate. Statistical tests and model structures specific to each hypothesis are described in the corresponding sections above. Where multiple species-level tests were conducted for the same hypothesis or predictor, P-values were adjusted using the Benjamini-Hochberg false discovery rate (FDR) procedure (Benjamini & Hochberg, 1995). Statistical significance was assessed at α = 0.05.

## Results

A total of 1,981 individuals representing 12 saproxylic beetle species were analysed, covering 412 sampling sites across the study area (Table 1; Table S1). The number of individuals per species ranged from 84 in *E. ferrugineus* to 201 in *I. typographus*, while the number of sampled sites ranged from 21 to 42. Following RAD-sequencing and data filtering, the number of retained RAD loci varied considerably among species, from 521 in *T. formicarius* to 10,771 in *P. cyanea*. Across the 12 species, 276,792 unique haplotypes and 101,948 polymorphic nucleotide positions were identified. The number of unique haplotypes ranged from 6,676 in *I. typographus* to 81,464 in *P. cyanea*, whereas the number of polymorphic positions ranged from 1,851 to 31,781, respectively (Table 1).

### Genetic diversity captured by multi-SNP microhaplotypes (H1)

Observed microhaplotype diversity was consistently lower than the analytical expectation under linkage equilibrium across all 12 saproxylic beetle species (Table S3; Figure 2). Mean observed H_d_ ranged from 0.388 in *C. cinnaberinus* to 0.456 in *S. melanura*, whereas mean expected H_d_ under linkage equilibrium ranged from 0.453 in *C. cinnaberinus* to 0.558 in *E. ferrugineus*.

**Figure 2.**
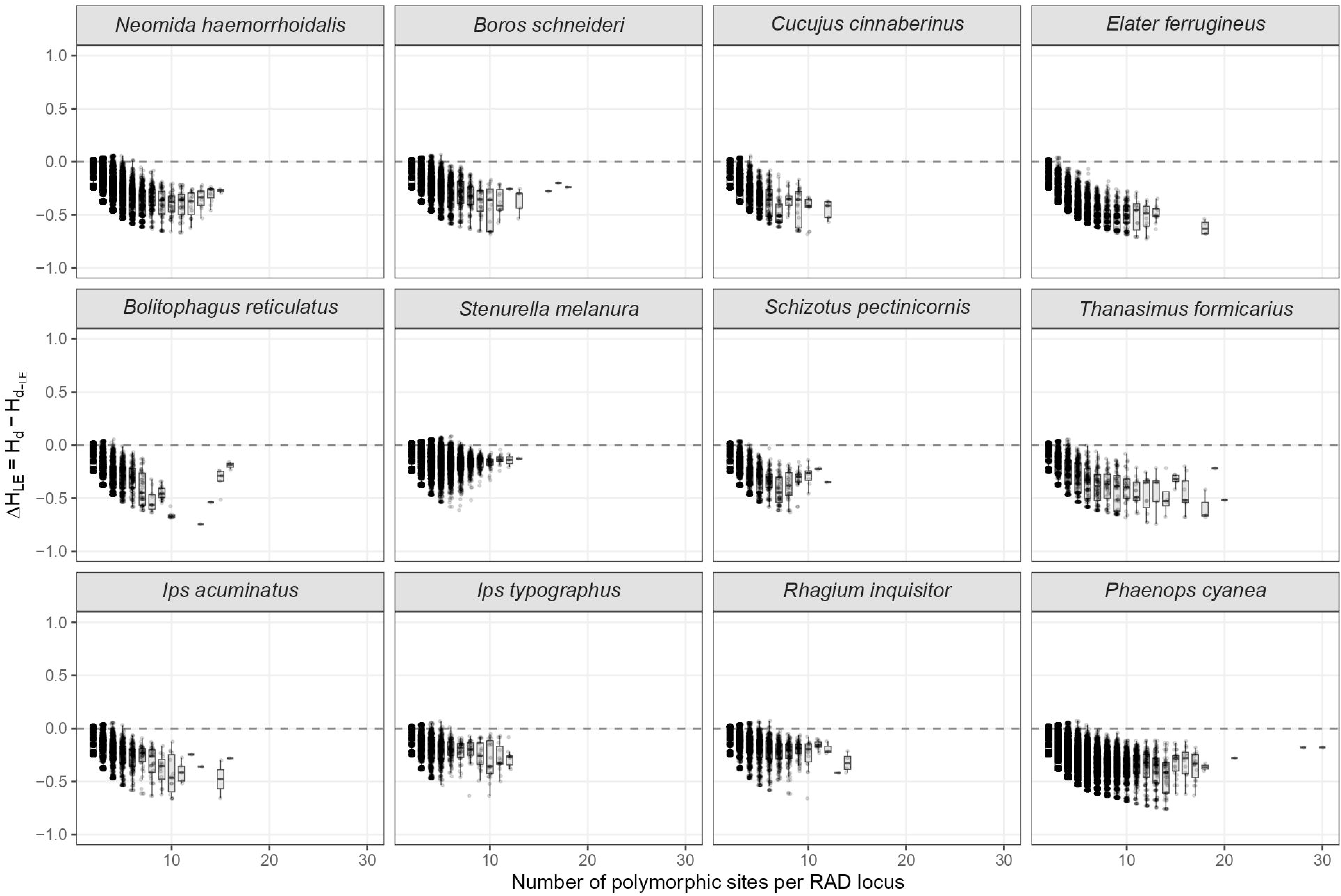
Distribution of differences between observed microhaplotype diversity and expected diversity under linkage equilibrium (ΔH_LE_ = H_d_ - H_d-LE_) across RAD loci containing two or more polymorphic SNP positions (≥ 2 SNPs) in 12 saproxylic beetle species. Panels represent individual species, and the x-axis indicates the number of polymorphic SNP positions per RAD locus. The horizontal dashed line at zero indicates equality between observed and expected diversity. Panels represent the 12 study species (*Br*: *Bolitophagus reticulatus*, *Bs*: *Boros schneideri*, *Cc*: *Cucujus cinnaberinus*, *Ef*: *Elater ferrugineus*, *Ia*: *Ips acuminatus*, *It*: *Ips typographus*, *Nh*: *Neomida haemorrhoidalis*, *Pc*: *Phaenops cyanea*, *Ri*: *Rhagium inquisitor*, *Sp*: *Schizotus pectinicornis*, *Sm*: *Stenurella melanura*, *Tf*: *Thanasimus formicarius*).

Accordingly, mean differences between observed and expected diversity (ΔH_d-LE_) were negative in all species, ranging from -0.060 in *N. haemorrhoidalis* to -0.155 in *E. ferrugineus*. The largest differences were observed in *E. ferrugineus* (-0.155), *P. cyanea* (-0.114) and *T. formicarius* (-0.096), whereas comparatively smaller differences were found in *N. haemorrhoidalis* (-0.060), *I. typographus* (-0.063) and *R. inquisitor* (-0.064) (Table S3; Figure 2).

### SNP- and microhaplotype-based differentiation (H2)

SNP-based F_ST_ and microhaplotype-based Φ_ST_ showed significant positive concordance in patterns of genetic differentiation among sampling sites across all 12 species (all FDR-adjusted P < 0.001; Figure 3). However, the strength of concordance varied markedly among species, with Mantel correlation coefficients ranging from ρ = 0.42 to 0.92. The strongest correspondence was observed in *I. acuminatus* and *S. pectinicornis* (ρ = 0.92), followed by *Br* and *Sm* (ρ = 0.87). Relatively strong concordance was also observed in *P. cyanea* (ρ = 0.78), *N. haemorrhoidalis* (ρ = 0.76), *T. formicarius* (ρ = 0.74), and *C. cinnaberinus* (ρ = 0.73). In contrast, weaker correspondence was observed in *E. ferrugineus* (ρ = 0.65), *I. typographus* (ρ = 0.59), *R. inquisitor* (ρ = 0.49), and particularly *B. schneideri* (ρ = 0.42) (Figure 3).

**Figure 3.**
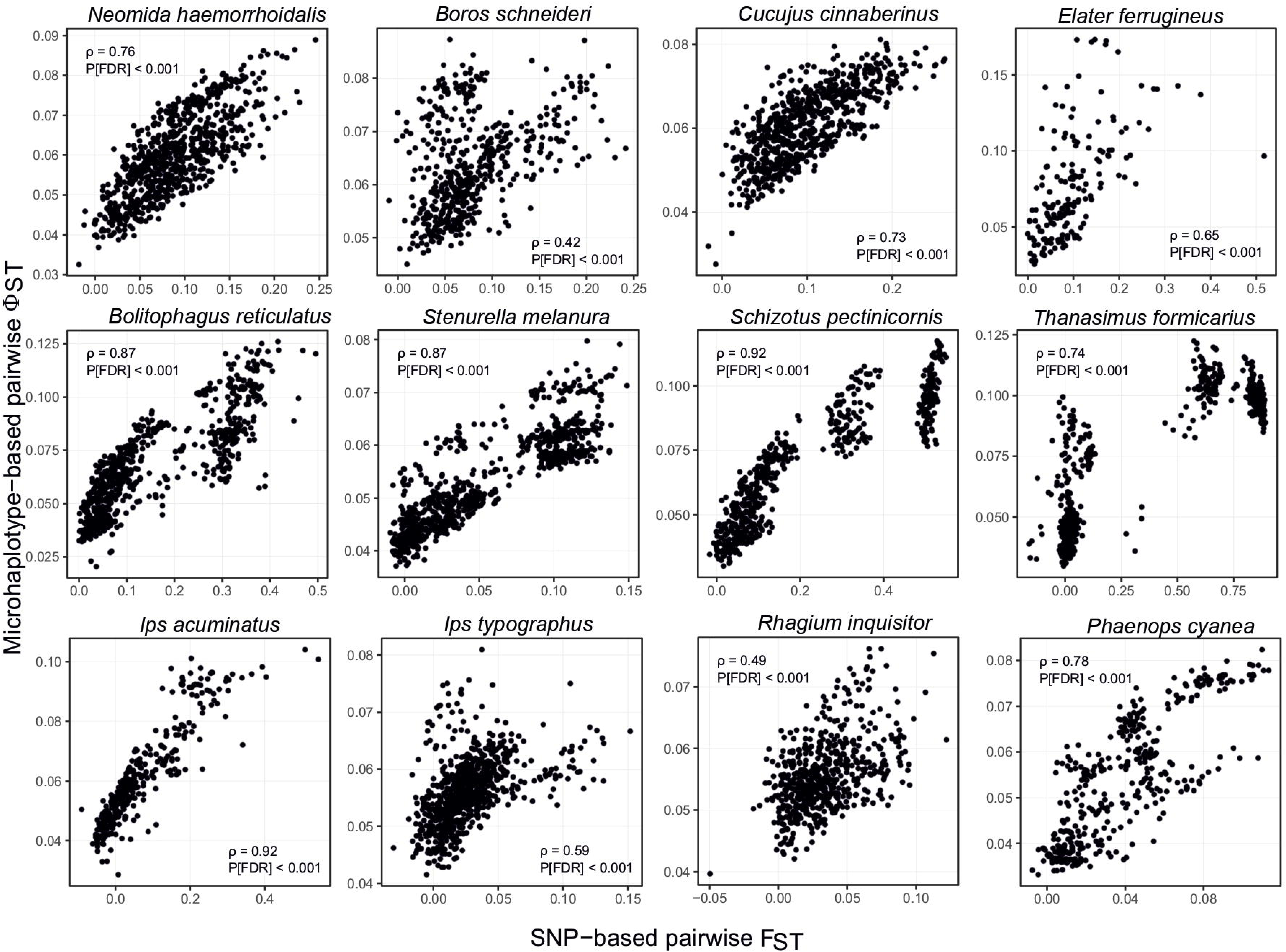
Concordance between SNP-based and microhaplotype-based estimates of pairwise genetic differentiation. Relationships between pairwise SNP-based (F_ST_) and microhaplotype-based (Φst) estimates among sampling sites for the 12 study species (*Br*: *Bolitophagus reticulatus*, *Bs*: *Boros schneideri*, *Cc*: *Cucujus cinnaberinus*, *Ef*: *Elater ferrugineus*, *Ia*: *Ips acuminatus*, *It*: *Ips typographus*, *Nh*: *Neomida haemorrhoidalis*, *Pc*: *Phaenops cyanea*, *Ri*: *Rhagium inquisitor*, *Sp*: *Schizotus pectinicornis*, *Sm*: *Stenurella melanura*, *Tf*: *Thanasimus formicarius*). Each point represents a pairwise comparison of sampling sites. Spearman Mantel correlation coefficients (ρ) and FDR-adjusted P-values are shown for each species. Positive correlations indicate concordance in the relative patterns of differentiation between marker representations.

### Site protection and management status with microhaplotype homozygosity (H3)

Species-specific models revealed variation among species in the association between site protection status and individual microhaplotype homozygosity (F_HAP_; Table 2 & Figure 4). Positive estimates indicated higher F_HAP_ in unprotected sites than in protected sites, whereas negative estimates indicated the opposite pattern. Nominal associations were detected in five species. *B. reticulatus* showed higher F_HAP_ in unprotected sites (estimate = 0.284, nominal *P* = 0.013) (Figure 4, Table 2). *C. cinnaberinus* showed the opposite pattern with lower F_HAP_ in unprotected than in protected sites (estimate = -0.120, nominal *P* = 0.008) (Figure 4, Table 2). Similarly, *R. inquisitor* showed higher F_HAP_ in unprotected sites (estimate = 0.051, nominal *P* = 0.028), while *B. schneideri* (estimate = -0.074, nominal *P* = 0.036) and *S. melanura* (estimate = -0.031, nominal *P* = 0.017) showed lower F_HAP_ in unprotected sites. However, none of these associations remained significant after correction for multiple testing (all FDR-adjusted *P* ≥ 0.070). No nominally significant association was detected in the remaining species (Table 2).

**Figure 4.**
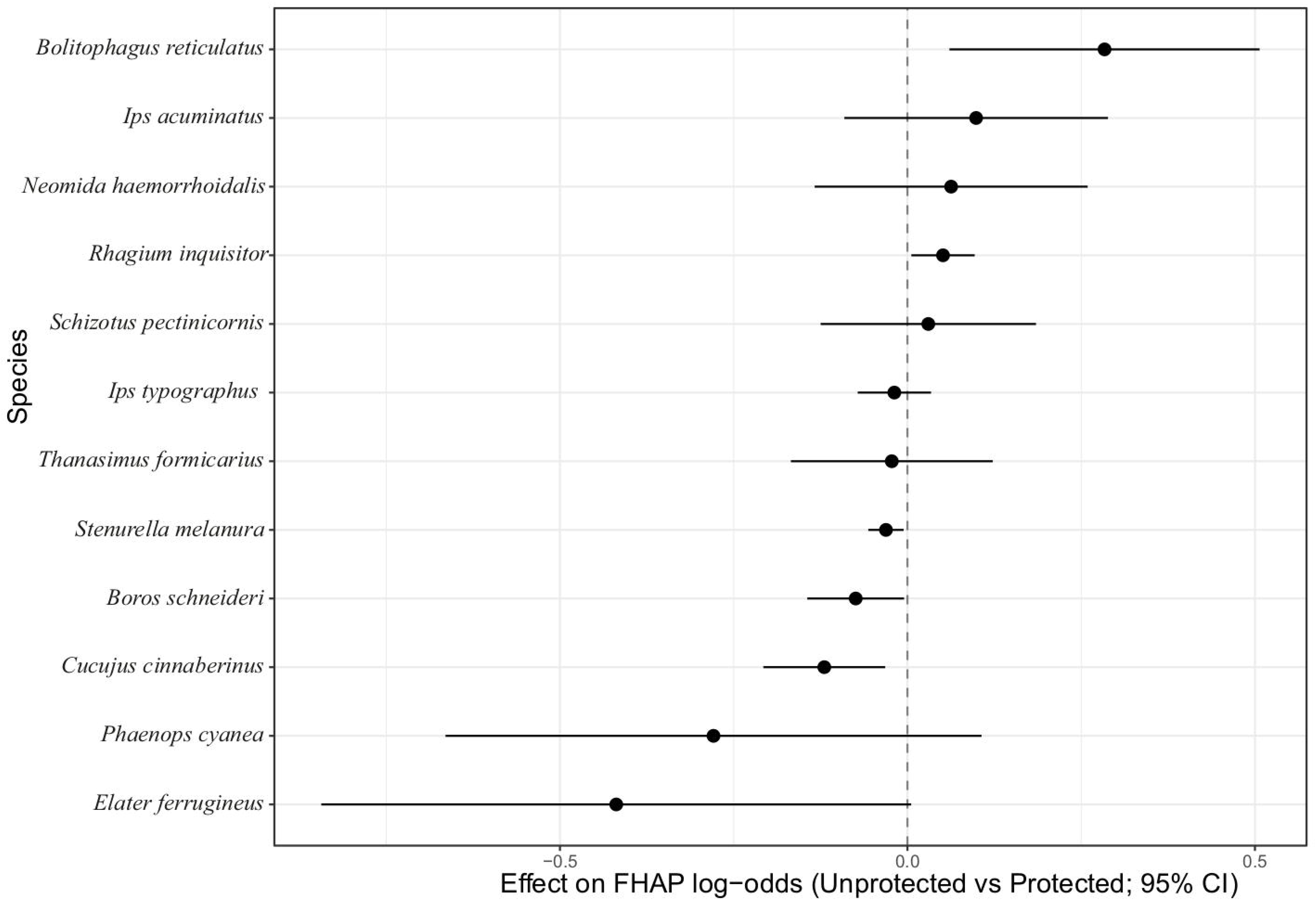
Species-specific associations between site protection status and individual microhaplotype homozygosity. Estimated effects of site protection status on individual microhaplotype homozygosity (F_HAP_) from species-specific beta-binomial mixed models adjusted for geographic region. Points indicate estimated effects on F_HAP_ log-odds for unprotected relative to protected sites, and horizontal lines indicate 95% confidence intervals. Positive estimates indicate higher F_HAP_ in unprotected sites, whereas negative estimates indicate higher F_HAP_ in protected sites. The vertical dashed line indicates no difference between site categories. The 12 study species are *Br*: *Bolitophagus reticulatus*, *Bs*: *Boros schneideri*, *Cc*: *Cucujus cinnaberinus*, *Ef*: *Elater ferrugineus*, *Ia*: *Ips acuminatus*, *It*: *Ips typographus*, *Nh*: *Neomida haemorrhoidalis*, *Pc*: *Phaenops cyanea*, *Ri*: *Rhagium inquisitor*, *Sp*: *Schizotus pectinicornis*, *Sm*: *Stenurella melanura*, *Tf*: *Thanasimus formicarius*.

**Table 2.** Species-specific effects of site protection status on individual microhaplotype homozygosity (F_HAP_), adjusted for geographic region. Estimates represent the contrast between unprotected and protected sites. The table reports effect estimates, standard errors (SE), test statistics, P-values, 95% confidence intervals, Akaike’s information criterion (AIC), and Benjamini-Hochberg false discovery rate-adjusted P-values (P_FDR).

| **Species** | **Estimate** | **SE** | **Statistic** | **P value** | **conf.low** | **conf.high** | **AIC** | **P_FDR** |
|---|---|---|---|---|---|---|---|---|
| *Neomida haemorrhoidalis* | 0.063 | 0.100 | 0.628 | 0.530 | -0.133 | 0.259 | 2301.97 | 0.636 |
| *Boros schneideri* | -0.074 | 0.036 | -2.093 | 0.036 | -0.144 | -0.005 | 1684.21 | 0.087 |
| *Cucujus cinnaberinus* | -0.120 | 0.045 | -2.673 | 0.008 | -0.207 | -0.032 | 2081.49 | 0.070 |
| *Elater ferrugineus* | -0.419 | 0.217 | -1.935 | 0.053 | -0.843 | 0.005 | 1031.56 | 0.106 |
| *Bolitophagus reticulatus* | 0.284 | 0.114 | 2.491 | 0.013 | 0.060 | 0.507 | 2220.25 | 0.070 |
| *Stenurella melanura* | -0.031 | 0.013 | -2.377 | 0.017 | -0.056 | -0.005 | 2158.08 | 0.070 |
| *Schizotus pectinicornis* | 0.030 | 0.079 | 0.380 | 0.704 | -0.125 | 0.185 | 1614.63 | 0.761 |
| *Thanasimus formicarius* | -0.022 | 0.074 | -0.304 | 0.761 | -0.168 | 0.123 | 1629.61 | 0.761 |
| *Ips acuminatus* | 0.099 | 0.097 | 1.022 | 0.307 | -0.091 | 0.288 | 1413.11 | 0.460 |
| *Ips typographus* | -0.019 | 0.027 | -0.698 | 0.485 | -0.072 | 0.034 | 1690.21 | 0.636 |
| *Rhagium inquisitor* | 0.051 | 0.023 | 2.202 | 0.028 | 0.006 | 0.097 | 1432.54 | 0.083 |
| *Phaenops cyanea* | -0.279 | 0.197 | -1.418 | 0.156 | -0.665 | 0.107 | 2151.89 | 0.268 |

Sensitivity analyses comparing protection-effect estimates from models with and without geographic adjustment showed broadly similar patterns, although the magnitude of some estimates differed (Figure S1).

In the supplementary management analysis, models showed variation in the direction and magnitude of management effects among species (Figure S2), however, no species showed a statistically significant management effect after correction for multiple testing (all FDR-adjusted *P* ≥ 0.49). The strongest nominal evidence was observed in *R. inquisitor*, for which F_HAP_ was estimated to be lower in managed than in primeval forests (nominal *P* = 0.045, FDR-adjusted *P* = 0.49). Sensitivity comparisons indicated that management-effect estimates could change in magnitude, and occasionally direction, depending on whether geographic region was included in the model (Figure S3). Because management types were unevenly distributed across regions (Table S4 & S5), these results should be interpreted cautiously and do not support a consistent association between forest management type and F_HAP_ across the 12 species. Also, we repeated the species-specific beta-binomial models of the association between forest management and F_HAP_ using only sites in the NE region, where both managed and primeval forests were represented. The analysis included 11 species, with 14-27 sampling sites per species due to a lack of *Ef* samples from managed sites in NE Poland (Table S4). No species showed a statistically significant management association after FDR correction (all FDR-adjusted P ≥ 0.448) (Table S6). *R. inquisitor* showed a nominal association (P = 0.041), but this did not remain significant after FDR correction (P_FDR = 0.448) (Table S6).

### Microhaplotype diversity and species-specific associations with forest characteristics (H4)

For each species, separate geography-adjusted linear regression models were fitted to evaluate associations between site-level microhaplotype diversity (H_d_) and each of six forest characteristics: time since last logging, deadwood diversity, total deadwood volume, total volume of living trees, tree diversity, and veterans. Although estimated effects varied in magnitude and direction among species and predictors (Figure 5), none remained statistically significant after Benjamini-Hochberg correction for multiple testing in the geography-adjusted models (all FDR-adjusted P > 0.112) (Table 3). Sensitivity analyses using models without geographic adjustment produced different results. Positive associations between deadwood diversity and H_d_ were detected in *B. reticulatus* (estimate = 0.050, nominal P = 0.004, FDR-adjusted P = 0.031) and *I. typographus* (estimate = 0.010, nominal P = 0.005, FDR-adjusted P = 0.031) (Table S7). Time since last logging was positively associated with H_d_ in *T. formicarius* (estimate = 0.008, nominal P = 0.003, FDR-adjusted P = 0.038) in the same sensitivity analysis (Table S7). However, these associations did not remain significant in the geography-adjusted primary models. This difference shows apparent relationships between microhaplotype diversity and forest characteristics that were sensitive to geographic adjustment (Table S7). Additionally, we compared H_d_ between managed and primeval forest sites using only sampling sites in the NE region, where both management categories were represented for 11 of the 12 species (Table S8). No species showed a statistically significant difference after FDR correction (all FDR-adjusted P ≥ 0.317) (Table S8). *P. cyanea* showed a nominal difference (P = 0.040), with higher mean H_d_ in managed than in primeval sites, but this association was not significant after correction (P_FDR = 0.317) (Table S8). *R. inquisitor* also showed a trend towards higher mean H_d_ in managed sites (P = 0.058, P_FDR = 0.317) (Table S8).

**Figure 5.**
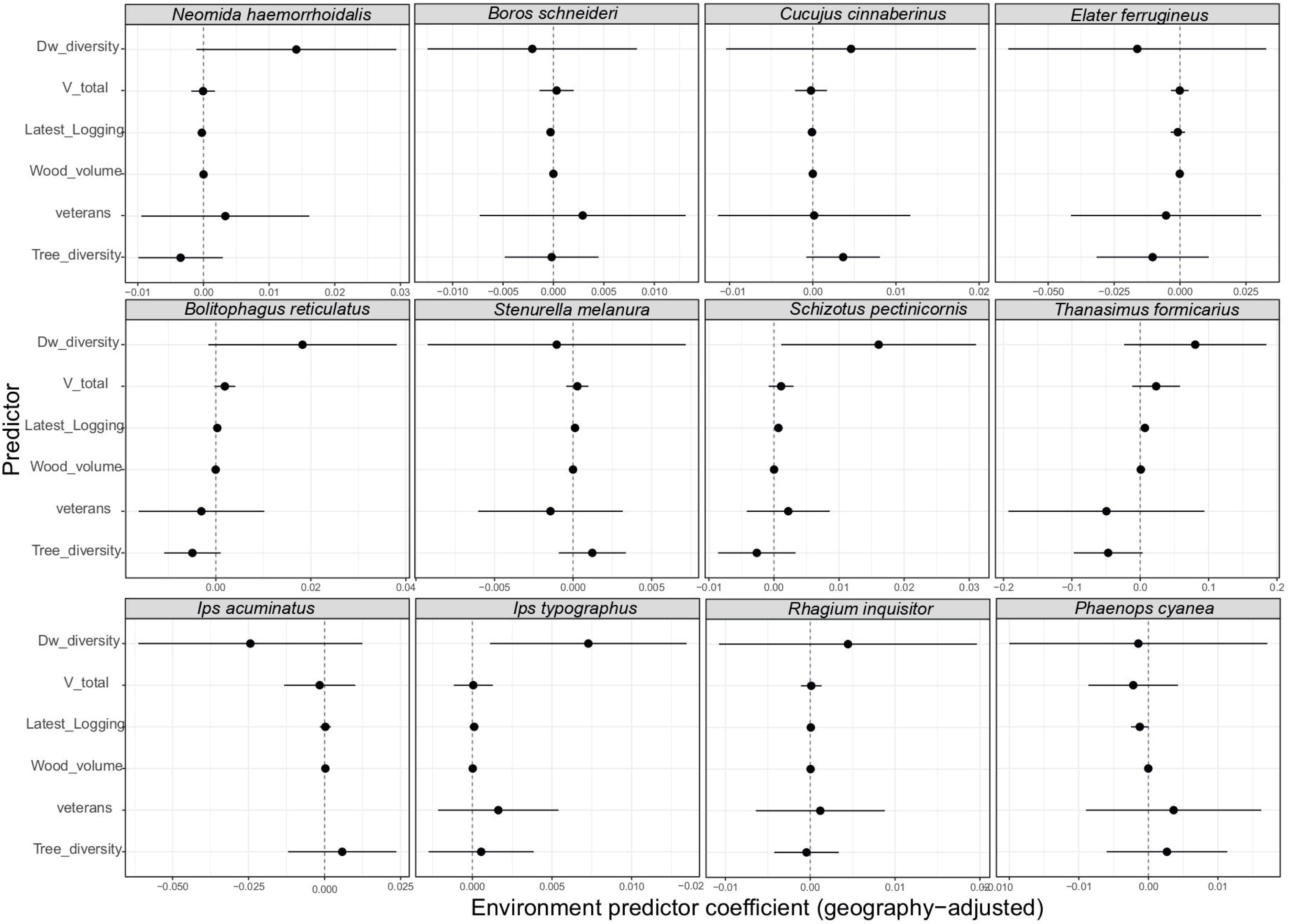
Species-specific associations between forest structural characteristics and site-level microhaplotype diversity. Estimated coefficients and 95% confidence intervals for associations between site-level microhaplotype diversity (H_d_) and six forest characteristics: deadwood diversity (Dw_diversity), deadwood total volume (V_total), time since the most recent logging event (Latest_logging), wood volume (Wood_volume), veteran-tree abundance (veterans), and tree diversity (Tree_diversity). Estimates are from species-specific models adjusted for geographic region. Panels represent the 12 study species, and the vertical dashed line indicates a coefficient of zero.

**Table 3.** Species-specific associations between forest characteristics and site-level microhaplotype diversity (H_d_) estimated using geography-adjusted linear regression models. Estimate represents the regression coefficient for the environmental predictor. R^2^ represents the proportion of variation explained by the full model, including the predictor and geographic region. FDR correction was performed separately for each environmental predictor across species. *I. acuminatus* has 23 sites with 0 veterans and 3 sites with only 1 veteran, so “veterans” was excluded.

| Species | Predictor | Estimate | SE | Statistic | P value | conf.low | conf.high | R2 | P_FDR |
|---|---|---|---|---|---|---|---|---|---|
| *Neomida haemorrhoidalis* | Latest_Logging | 0 | 0 | -0.787 | 0.437 | -0.001 | 0 | 0.103 | 0.656 |
|  | Dw_diversity | 0.014 | 0.007 | 1.897 | 0.067 | -0.001 | 0.029 | 0.182 | 0.209 |
|  | V_total | 0 | 0.001 | -0.059 | 0.953 | -0.002 | 0.002 | 0.084 | 0.953 |
|  | Tree_diversity | -0.003 | 0.003 | -1.105 | 0.278 | -0.01 | 0.003 | 0.12 | 0.609 |
|  | Wood_volume | 0 | 0 | 0.301 | 0.765 | 0 | 0 | 0.087 | 0.929 |
|  | veterans | 0.003 | 0.006 | 0.532 | 0.599 | -0.009 | 0.016 | 0.093 | 0.828 |
| *Boros schneideri* | Latest_Logging | 0 | 0 | -1.82 | 0.079 | -0.001 | 0 | 0.143 | 0.238 |
|  | Dw_diversity | -0.002 | 0.005 | -0.414 | 0.682 | -0.012 | 0.008 | 0.048 | 0.818 |
|  | V_total | 0 | 0.001 | 0.377 | 0.709 | -0.001 | 0.002 | 0.047 | 0.953 |
|  | Tree_diversity | 0 | 0.002 | -0.078 | 0.938 | -0.005 | 0.004 | 0.042 | 0.938 |
|  | Wood_volume | 0 | 0 | -0.09 | 0.929 | 0 | 0 | 0.042 | 0.929 |
|  | veterans | 0.003 | 0.005 | 0.582 | 0.565 | -0.007 | 0.013 | 0.053 | 0.828 |
| *Cucujus cinnaberinus* | Latest_Logging | 0 | 0 | -0.387 | 0.701 | -0.001 | 0 | 0.1 | 0.79 |
|  | Dw_diversity | 0.005 | 0.007 | 0.624 | 0.537 | -0.01 | 0.02 | 0.106 | 0.74 |
|  | V_total | 0 | 0.001 | -0.226 | 0.822 | -0.002 | 0.002 | 0.097 | 0.953 |
|  | Tree_diversity | 0.004 | 0.002 | sty.68 | 0.102 | -0.001 | 0.008 | 0.167 | 0.409 |
|  | Wood_volume | 0 | 0 | 0.175 | 0.862 | 0 | 0 | 0.097 | 0.929 |
|  | veterans | 0 | 0.006 | 0.03 | 0.976 | -0.011 | 0.012 | 0.096 | 0.976 |
| *Elater ferrugineus* | Latest_Logging | -0.001 | 0.001 | -0.679 | 0.511 | -0.004 | 0.002 | 0.151 | 0.681 |
|  | Dw_diversity | -0.016 | 0.022 | -0.73 | 0.481 | -0.065 | 0.033 | 0.156 | 0.74 |
|  | V_total | 0 | 0.002 | -0.07 | 0.945 | -0.003 | 0.003 | 0.115 | 0.953 |
|  | Tree_diversity | -0.01 | 0.01 | -1.077 | 0.305 | -0.032 | 0.011 | 0.199 | 0.609 |
|  | Wood_volume | 0 | 0 | -0.738 | 0.476 | 0 | 0 | 0.157 | 0.929 |
|  | veterans | -0.005 | 0.016 | -0.324 | 0.752 | -0.041 | 0.031 | 0.123 | 0.828 |
| *Bolitophagus reticulatus* | Latest_Logging | 0 | 0 | 0.821 | 0.418 | 0 | 0.001 | 0.751 | 0.656 |
|  | Dw_diversity | 0.018 | 0.01 | 1.872 | 0.07 | -0.002 | 0.038 | 0.77 | 0.209 |
|  | V_total | 0.002 | 0.001 | 1.748 | 0.09 | 0 | 0.004 | 0.767 | 0.953 |
|  | Tree_diversity | -0.005 | 0.003 | -1.686 | 0.101 | -0.011 | 0.001 | 0.766 | 0.409 |
|  | Wood_volume | 0 | 0 | -0.628 | 0.534 | 0 | 0 | 0.749 | 0.929 |
|  | veterans | -0.003 | 0.007 | -0.466 | 0.644 | -0.016 | 0.01 | 0.748 | 0.828 |
| *Stenurella melanura* | Latest_Logging | 0 | 0 | 1.227 | 0.227 | 0 | 0 | 0.75 | 0.545 |
|  | Dw_diversity | -0.001 | 0.004 | -0.256 | 0.799 | -0.009 | 0.007 | 0.74 | 0.872 |
|  | V_total | 0 | 0 | 0.786 | 0.437 | 0 | 0.001 | 0.744 | 0.953 |
|  | Tree_diversity | 0.001 | 0.001 | sty.16 | 0.253 | -0.001 | 0.003 | 0.749 | 0.609 |
|  | Wood_volume | 0 | 0 | -0.106 | 0.916 | 0 | 0 | 0.74 | 0.929 |
|  | veterans | -0.001 | 0.002 | -0.632 | 0.531 | -0.006 | 0.003 | 0.743 | 0.828 |
| *Schizotus pectinicornis* | Latest_Logging | 0.001 | 0 | 2.417 | 0.023 | 0 | 0.001 | 0.574 | 0.136 |
|  | Dw_diversity | 0.016 | 0.007 | 2.206 | 0.036 | 0.001 | 0.031 | 0.561 | 0.209 |
|  | V_total | 0.001 | 0.001 | 1.172 | 0.251 | -0.001 | 0.003 | 0.507 | 0.953 |
|  | Tree_diversity | -0.003 | 0.003 | -0.91 | 0.371 | -0.009 | 0.003 | 0.497 | 0.636 |
|  | Wood_volume | 0 | 0 | 0.402 | 0.691 | 0 | 0 | 0.485 | 0.929 |
|  | veterans | 0.002 | 0.003 | 0.707 | 0.486 | -0.004 | 0.009 | 0.491 | 0.828 |
| *Thanasimus formicarius* | Latest_Logging | 0.007 | 0.003 | 2.742 | 0.01 | 0.002 | 0.012 | 0.395 | 0.119 |
|  | Dw_diversity | 0.08 | 0.051 | 1.575 | 0.125 | -0.024 | 0.185 | 0.307 | 0.3 |
|  | V_total | 0.023 | 0.017 | 1.352 | 0.186 | -0.012 | 0.058 | 0.294 | 0.953 |
|  | Tree_diversity | -0.047 | 0.025 | -1.894 | 0.067 | -0.097 | 0.004 | 0.329 | 0.409 |
|  | Wood_volume | 0.001 | 0 | 2.433 | 0.021 | 0 | 0.002 | 0.37 | 0.124 |
|  | veterans | -0.049 | 0.07 | -0.702 | 0.488 | -0.193 | 0.094 | 0.265 | 0.828 |
| *Ips acuminatus* | Latest_Logging | 0 | 0.001 | 0.27 | 0.79 | -0.002 | 0.002 | 0.123 | 0.79 |
|  | Dw_diversity | -0.024 | 0.018 | -1.374 | 0.183 | -0.061 | 0.012 | 0.189 | 0.367 |
|  | V_total | -0.002 | 0.006 | -0.287 | 0.777 | -0.013 | 0.01 | 0.123 | 0.953 |
|  | Tree_diversity | 0.006 | 0.009 | 0.675 | 0.507 | -0.012 | 0.024 | 0.138 | 0.709 |
|  | Wood_volume | 0 | 0 | 2.851 | 0.009 | 0 | 0 | 0.357 | 0.112 |
| *Ips typographus* | Latest_Logging | 0 | 0 | 0.798 | 0.43 | 0 | 0 | 0.278 | 0.656 |
|  | Dw_diversity | 0.007 | 0.003 | lut.39 | 0.022 | 0.001 | 0.013 | 0.364 | 0.209 |
|  | V_total | 0 | 0.001 | 0.091 | 0.928 | -0.001 | 0.001 | 0.266 | 0.953 |
|  | Tree_diversity | 0.001 | 0.002 | 0.338 | 0.737 | -0.003 | 0.004 | 0.268 | 0.885 |
|  | Wood_volume | 0 | 0 | 0.885 | 0.382 | 0 | 0 | 0.281 | 0.929 |
|  | veterans | 0.002 | 0.002 | 0.871 | 0.389 | -0.002 | 0.005 | 0.281 | 0.828 |
| *Rhagium inquisitor* | Latest_Logging | 0 | 0 | 0.353 | 0.727 | 0 | 0 | 0.225 | 0.79 |
|  | Dw_diversity | 0.004 | 0.007 | 0.597 | 0.555 | -0.011 | 0.02 | 0.231 | 0.74 |
|  | V_total | 0 | 0.001 | 0.194 | 0.848 | -0.001 | 0.001 | 0.223 | 0.953 |
|  | Tree_diversity | 0 | 0.002 | -0.237 | 0.815 | -0.004 | 0.003 | 0.224 | 0.889 |
|  | Wood_volume | 0 | 0 | 1.426 | 0.164 | 0 | 0 | 0.272 | 0.657 |
|  | veterans | 0.001 | 0.004 | 0.318 | 0.753 | -0.006 | 0.009 | 0.225 | 0.828 |
| *Phaenops cyanea* | Latest_Logging | -0.001 | 0.001 | -2.051 | 0.051 | -0.002 | 0 | 0.188 | 0.206 |
|  | Dw_diversity | -0.001 | 0.009 | -0.161 | 0.874 | -0.02 | 0.017 | 0.047 | 0.874 |
|  | V_total | -0.002 | 0.003 | -0.696 | 0.493 | -0.009 | 0.004 | 0.065 | 0.953 |
|  | Tree_diversity | 0.003 | 0.004 | 0.635 | 0.531 | -0.006 | 0.011 | 0.061 | 0.709 |
|  | Wood_volume | 0 | 0 | -0.242 | 0.811 | 0 | 0 | 0.048 | 0.929 |
|  | veterans | 0.004 | 0.006 | 0.595 | 0.558 | -0.009 | 0.016 | 0.06 | 0.828 |

### Species traits and management-associated changes in microhaplotype homozygosity (H5)

Management-associated changes in microhaplotype homozygosity ΔF_HAP_ (managed - primeval) varied among the 12 species, with both positive and negative contrasts observed (Figure 6).

**Figure 6.**
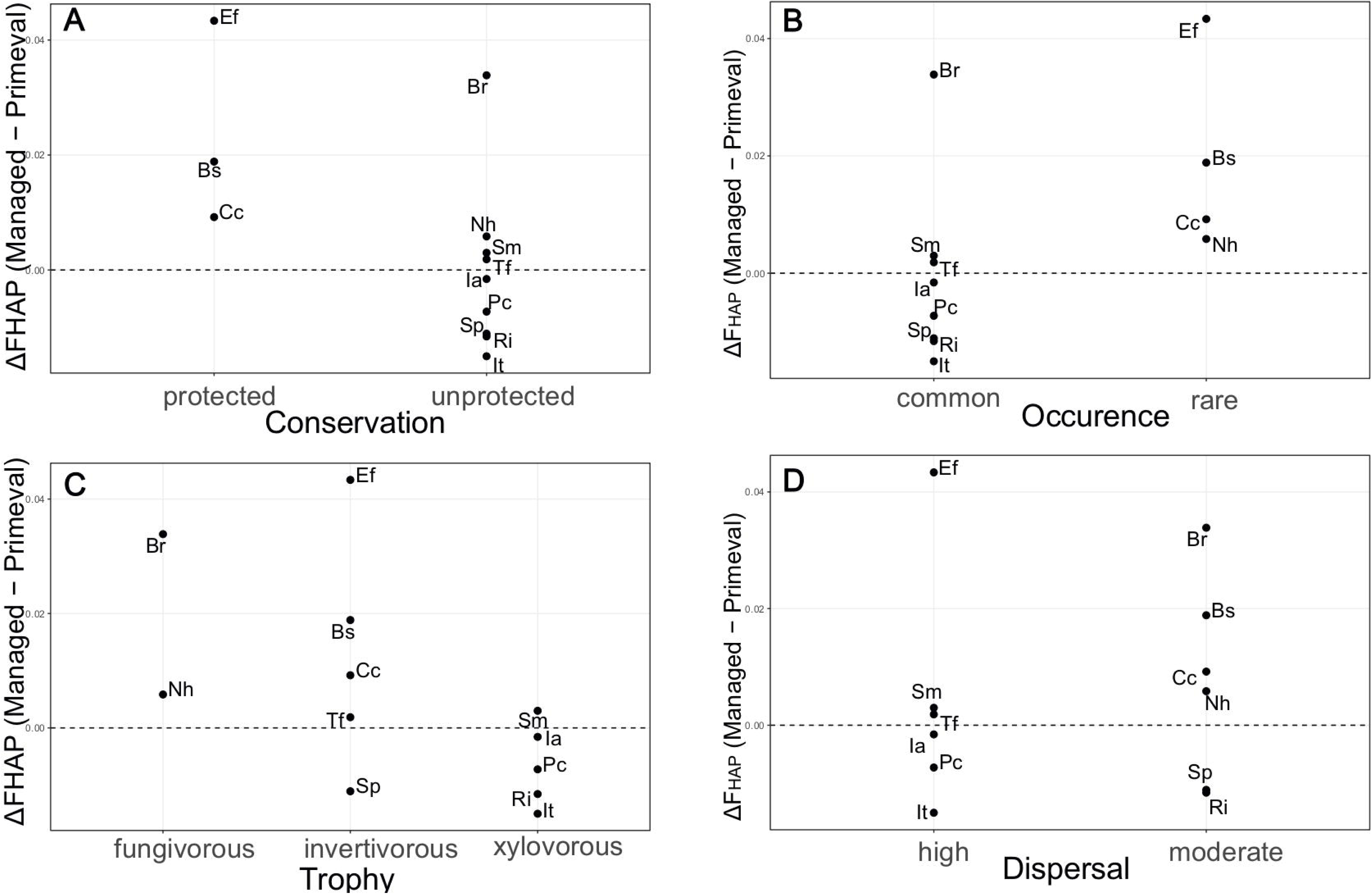
Relationships between species-level ecological traits and management-associated differences in microhaplotype homozygosity. Differences in microhaplotype homozygosity between managed and primeval sites (ΔF_HAP_) across 12 species (*Br*: *Bolitophagus reticulatus*, *Bs*: *Boros schneideri*, *Cc*: *Cucujus cinnaberinus*, *Ef*: *Elater ferrugineus*, *Ia*: *Ips acuminatus*, *It*: *Ips typographus*, *Nh*: *Neomida haemorrhoidalis*, *Pc*: *Phaenops cyanea*, *Ri*: *Rhagium inquisitor*, *Sp*: *Schizotus pectinicornis*, *Sm*: *Stenurella melanura*, *Tf*: *Thanasimus formicarius*), grouped by (A) conservation status, (B) occurrence status, (C) trophic group, and (D) dispersal category. Positive values indicate higher F_HAP_ in managed sites, whereas negative values indicate higher F_HAP_ in primeval sites. Each point represents one species.

Conservation status showed the strongest association with ΔF_HAP_ (F_1,10_ = 5.522, P = 0.041) (Figure 6A; Table 4). Protected species tended to show positive ΔF_HAP_, indicating greater microhaplotype homozygosity in managed than in primeval sites, whereas responses among unprotected species were generally smaller and included both positive and negative values (Figure 6A). Occurrence status showed a similar but weaker pattern (F_1,10_ = 4.301, P = 0.065) (Figure 6B; Table 4). Rare species tended to exhibit positive ΔF_HAP_, whereas responses among common species were predominantly small or negative (Figure 6B). Trophy also showed variation in management-associated homozygosity (F_2,9_ = 2.705, P = 0.120). Fungivores showed positive responses, xylovores were predominantly negative, and invertivores displayed greater interspecific variation (Figure 6C; Table 4). In contrast, dispersal category showed little evidence of an association with ΔF_HAP_ (F_1,10_ = 0.099, P = 0.759), with both moderate- and high-dispersal species exhibiting heterogeneous responses (Figure 6D; Table 4). None of these associations remained statistically significant after Benjamini-Hochberg correction (all FDR-adjusted P ≥ 0.120).

**Table 4.** Associations between species traits (Conservation, Occurrence, Trophy, and dispersal) and management-associated differences in microhaplotype homozygosity (ΔF_HAP_). ΔF_HAP_ was calculated for each species as the difference in mean microhaplotype homozygosity between managed and primeval forests (F_HAP,Managed_ - F_HAP,Primeval_).

| **Term** | **df** | **F value** | **P value** | **P_FDR** |
|---|---|---|---|---|
| **Conservation** | 1 | 5.522 | 0.041 | 0.120 |
| **Occurrence** | 1 | 4.301 | 0.065 | 0.121 |
| **Trophy** | 2 | 2.705 | 0.120 | 0.160 |
| **Dispersal** | 1 | 0.099 | 0.759 | 0.759 |

## Discussion

Our microhaplotype analysis provides a complementary perspective on genetic variation in 12 saproxylic beetle species previously examined using SNPs from the same ddRADseq dataset (Amer et al., 2026; Krovi et al., 2026a,b). Five main patterns emerged. First, comparisons of observed microhaplotype diversity with analytical expectations under linkage equilibrium revealed differences between observed and expected diversity, with the magnitude and direction of these differences varying among species and RAD-locus classes. Second, SNP-based F_ST_ and microhaplotype-based Φ_ST_ recovered significantly concordant patterns of differentiation in all species, although the strength of this correspondence varied considerably among taxa. Third, protection-associated differences in individual microhaplotype homozygosity varied in direction and magnitude among species; the species-specific findings should be interpreted alongside the geographic sampling design and the results of sensitivity analyses. Fourth, associations between site-level haplotype diversity and forest characteristics were similarly species-dependent. Finally, trait analyses suggested possible associations of conservation and occurrence status with interspecific variation in management-associated homozygosity, although none of these trait effects remained significant after FDR correction. Together, these findings demonstrate the value of retaining linked allelic combinations within RAD loci while showing that associations between forest context and genetic variation are species-dependent.

### Microhaplotype diversity differs from expectations under linkage equilibrium

Across all 12 species, observed microhaplotype diversity was lower than its analytical expectation under linkage equilibrium. This consistent direction indicates that the observed combinations of alleles at multi-SNP RAD loci were less diverse than expected from the marginal allele frequencies under the assumption of independence among SNP positions. The magnitude of this difference varied among species, with the largest difference in *E. ferrugineus* and the smallest in *N. haemorrhoidalis*. These differences may reflect non-random associations among constituent SNPs, although sampling variation and the relatively small number of individuals per population may also contribute. Because the comparison was descriptive and did not involve a formal inferential test, these patterns should not be interpreted as statistically confirmed departures from linkage equilibrium. Estimates of haplotype diversity and linkage disequilibrium can be sensitive to sample size, particularly when many genetic markers are examined in relatively small samples (Visscher, 2007). In particular, the relatively small number of individuals sampled per site means that allele and haplotype frequencies may be estimated imprecisely, and rare variants or haplotypes may be underrepresented (Lachance & Tishkoff, 2013; Visscher, 2007). Therefore, the observed differences should be interpreted as descriptive departures from the analytical limkage equilibrium expectation rather than as formal statistical evidence of linkage disequilibrium.

These results illustrate that the diversity of a multi-SNP microhaplotype depends not only on the number of polymorphic positions within a RAD locus but also on the frequencies and combinations of alleles at those positions. Microhaplotypes represent combinations of linked SNPs as multi-allelic markers, thereby retaining information that would be lost if each SNP were considered independently (Kidd & Speed, 2015; Baetscher et al., 2018). Consequently, the diversity expected under independence among SNP positions does not necessarily correspond to the diversity observed in sampled populations. The comparison provides a useful way to describe the additional structure retained when linked SNPs are represented jointly as multi-allelic haplotypes, while emphasizing the need to consider sampling limitations when interpreting within-locus diversity. The combined H1-H2 results show that microhaplotypes retain linked allelic combinations within RAD loci while broadly recovering the patterns of population differentiation identified using SNPs. Populations with similar allele frequencies at individual SNPs may differ in the frequencies of haplotypes formed by combinations of those variants, meaning that microhaplotype informativeness depends not only on the number of SNP positions but also on the frequency distribution of the resulting haplotypes (Kidd & Speed, 2015; Baetscher et al., 2018). This additional representation of within-locus variation can be extracted from existing ddRADseq data without additional sequencing. However, our results demonstrate complementary genetic information rather than universally greater statistical power or superior population-structure inference, which would require a different experimental comparison.

### Forest protection and management context have heterogeneous genetic associations

Site protection and management context showed heterogeneous associations with microhaplotype homozygosity across species, with no consistent directional response. The species-specific protection models identified differences in F_HAP_ for some taxa, but these associations should be interpreted in the context of geographic structure and uneven representation of site categories. In the NE-only management sensitivity analysis, no species showed a statistically significant management association after FDR correction. This regional result does not demonstrate the absence of management effects across the full study area; rather, it indicates that this spatially restricted analysis did not detect statistically significant differences after accounting for multiple testing. Together, these results suggest that associations between forest context and microhaplotype homozygosity are predominantly species-specific rather than general across saproxylic beetles.

This heterogeneity is ecologically plausible because saproxylic beetles differ greatly in substrate specialization, trophic ecology, microclimatic requirements and dispersal. Deadwood tree species, size, decay stage, moisture, sun exposure and spatial continuity can all influence saproxylic assemblages, while intensive forest management often reduces old trees, deadwood and habitat continuity (Grove, 2002; Stokland et al., 2012; Ulyshen & Šobotník, 2018).

Consequently, neither legal protection nor a broad managed-primeval classification necessarily captures the particular habitat conditions relevant to every saproxylic species. Protected forests may generally retain important structural features, but suitable deadwood substrates can also occur in managed or otherwise unprotected forests, while the habitat requirements of individual species may depend on particular tree species, decay stages, microclimates or disturbance histories. Various species-specific genetic responses to forest history and management are also likely determined by spatial and temporal scales in which beetles operate; these animals live in “microcosmos” microhabitats hidden in deadwood (Hjältén et al. 2012). Human-oriented perspective on forest naturalness and effects of management (or protection) could be misleading as it mostly refers to broader landscape scales, as well as to time measured in years, whereas beetles are living locally and breed annually (at least most of species). Moreover, forests are inhabited by thousands or millions of beetle individuals (Gossner et al. 2016), and even rare taxa are represented by hundreds of specimens rather than single individuals. This may contribute to the resilience of saproxylic beetles to environmental changes or result in delayed genetic responses (Gargiulo et al. 2025), which may become apparent only over timescales longer than those associated with modern forest management.

Experimental and observational studies show that deadwood tree identity and diversity, habitat heterogeneity and microclimate strongly affect saproxylic beetle assemblages (Gossner et al., 2016; Seibold et al., 2016, 2017). Moreover, deadwood connectivity can influence saproxylic insect occurrence at relatively local scales (Schiegg, 2000). This ecological complexity may partly explain why both the protected-unprotected and managed-primeval contrasts produced strongly species-dependent patterns rather than a common shift in F_HAP_. The two classifications also capture partly different aspects of forest condition: the former emphasizes site protection and recent management context, whereas the latter represents a broader contrast in forest naturalness and management history. Their broadly heterogeneous outcomes therefore reinforce the importance of considering the ecological context of individual species rather than treating forest management as a single uniform pressure.

Although none of the species showed a statistically significant association between site protection status and F_HAP_ after FDR correction, several species exhibited nominal associations, including *B. reticulatus*, *C. cinnaberinus*, *S. melanura*, *R. inquisitor*, and *B. schneideri*. These patterns suggest that differences in microhaplotype homozygosity between site categories may vary among species, although the evidence remains insufficient to establish consistent effects of protection status across the studied species. The association observed for *C. cinnaberinus* is particularly noteworthy because its negative estimate indicates lower F_HAP_ in unprotected than in protected sites. However, this pattern should not be interpreted as a direct genetic effect of protection. Previous ecological studies indicate that *C. cinnaberinus* responds strongly to specific deadwood and microhabitat characteristics and is not restricted exclusively to protected or old-growth forests (Horák et al., 2010, 2012; Goczał & Rossa, 2017). The species can occupy managed forests where appropriate deadwood resources persist, with factors including sun exposure, deadwood moisture, bark characteristics and decay conditions influencing habitat suitability (Horák et al., 2010, 2012; Goczał & Rossa, 2017). In Białowieża Forest, its occurrence has likewise been related to particular deadwood resources rather than management category alone (Jaworski et al., 2019). Thus, the observed protected-unprotected difference in F_HAP_ may reflect differences in habitat structure, demographic history, spatial population structure or other environmental characteristics associated with protection status rather than protection itself. Moreover, this species has been in expansion since at least 10-20 years (Thomaes et al. 2019), so its demography could have an impact on estimates of genetic diversity (e.g. homozygosity).

Species-specific ecology further illustrates why responses to forest conditions are unlikely to be universal. *E. ferrugineus*, for example, is associated with old hollow trees, and landscape-genetic studies have demonstrated isolation by distance as well as the importance of tree avenues as dispersal corridors in fragmented landscapes (Oleksa et al., 2013, 2015). Conversely, the highly mobile bark beetle *I. typographus* can respond to management through altered dispersal. Montano et al. (2016) documented greater outgoing gene flow from managed toward unmanaged stands, plausibly associated with removal of breeding material from managed forest. In contrast, *R. inquisitor* is a common saproxylic longhorn beetle associated particularly with coniferous forests, where its larvae develop beneath the bark of dead trees, logs and stumps; it can also occur abundantly in recently felled or otherwise disturbed stands where suitable deadwood remains available (Burakowski et al., 1990). Such contrasting habitat and dispersal ecologies make a universal direction of F_HAP_ response unlikely.

The same caution applies to the interpretation of F_HAP_ itself. F_HAP_ measures the proportion of scored microhaplotype loci that are homozygous within an individual; it is not a direct estimator of pedigree inbreeding, autozygosity or inbreeding load. Differences between protected and unprotected sites, or between managed and primeval forests, therefore represent differences in individual microhaplotype homozygosity rather than evidence that forest protection or management directly altered inbreeding. Because the present comparisons are observational and do not fully separate protection status from geography, habitat structure and population history, they should be interpreted as associations rather than causal effects.

### Deadwood diversity provides the central habitat-genetic association

Site-level microhaplotype diversity showed no consistent differences between managed and primeval sites across species. Associations between microhaplotype diversity and forest characteristics also varied among species. In the geography-adjusted models, none of these associations remained statistically significant after correction for multiple testing. In contrast, sensitivity analyses without geographic adjustment revealed positive associations between deadwood diversity and microhaplotype diversity in *B. reticulatus* and *I. typographus*, with the strongest association observed in *B. reticulatus*. These findings indicate that the observed relationships were sensitive to the inclusion of geographic region and should therefore be interpreted as exploratory rather than as evidence of independent environmental effects. An interesting comparison is provided by our recently published SNP-based analysis of the same sampling system (Krovi et al., 2026a). Using population-level observed heterozygosity and the inbreeding coefficient across 13 saproxylic beetle species, that study identified a positive association between deadwood diversity and observed heterozygosity after accounting for geographic region. Deadwood diversity was also negatively associated with F_IS_, whereas total deadwood volume was not significantly associated with either genetic measure. These findings suggest that the diversity of deadwood resources may capture an ecologically relevant aspect of forest structure that is not represented by deadwood quantity alone. However, the SNP-based study also demonstrated substantial variation among species in the direction and precision of species-level relationships, and observed heterozygosity showed limited predictive transferability across regions and species. Thus, the pooled SNP-based association does not imply a uniform response among species or establish genetic diversity as a reliable indicator of local deadwood conditions.

The difference between the SNP-based and microhaplotype-based findings may partly reflect their distinct analytical approaches and genetic response variables. The SNP-based analysis evaluated a community-level association across species while accounting for species identity, whereas the present microhaplotype analysis examined species-specific relationships between site-level haplotype diversity and environmental characteristics. Consequently, a general association across species may coexist with weak or variable species-specific relationships that do not remain significant after multiple-testing correction. Furthermore, observed heterozygosity, F_IS_ and microhaplotype diversity describe different aspects of genetic variation and should not be treated as interchangeable measures. The results from the two studies are therefore complementary: the SNP-based analysis provides evidence for an overall association between deadwood diversity and within-population genetic variation, while the present analysis indicates that this relationship is not consistently detectable at the species level using microhaplotype diversity.

A potential ecological explanation is that deadwood heterogeneity provides a variety of host-tree species, decay conditions, microclimates and associated fungal resources. Saproxylic beetles differ in their substrate requirements and may depend on particular combinations of these resources for reproduction and local persistence. Previous studies have demonstrated that deadwood tree-species diversity and microclimatic heterogeneity influence saproxylic beetle assemblages (Seibold et al., 2016; Gossner et al., 2016). Such heterogeneity could contribute to the persistence of local populations and the maintenance of genetic variation by supporting a wider range of suitable breeding substrates. Nevertheless, these mechanisms were not directly tested in either genetic analysis, and the ecological relevance of deadwood diversity should not be interpreted as proof of a direct effect on genetic diversity.

Geographic structure provides an additional consideration when interpreting these associations. The SNP-based study identified species-specific patterns of effective migration, including pronounced regional discontinuities in some habitat-specialist species and mosaic-like patterns in *B. reticulatus*. These results highlight that local genetic variation is shaped not only by conditions within individual sampling sites but also by broader spatial patterns of population connectivity and demographic history (Krovi et al., 2026a). In particular, previous SNP-based analyses identified pronounced geographic differentiation in *B. reticulatus* (Amer et al., 2026). The sensitivity of the present deadwood-diversity associations to geographic adjustment is therefore important: the positive unadjusted relationships may partly overlap with regional genetic structure rather than represent independent relationships with local deadwood characteristics. However, the available analyses do not establish the precise contribution of geographic structure to these associations.

Taken together, the two genetic approaches suggest that deadwood diversity is an ecologically relevant forest characteristic associated with genetic variation at the broader multi-species level, but its relationship with genetic diversity is not necessarily consistent among individual species. The absence of significant geography-adjusted species-specific associations in the present microhaplotype analysis does not negate the pooled SNP-based result. Instead, it highlights the importance of considering the scale of analysis, genetic metric and species-specific ecology when evaluating environmental associations. Further studies using independent populations sampled across a wider gradient of deadwood conditions, together with more detailed information on population connectivity and demographic history, will be needed to establish whether deadwood diversity has a consistent and independently detectable relationship with microhaplotype diversity.

### Species traits may help explain heterogeneous responses to management

The interspecific trait analysis provides an exploratory perspective on the heterogeneity of management-associated changes in F_HAP_. Conservation status showed the strongest uncorrected relationship with ΔF_HAP_, while occurrence status showed a similar pattern. Neither relationship remained statistically significant after FDR correction, and the analysis included only 12 species. These patterns therefore provide preliminary indications of possible trait-related variation rather than evidence that conservation status or “rarity” consistently predicts genomic responses to forest management. The parallel direction of the uncorrected patterns is nevertheless worth investigating in larger comparative datasets that can reflect ecological specialization, restricted distribution, population size and dependence on habitat structures that may be affected by forest management. This interpretation is consistent with complementary SNP-based analyses of the same dataset, which highlighted metric-dependent patterns of genomic diversity and the contribution of phylogenetic and ecological structure to variation among saproxylic beetle species (Krovi et al., 2026b).

Conservation status showed the strongest uncorrected relationship with ΔF_HAP_: protected species tended to have higher homozygosity in managed relative to primeval forests, whereas unprotected species generally showed smaller and more variable responses. Occurrence status showed a similar pattern, with rare species tending toward more positive ΔF_HAP_ than common species. These two types of beetle clustering are interconnected as the majority of rare species are also protected. Although neither relationship remained significant after FDR correction and the comparison included only 12 species, their parallel direction is noteworthy because conservation status and rarity can reflect ecological specialization, restricted distributions, small population sizes and dependence on habitat structures that are disproportionately lost from managed forests. This interpretation is consistent with the broader ecology of saproxylic insects. Forest management has historically reduced deadwood, veteran trees and habitat continuity, and many threatened saproxylic species are strongly associated with these structures (Grove, 2002; Stokland et al., 2012; Ulyshen & Šobotník, 2018). More generally, maintaining within-species genetic diversity is an important component of conservation because it contributes to evolutionary potential and population persistence, and its monitoring is increasingly recognized as an explicit biodiversity-conservation objective (Hoban et al., 2020, 2021). Our tentative conservation- and occurrence-status patterns therefore suggest that species already characterized by rarity or conservation concern may also show stronger genetic contrasts between managed and primeval habitats. However, future research encompassing a broader range of species will be valuable for determining whether this relationship holds across different taxonomic groups and for establishing its generality.

The trophy showed a weaker but ecologically interesting pattern. Fungivorous species tended toward positive management-associated changes in homozygosity, xylovorous species were predominantly negative, and invertivorous species responded differently. Saproxylic trophy exploit different components of the deadwood trophic web: xylophages use woody tissues directly, fungivores depend on wood-associated fungi, and predators depend partly on the distribution of other saproxylic organisms (Ulyshen & Šobotník, 2018). Recent large-scale trait analyses also demonstrated that trophies respond differently to forest naturalness and habitat context. Wetherbee et al. (2023), analysed 339 obligate saproxylic beetle species and found that trophic and morphological traits helped distinguish near-natural from managed forests, while a recent trait-environment study found positive associations of fungivorous and predatory beetles with old forest in the surrounding landscape (Wetherbee et al., 2023; Bergmark et al., 2024). Thus, the direction of our trophy pattern is biologically plausible, but the non-significant results obtained by us suggest that it should generate hypotheses rather than support a general trophy-level conclusion.

The dispersal category, in contrast, explained little variation in ΔF_HAP_. However, this should not be interpreted as evidence that dispersal is unimportant. Dispersal influences the potential for gene flow and can be critical for persistence in patchy deadwood habitats, but broad categorical assignments may poorly represent realized connectivity, which also depends on habitat distribution, landscape permeability and population history (Schiegg, 2000; Oleksa et al., 2015). Population-genetic studies of saproxylic insects have demonstrated that dispersal limitation can depend strongly on species and habitat configuration. The lack of an association with ΔF_HAP_ may therefore indicate that realized landscape connectivity is more informative than broad species-level dispersal classes for predicting within-species genomic responses to management. Finally, beetle dispersal abilities may be greater than currently assumed due to the limited availability of reliable empirical data (difficulties in tracking beetles over larger distances and mostly indirect signs based on wing morphology; Białkowski et al. 2024). Only some of examined saproxylic beetles have gained genetic support for their dispersal abilities (e.g. Oleksa et al., 2013, 2015). Taken together, our results suggests a potentially useful direction for comparative conservation genomics. Rather than expecting all saproxylic species to respond similarly to forest management, ecological and conservation traits may help identify which species are most likely to show genomic responses and why. Our strongest signals involved conservation and occurrence status, trophy provided weaker ecological differentiation, and dispersal categories provided little explanatory value. Because species’ low sample size (N = 12) and mostly insignificant results after FDR correction, these patterns require validation across further species. Nevertheless, they provide testable hypotheses for larger multispecies datasets and help explain why averaging responses across an ecologically diverse saproxylic assemblage may obscure biologically meaningful differences.

### Study limitation and future directions

The relatively small number of individuals sampled per site may have limited the precision of population-level estimates. In addition, both SNP- and microhaplotype-based analyses were derived from the same ddRADseq data and therefore represent complementary, rather than independent, assessments of genetic differentiation. The uneven geographic distribution of forest categories also limited the separation of management effects from geographic and population-history influences, despite the use of geographic adjustment and sensitivity analyses. Future studies using balanced sampling designs, larger multispecies datasets and complementary genomic and ecological measures will help address these limitations.

## Conclusions

Our study shows that microhaplotypes provide a complementary approach for capturing linked allelic variation within existing ddRADseq datasets. SNP- and microhaplotype-based differentiation showed positive concordance across all 12 species, although the strength of this relationship varied among taxa. Associations between genetic variation and forest conditions were also species-dependent, with no consistent management-related patterns across species. The exploratory relationships between species traits and management-associated homozygosity require further investigation. Overall, our findings highlight the importance of considering species-specific ecology and geographic population structure when interpreting genetic variation in saproxylic beetles, while demonstrating the potential of microhaplotypes as an additional tool in conservation-genetic research.

## Funding support

This study was funded by a grant from the National Science Center, Poland (UMO-2021/43/B/NZ9/00991, PI – Łukasz Kajtoch).

## Conflict of interest

The authors declare no conflicts of interest

## Data Accessibility

Raw sequencing data (FASTQ files) and associated sample metadata have been deposited in the NCBI Sequence Read Archive (SRA) under BioProject accession PRJNA1399704 (https://www.ncbi.nlm.nih.gov/sra/PRJNA1399704).

Filtered single-nucleotide polymorphism (SNP) datasets generated from double-digest RAD sequencing (ddRAD-seq) for multiple saproxylic beetle species are available as species-specific Variant Call Format (VCF) files in RepoOD of the Institute of Systematics and Animal Evolution PAS (https://doi.org/10.18150/MYUQJG).

## Benefit-Sharing section

The whole sampling was conducted in Poland; therefore, the Nagoya Protocol is not applicable. Benefits from this research accrue from the sharing of our data and results on public databases as described above.

## Ethical statement

Collection of protected beetle species and in protected areas was executed according to permissions granted by Polish Ministry of Environment (DOP-1.61.61.2021.TP 1516071.4994962.3977511 and DOP-1.61.61.2021.TP 1516071.4994586.3977515), General Directorate of Environmental Protection (DZP-WF.6401.68.2021.AS) and Regional Directorates of Environmental Protection in Białystok (WPN.6205.74.2021.MM), Rzeszów (WPN.6205.116.2021.MP.2 and WPN.6205.114.2021.ŁL.2), Kielce (WPN.I.6205.1.49.2021.EJP.2) and Wrocław (WPN.6205.143.2021.AR).

## Patient consent statement

Not applicable

## Permission to reproduce material from other sources

Not applicable

## Clinical trial registration

Not applicable

## Declaration of AI and AI-Assisted Technologies

During the preparation of this work, the authors utilized (chat-gpt AI) in order to polish academic grammar, streamline sentence structures, and refine the narrative flow of response templates during peer review. After using this tool, the authors thoroughly reviewed and edited the content as needed and took full responsibility for the ultimate scientific and statistical validity of the manuscript.

## Notes

https://doi.org/10.18150/MYUQJG

https://www.ncbi.nlm.nih.gov/sra/PRJNA1399704

### Competing Interest Statement

The authors have declared no competing interest.

## References

Amer, N. R., et al. (2026). Drivers of genetic diversity in keystone forest species: Contrasting responses of rare and common saproxylic beetles to habitat location and naturalness. Molecular Ecology, 35, e70454. 10.1111/mec.70454

Andrews, K. R., Good, J. M., Miller, M. R., Luikart, G. & Hohenlohe, P. A. (2016). Harnessing the power of RADseq for ecological and evolutionary genomics. Nature Reviews Genetics, 17, 81–92. 10.1038/nrg.2015.28

Andrews, S. (2010). FastQC: A quality control tool for high throughput sequence data. Babraham Bioinformatics. https://www.bioinformatics.babraham.ac.uk/projects/fastqc/

Baetscher, D. S., Clemento, A. J., Ng, T. C., Anderson, E. C. & Garza, J. C. (2018). Microhaplotypes provide increased power from short-read DNA sequences for relationship inference. Molecular Ecology Resources, 18, 296–305. 10.1111/1755-0998.12737

Baird, N. A., Etter, P. D., Atwood, T. S., Currey, M. C., Shiver, A. L., Lewis, Z. A., Selker, E. U., Cresko, W. A. & Johnson, E. A. (2008). Rapid SNP discovery and genetic mapping using sequenced RAD markers. PLoS ONE, 3, e3376. 10.1371/journal.pone.0003376

Bayona-Vásquez, N. J., Glenn, T. C., Kieran, T. J., et al. (2019). Adapterama III: Quadruple-indexed, double/triple-enzyme RADseq libraries (2RAD/3RAD). PeerJ, 7, e7724. 10.7717/peerj.7724

Benjamini, Y. & Hochberg, Y. (1995). Controlling the false discovery rate: A practical and powerful approach to multiple testing. Journal of the Royal Statistical Society: Series B (Methodological), 57, 289–300. 10.1111/j.2517-6161.1995.tb02031.x

Bergmark, P., Hjältén, J., Svensson, J., Neumann, W. & Hekkala, A. (2024). Trait-environment interactions of saproxylic beetles as a guide to biodiversity conservation strategies. Journal of Environmental Management, 360, 121080. 10.1016/j.jenvman.2024.121080

Białkowski, J., Rossa, R., Ziemiakowicz, A., Gohli, J., Dymek, J. & Goczał, J. (2024). Evolution, types, and distribution of flight control devices on wings and elytra in bark beetles. Scientific Reports, 14, 6999. 10.1038/s41598-024-57658-y

Brooks, M. E., Kristensen, K., van Benthem, K. J., Magnusson, A., Berg, C. W., Nielsen, A., Skaug, H. J., Mächler, M. & Bolker, B. M. (2017). glmmTMB balances speed and flexibility among packages for zero-inflated generalized linear mixed modeling. The R Journal, 9, 378–400. 10.32614/RJ-2017-066

Burakowski, B., Mroczkowski, M. & Stefańska, J. (1990). Chrząszcze - Coleoptera. Cerambycidae i Bruchidae. Katalog Fauny Polski, XXIII, 15. Państwowe Wydawnictwo Naukowe.

Ewels, P., Magnusson, M., Lundin, S. & Käller, M. (2016). MultiQC: Summarize analysis results for multiple tools and samples in a single report. Bioinformatics, 32, 3047–3048. 10.1093/bioinformatics/btw354

Excoffier, L., Smouse, P. E. & Quattro, J. M. (1992). Analysis of molecular variance inferred from metric distances among DNA haplotypes: Application to human mitochondrial DNA restriction data. Genetics, 131, 479–491. 10.1093/genetics/131.2.479

Gargiulo, R., Budde, K. B. & Heuertz, M. (2025). Mind the lag: Understanding genetic extinction debt for conservation. Trends in Ecology & Evolution, 40, 228–237.

Goczał, J. & Rossa, R. (2017). Dead wood complexity shapes the pattern of the occurrence of threatened saproxylic beetle *Cucujus cinnaberinus*. Polish Journal of Ecology, 65, 158–165. 10.3161/15052249PJE2017.65.1.014

Gossner, M. M., Wende, B., Levick, S., Schall, P., Floren, A., Linsenmair, K. E., Steffan-Dewenter, I., Schulze, E.-D. & Weisser, W. W. (2016). Deadwood enrichment in European forests—Which tree species should be used to promote saproxylic beetle diversity? Biological Conservation, 201, 92–102. 10.1016/j.biocon.2016.06.032

Grove, S. J. (2002). Saproxylic insect ecology and the sustainable management of forests. Annual Review of Ecology and Systematics, 33, 1–23. 10.1146/annurev.ecolsys.33.010802.150507

Hjältén, J., Stenbacka, F., Pettersson, R. B., Gibb, H., Johansson, T., Danell, K. & Hilszczański, J. (2012). Micro and macro-habitat associations in saproxylic beetles: Implications for biodiversity management. PLoS ONE, 7, e41100.

Hoban, S., Bruford, M., D’Urban Jackson, J., et al. (2020). Genetic diversity targets and indicators in the CBD post-2020 Global Biodiversity Framework must be improved. Biological Conservation, 248, 108654. 10.1016/j.biocon.2020.108654

Hoban, S., Bruford, M. W., Funk, W. C., et al. (2021). Global commitments to conserving and monitoring genetic diversity are now necessary and feasible. BioScience, 71, 964–976. 10.1093/biosci/biab054

Horák, J., Vávrová, E. & Chobot, K. (2010). Habitat preferences influencing populations, distribution and conservation of the endangered saproxylic beetle *Cucujus cinnaberinus* (Coleoptera: Cucujidae) at the landscape level. European Journal of Entomology, 107, 81–88. 10.14411/eje.2010.011

Horák, J., Chumanová, E. & Hilszczański, J. (2012). Saproxylic beetle thrives on the openness in management: A case study on the ecological requirements of *Cucujus cinnaberinus* from Central Europe. Insect Conservation and Diversity, 5, 403–413. 10.1111/j.1752-4598.2011.00173.x

Jaworski, T., Plewa, R., Tarwacki, G., Sućko, K., Hilszczański, J. & Horák, J. (2019). Ecologically similar saproxylic beetles depend on diversified deadwood resources: From habitat requirements to management implications. Forest Ecology and Management, 449, 117462.

Kidd, K. K., Pakstis, A. J., Speed, W. C., Lagacé, R., Chang, J., Wootton, S., Haigh, E. & Kidd, J. R. (2014). Current sequencing technology makes microhaplotypes a powerful new type of genetic marker for forensics. Forensic Science International: Genetics, 12, 215–224. 10.1016/j.fsigen.2014.06.014

Kidd, K. K. & Speed, W. C. (2015). Criteria for selecting microhaplotypes: Mixture detection and deconvolution. Investigative Genetics, 6, 1. 10.1186/s13323-014-0018-3

Knutsen, H., Rukke, B. A., Jorde, P. E. & Ims, R. A. (2000). Genetic differentiation among populations of the beetle *Bolitophagus reticulatus* in a fragmented and a continuous landscape. Heredity, 84, 667–676. 10.1046/j.1365-2540.2000.00691.x

Krovi, R. S., Amer, N. R., Oczkowicz, M. & Kajtoch, Ł. (2025). Meta-analysis of spatial genetic patterns among European saproxylic beetles. Biodiversity and Conservation, 34, 1–27. 10.1007/s10531-024-02940-8

Krovi, R. S., Amer, N. R., Wierzbicka, A., Plewa, R., Kadej, M., Jaworski, T., Smolis, A., Szmatoła, T., Oczkowicz, M. & Kajtoch, Ł. (2026a). Linking deadwood resources, forest connectivity, and genetic diversity for the conservation of saproxylic beetles. Forest Ecology and Management, 625, 124410. 10.1016/j.foreco.2026.124410

Krovi, R. S., Amer, N. R., Wierzbicka, A., Plewa, R., Kadej, M., Jaworski, T., Smolis, A., Szmatoła, T., Oczkowicz, M. & Kajtoch, Ł. (2026b). Metric-dependent genomic diversity reveals phylogenetic and ecological structure in saproxylic beetles. bioRxiv. 10.64898/2026.08.30.748084

Lachance, J. & Tishkoff, S. A. (2013). SNP ascertainment bias in population genetic analyses: Why it is important, and how to correct it. BioEssays, 35, 780–786. 10.1002/bies.201300014

Mantel, N. (1967). The detection of disease clustering and a generalized regression approach. Cancer Research, 27, 209–220.

Montano, V., Bertheau, C., Doležal, P., Krumböck, S., Okrouhlík, J., Stauffer, C. & Moodley, Y. (2016). How differential management strategies affect *Ips typographus* L. dispersal. Forest Ecology and Management, 360, 195–204. 10.1016/j.foreco.2015.10.037

Oettel, J. & Lapin, K. (2021). Linking forest management and biodiversity indicators to strengthen sustainable forest management in Europe. Ecological Indicators, 122, 107275. 10.1016/j.ecolind.2020.107275

Oksanen, J., Simpson, G. L., Blanchet, F. G., Kindt, R., Legendre, P., Minchin, P. R., O’Hara, R. B., Solymos, P., Stevens, M. H. H., Szoecs, E., Wagner, H., Bedward, M., Bolker, B., Borcard, D., Carvalho, G., De Caceres, M., Durand, S., Evangelista, H. B. A., Hannigan, G., Hill, M. O., Lahti, L., Martino, C., Ouellette, M.-H., Ribeiro Cunha, E., Smith, T., Stier, A., Ter Braak, C. J. F. & Weedon, J. (2026). vegan: Community Ecology Package. R package version 2.7-6. 10.32614/CRAN.package.vegan

Oleksa, A., Chybicki, I. J., Gawroński, R., Svensson, G. P. & Burczyk, J. (2013). Isolation by distance in saproxylic beetles may increase with niche specialization. Journal of Insect Conservation, 17, 219–233. 10.1007/s10841-012-9499-7

Oleksa, A., Chybicki, I. J., Larsson, M. C., Svensson, G. P. & Gawroński, R. (2015). Rural avenues as dispersal corridors for the vulnerable saproxylic beetle *Elater ferrugineus* in a fragmented agricultural landscape. Journal of Insect Conservation, 19, 567–580. 10.1007/s10841-015-9778-1

Peterson, B. K., Weber, J. N., Kay, E. H., Fisher, H. S. & Hoekstra, H. E. (2012). Double digest RADseq: An inexpensive method for de novo SNP discovery and genotyping in model and non-model species. PLoS ONE, 7, e37135. 10.1371/journal.pone.0037135

R Core Team. (2024). R: A language and environment for statistical computing. R Foundation for Statistical Computing, Vienna, Austria.

Rochette, N. C., Rivera-Colón, A. G. & Catchen, J. M. (2019). Stacks 2: Analytical methods for paired-end sequencing improve RADseq-based population genomics. Molecular Ecology, 28, 4737–4754. 10.1111/mec.15253

Schiegg, K. (2000). Effects of dead wood volume and connectivity on saproxylic insect species diversity. Écoscience, 7, 290–298. 10.1080/11956860.2000.11682598

Seibold, S., Bässler, C., Brandl, R., Büche, B., Szallies, A., Thorn, S., Ulyshen, M. D. & Müller, J. (2016). Microclimate and habitat heterogeneity as the major drivers of beetle diversity in dead wood. Journal of Applied Ecology, 53, 934–943. 10.1111/1365-2664.12607

Seibold, S., Bässler, C., Brandl, R., Fahrig, L., Förster, B., Heurich, M., Hothorn, T., Scheipl, F., Thorn, S. & Müller, J. (2017). An experimental test of the habitat-amount hypothesis for saproxylic beetles in a forested region. Ecology, 98, 1613–1622. 10.1002/ecy.1819

Slatkin, M. (2008). Linkage disequilibrium - Understanding the evolutionary past and mapping the medical future. Nature Reviews Genetics, 9, 477–485. 10.1038/nrg2361

Stokland, J. N., Siitonen, J. & Jonsson, B. G. (2012). Biodiversity in dead wood. Cambridge University Press.

Thomaes, A., Marchand, S., Lammerant, R. & Crèvecoeur, L. (2019). Hoe komt een oerbossoort in onze populierenbossen terecht?: De Vermiljoenkever: een nieuwe habitatrichtlijnsoort voor Vlaanderen. Natuur.Focus, 18, 19–23.

Ulyshen, M. D. & Šobotník, J. (2018). An introduction to the diversity, ecology, and conservation of saproxylic insects. In M. D. Ulyshen (Ed.), Saproxylic insects: Diversity, ecology and conservation (pp. 1–47). Springer.

Visscher, P. M. (2007). Variation of estimates of SNP and haplotype diversity and linkage disequilibrium in samples from the same population due to experimental and evolutionary sample size. Annals of Human Genetics, 71, 119–126. 10.1111/j.1469-1809.2006.00305.x

Weir, B. S. & Cockerham, C. C. (1984). Estimating F-statistics for the analysis of population structure. Evolution, 38, 1358–1370. 10.1111/j.1558-5646.1984.tb05657.x

Wetherbee, R., et al. (2023). Saproxylic beetles’ morphological traits and higher trophic guilds indicate boreal forest naturalness. Ecology and Evolution, 13, e10739. 10.1002/ece3.10739

Wickham, H. (2016). ggplot2: Elegant graphics for data analysis. Springer. 10.1007/978-3-319-24277-4

Wickham, H., Averick, M., Bryan, J., Chang, W., McGowan, L. D., François, R., Grolemund, G., Hayes, A., Henry, L., Hester, J., Kuhn, M., Pedersen, T. L., Miller, E., Bache, S. M., Müller, K., Ooms, J., Robinson, D., Seidel, D. P., Spinu, V., Takahashi, K., Vaughan, D., Wilke, C., Woo, K. & Yutani, H. (2019). Welcome to the tidyverse. Journal of Open Source Software, 4, 1686. 10.21105/joss.01686

