## Supplementary file for "Contrasting microhaplotype diversity responses of saproxylic beetles to forest management"

Submitted to *Molecular ecology*

**Table S1.** Sampling sites of the 12 studied saproxylic beetle species with corresponding geographic coordinates, forest locations, management category, and site’s characteristics. Deadwood volume (V total) and living wood volume (Wood volume) are reported in m^3^/ha; Deadwood diversity (Dw_diversity), veteran-tree abundance (veterans) and number of tree species (Tree diversity) are per 500-m² plot; time since logging (Latest Logging) are in years.

| **Species** | **Forest** | **site** | **Latitude** | **Longitude** | **Geographic region** | **Management Forest** | **Management Site** | **Conservation** | V total (m3/ha ) | Dw diversity | Veterans (N per plot) | Wood volume (m3/ha) | Tree diversity | Latest Logging (year) |
| --- | --- | --- | --- | --- | --- | --- | --- | --- | --- | --- | --- | --- | --- | --- |
| *Neomida haemorrhoidalis* | AF | NhAFa | 54.02658 | 23.50595 | NE | Managed | Reserve | Protected | 1.6 | 2 | 0 | 268 | 7 | 42 |
| *Neomida haemorrhoidalis* | AF | NhAFb | 53.87032 | 23.30585 | NE | Managed | Reserve | Protected | 3.4 | 4 | 0 | 496 | 8 | 66 |
| *Neomida haemorrhoidalis* | AF | NhAFc | 53.83816 | 23.42504 | NE | Managed | Logged | Unprotected | 2.3 | 2 | 0 | 362 | 6 | 5 |
| *Neomida haemorrhoidalis* | AF | NhAFd | 53.92646 | 23.42419 | NE | Managed | Logged | Unprotected | 3.2 | 2 | 0 | 241 | 6 | 5 |
| *Neomida haemorrhoidalis* | AF | NhAFe | 53.9016 | 23.13949 | NE | Managed | Logged | Unprotected | 1.2 | 1 | 1 | 467 | 4 | 0 |
| *Neomida haemorrhoidalis* | BF | NhBFa | 52.87256 | 23.62713 | NE | Primeval | Primeval not protected | Unprotected | 6.1 | 2 | 0 | 281 | 8 | 10 |
| *Neomida haemorrhoidalis* | BF | NhBFb | 52.87669 | 23.61505 | NE | Primeval | Primeval not protected | Unprotected | 4.4 | 1 | 2 | 286 | 10 | 0 |
| *Neomida haemorrhoidalis* | BF | NhBFc | 52.63199 | 23.69212 | NE | Primeval | Primeval protected | Protected | 4.8 | 2 | 0 | 288 | 7 | 22 |
| *Neomida haemorrhoidalis* | BF | NhBFd | 52.61999 | 23.72195 | NE | Primeval | Primeval protected | Protected | 3.9 | 2 | 0 | 325 | 12 | 10 |
| *Neomida haemorrhoidalis* | BF | NhBFe | 52.65035 | 23.76027 | NE | Primeval | Primeval not protected | Unprotected | 8.8 | 2 | 0 | 170 | 8 | 10 |
| *Neomida haemorrhoidalis* | BF | NhBFf | 52.78635 | 23.71268 | NE | Primeval | Primeval not protected | Unprotected | 2.8 | 2 | 0 | 461 | 8 | 5 |
| *Neomida haemorrhoidalis* | CM | NhCMa | 49.51812 | 22.41941 | SE | Primeval | Primeval protected | Protected | 27.6 | 2 | 1 | 673 | 5 | 29 |
| *Neomida haemorrhoidalis* | CM | NhCMb | 49.51529 | 22.42565 | SE | Primeval | Primeval protected | Protected | 12.1 | 1 | 0 | 592 | 4 | 29 |
| *Neomida haemorrhoidalis* | CM | NhCMc | 49.53254 | 22.56843 | SE | Primeval | Primeval protected | Protected | 27.7 | 2 | 3 | 322 | 9 | 29 |
| *Neomida haemorrhoidalis* | CM | NhCMd | 49.5279 | 22.57353 | SE | Primeval | Primeval protected | Protected | 32.7 | 4 | 4 | 417 | 6 | 29 |
| *Neomida haemorrhoidalis* | CM | NhCMe | 49.56737 | 22.58773 | SE | Primeval | Primeval protected | Protected | 22.8 | 3 | 0 | 524 | 3 | 29 |
| *Neomida haemorrhoidalis* | CM | NhCMf | 49.56397 | 22.59064 | SE | Primeval | Primeval protected | Protected | 18.9 | 2 | 2 | 695 | 4 | 29 |
| *Neomida haemorrhoidalis* | DB | NhDBa | 51.49745 | 15.69447 | SW | Managed | Reserve | Protected | 27 | 2 | 2 | 488 | 7 | 23 |
| *Neomida haemorrhoidalis* | DB | NhDBb | 51.50142 | 15.69462 | SW | Managed | Reserve | Protected | 8.7 | 1 | 2 | 369 | 5 | 23 |
| *Neomida haemorrhoidalis* | DB | NhDBc | 51.50322 | 15.69162 | SW | Managed | Reserve | Protected | 5.3 | 2 | 0 | 435 | 6 | 23 |
| *Neomida haemorrhoidalis* | DB | NhDBd | 51.50553 | 15.68472 | SW | Managed | Reserve | Protected | 22.5 | 2 | 1 | 322 | 6 | 23 |
| *Neomida haemorrhoidalis* | DB | NhDBe | 51.50857 | 15.67905 | SW | Managed | Reserve | Protected | 32.9 | 2 | 0 | 432 | 7 | 23 |
| *Neomida haemorrhoidalis* | DB | NhDBf | 51.50145 | 15.68457 | SW | Managed | Logged | Unprotected | 0.9 | 1 | 0 | 182 | 6 | 5 |
| *Neomida haemorrhoidalis* | HM | NhHMa | 50.88694 | 21.10548 | SE | Primeval | Primeval protected | Protected | 21.9 | 1 | 0 | 394 | 5 | 75 |
| *Neomida haemorrhoidalis* | HM | NhHMb | 50.88528 | 21.1053 | SE | Primeval | Primeval protected | Protected | 18.5 | 1 | 1 | 415 | 4 | 75 |
| *Neomida haemorrhoidalis* | HM | NhHMc | 50.888 | 21.09795 | SE | Primeval | Primeval protected | Protected | 19.2 | 2 | 0 | 412 | 4 | 75 |
| *Neomida haemorrhoidalis* | HM | NhHMd | 51.05906 | 20.70155 | SE | Primeval | Primeval protected | Protected | 12.1 | 2 | 0 | 194 | 7 | 72 |
| *Neomida haemorrhoidalis* | HM | NhHMe | 51.05809 | 20.70623 | SE | Primeval | Primeval protected | Protected | 17.6 | 3 | 2 | 317 | 10 | 72 |
| *Neomida haemorrhoidalis* | HM | NhHMf | 51.05453 | 20.70535 | SE | Primeval | Primeval protected | Protected | 5.8 | 2 | 0 | 489 | 5 | 72 |
| *Neomida haemorrhoidalis* | KF | NhKFa | 53.20771 | 23.58145 | NE | Managed | Reserve | Protected | 6.4 | 3 | 0 | 505 | 7 | 38 |
| *Neomida haemorrhoidalis* | KF | NhKFb | 53.33876 | 23.30416 | NE | Managed | Reserve | Protected | 2.6 | 2 | 0 | 387 | 10 | 38 |
| *Neomida haemorrhoidalis* | KF | NhKFc | 53.28704 | 23.11028 | NE | Managed | Reserve | Protected | 1.8 | 1 | 0 | 375 | 9 | 38 |
| *Neomida haemorrhoidalis* | KF | NhKFd | 53.28543 | 23.1081 | NE | Managed | Reserve | Protected | 1.8 | 1 | 0 | 375 | 9 | 38 |
| *Neomida haemorrhoidalis* | KF | NhKFe | 53.29085 | 23.29906 | NE | Managed | Logged | Unprotected | 6.5 | 2 | 0 | 341 | 11 | 5 |
| *Neomida haemorrhoidalis* | SF | NhSFa | 51.53361 | 17.51304 | SW | Managed | Logged | Unprotected | 27 | 2 | 2 | 488 | 7 | 23 |
| *Neomida haemorrhoidalis* | SF | NhSFb | 51.5372 | 17.39519 | SW | Managed | Reserve | Protected | 8.7 | 1 | 2 | 369 | 5 | 23 |
| *Neomida haemorrhoidalis* | SF | NhSFc | 51.50592 | 16.93765 | SW | Managed | Logged | Unprotected | 5.3 | 2 | 0 | 435 | 6 | 23 |
| *Neomida haemorrhoidalis* | SF | NhSFd | 51.46783 | 17.26448 | SW | Managed | Reserve | Protected | 22.5 | 2 | 1 | 322 | 6 | 23 |
| *Neomida haemorrhoidalis* | SF | NhSFe | 51.44053 | 17.24912 | SW | Managed | Logged | Unprotected | 32.9 | 2 | 0 | 432 | 7 | 23 |
| *Neomida haemorrhoidalis* | SF | NhSFf | 51.44053 | 17.24912 | SW | Managed | Logged | Unprotected | 0.9 | 1 | 0 | 182 | 6 | 5 |
| *Boros schneideri* | AF | BsAFa | 53.86137 | 23.33082 | NE | Managed | Reserve | Protected | 7.8 | 3 | 0 | 487 | 6 | 65 |
| *Boros schneideri* | AF | BsAFb | 53.87063 | 23.29963 | NE | Managed | Reserve | Protected | 14.6 | 2 | 1 | 391 | 7 | 66 |
| *Boros schneideri* | AF | BsAFc | 53.87254 | 23.30556 | NE | Managed | Reserve | Protected | 4.5 | 2 | 2 | 477 | 6 | 66 |
| *Boros schneideri* | AF | BsAFd | 53.87649 | 23.29669 | NE | Managed | Reserve | Protected | 8.9 | 2 | 0 | 490 | 5 | 66 |
| *Boros schneideri* | AF | BsAFe | 53.91163 | 23.23372 | NE | Managed | Logged | Unprotected | 3.6 | 1 | 0 | 0 | 3 | 0 |
| *Boros schneideri* | AF | BsAFf | 53.88132 | 23.37915 | NE | Managed | Logged | Unprotected | 1.2 | 1 | 2 | 438 | 5 | 5 |
| *Boros schneideri* | AF | BsAFg | 53.85022 | 23.30655 | NE | Managed | Logged | Unprotected | 1.5 | 1 | 0 | 270 | 6 | 0 |
| *Boros schneideri* | AF | BsAFh | 53.99144 | 23.47705 | NE | Managed | Logged | Unprotected | 16.9 | 2 | 0 | 433 | 6 | 5 |
| *Boros schneideri* | AF | BsAFi | 53.90047 | 23.41637 | NE | Managed | Logged | Unprotected | 1.5 | 1 | 0 | 511 | 5 | 0 |
| *Boros schneideri* | BF | BsBFa | 52.69084 | 23.72788 | NE | Primeval | Primeval not protected | Unprotected | 20.8 | 4 | 0 | 131 | 7 | 10 |
| *Boros schneideri* | BF | BsBFb | 52.69105 | 23.72878 | NE | Primeval | Primeval not protected | Unprotected | 20.8 | 4 | 0 | 131 | 7 | 10 |
| *Boros schneideri* | BF | BsBFc | 52.66844 | 23.76889 | NE | Primeval | Primeval not protected | Unprotected | 6.4 | 2 | 0 | 274 | 7 | 10 |
| *Boros schneideri* | BF | BsBFd | 52.67616 | 23.84657 | NE | Primeval | Primeval not protected | Unprotected | 2.1 | 1 | 0 | 41 | 6 | 5 |
| *Boros schneideri* | BF | BsBFe | 52.68902 | 23.79492 | NE | Primeval | Primeval not protected | Unprotected | 5.7 | 2 | 0 | 262 | 5 | 5 |
| *Boros schneideri* | BF | BsBFf | 52.63309 | 23.71755 | NE | Primeval | Primeval protected | Protected | 5.9 | 2 | 0 | 240 | 5 | 22 |
| *Boros schneideri* | CM | BsCMa | 49.53679 | 22.63026 | SE | Primeval | Primeval protected | Protected | 10.1 | 1 | 2 | 245 | 8 | 5 |
| *Boros schneideri* | CM | BsCMb | 49.59744 | 22.62185 | SE | Primeval | Primeval protected | Protected | 14 | 2 | 4 | 756 | 8 | 30 |
| *Boros schneideri* | CM | BsCMc | 49.43307 | 22.61117 | SE | Primeval | Primeval protected | Protected | 14 | 2 | 2 | 425 | 11 | 0 |
| *Boros schneideri* | CM | BsCMd | 49.43006 | 22.62036 | SE | Primeval | Primeval protected | Protected | 9.8 | 2 | 3 | 396 | 9 | 0 |
| *Boros schneideri* | CM | BsCMe | 49.43367 | 22.63153 | SE | Primeval | Primeval protected | Protected | 16.9 | 2 | 1 | 441 | 5 | 0 |
| *Boros schneideri* | CM | BsCMf | 49.60129 | 22.67311 | SE | Primeval | Primeval protected | Protected | 5.8 | 1 | 4 | 370 | 6 | 5 |
| *Boros schneideri* | HM | BsHMa | 51.06814 | 20.72167 | SE | Primeval | Primeval protected | Protected | 9.7 | 3 | 1 | 578 | 5 | 66 |
| *Boros schneideri* | HM | BsHMb | 51.07565 | 20.76524 | SE | Primeval | Primeval protected | Protected | 4.4 | 1 | 1 | 415 | 4 | 47 |
| *Boros schneideri* | HM | BsHMc | 51.06923 | 20.73328 | SE | Primeval | Primeval protected | Protected | 7.5 | 1 | 1 | 394 | 5 | 47 |
| *Boros schneideri* | HM | BsHMd | 51.08161 | 20.7634 | SE | Primeval | Primeval protected | Protected | 12.1 | 1 | 0 | 194 | 7 | 72 |
| *Boros schneideri* | HM | BsHMe | 51.04339 | 20.68314 | SE | Primeval | Primeval protected | Protected | 17.6 | 3 | 2 | 317 | 10 | 72 |
| *Boros schneideri* | KF | BsKFa | 53.2641 | 23.21572 | NE | Managed | Logged | Unprotected | 7.1 | 1 | 0 | 125 | 1 | 0 |
| *Boros schneideri* | KF | BsKFb | 53.06373 | 23.72765 | NE | Managed | Logged | Unprotected | 1.7 | 2 | 0 | 325 | 11 | 0 |
| *Boros schneideri* | KF | BsKFc | 53.1485 | 23.6818 | NE | Managed | Logged | Unprotected | 5.2 | 3 | 0 | 36 | 6 | 0 |
| *Boros schneideri* | KF | BsKFd | 53.14747 | 23.67947 | NE | Managed | Logged | Unprotected | 5.2 | 3 | 0 | 36 | 6 | 0 |
| *Boros schneideri* | KF | BsKFe | 53.14608 | 23.68136 | NE | Managed | Logged | Unprotected | 5.2 | 3 | 0 | 36 | 6 | 0 |
| *Cucujus cinnaberinus* | AF | CcAFa | 53.8682 | 23.30333 | NE | Managed | Reserve | Protected | 7.6 | 2 | 0 | 489 | 7 | 66 |
| *Cucujus cinnaberinus* | AF | CcAFb | 53.87159 | 23.34875 | NE | Managed | Reserve | Protected | 24.2 | 2 | 1 | 466 | 10 | 65 |
| *Cucujus cinnaberinus* | AF | CcAFc | 53.86998 | 23.36241 | NE | Managed | Logged | Unprotected | 2.4 | 2 | 0 | 276 | 5 | 5 |
| *Cucujus cinnaberinus* | AF | CcAFd | 53.91163 | 23.23372 | NE | Managed | Logged | Unprotected | 3.6 | 1 | 0 | 0 | 3 | 0 |
| *Cucujus cinnaberinus* | AF | CcAFe | 53.84873 | 23.31902 | NE | Managed | Logged | Unprotected | 2.6 | 2 | 0 | 349 | 4 | 5 |
| *Cucujus cinnaberinus* | AF | CcAFf | 53.99144 | 23.47705 | NE | Managed | Logged | Unprotected | 16.9 | 2 | 0 | 433 | 6 | 5 |
| *Cucujus cinnaberinus* | BF | CcBFa | 52.6865 | 23.70881 | NE | Primeval | Primeval not protected | Unprotected | 0.9 | 2 | 0 | 496 | 8 | 10 |
| *Cucujus cinnaberinus* | BF | CcBFb | 52.69084 | 23.72788 | NE | Primeval | Primeval not protected | Unprotected | 20.8 | 4 | 0 | 131 | 7 | 10 |
| *Cucujus cinnaberinus* | BF | CcBFc | 52.66844 | 23.76889 | NE | Primeval | Primeval not protected | Unprotected | 6.4 | 2 | 0 | 274 | 7 | 10 |
| *Cucujus cinnaberinus* | BF | CcBFd | 52.68725 | 23.87886 | NE | Primeval | Primeval not protected | Unprotected | 8.8 | 2 | 0 | 422 | 10 | 10 |
| *Cucujus cinnaberinus* | BF | CcBFe | 52.70284 | 23.88965 | NE | Primeval | Primeval not protected | Unprotected | 2.4 | 2 | 0 | 549 | 12 | 5 |
| *Cucujus cinnaberinus* | CM | CcCMa | 49.69587 | 22.53875 | SE | Primeval | Primeval protected | Protected | 4.7 | 1 | 1 | 499 | 7 | 34 |
| *Cucujus cinnaberinus* | CM | CcCMb | 49.65681 | 22.50074 | SE | Primeval | Primeval protected | Protected | 13.5 | 1 | 2 | 594 | 4 | 29 |
| *Cucujus cinnaberinus* | CM | CcCMc | 49.59091 | 22.63969 | SE | Primeval | Primeval protected | Protected | 14 | 2 | 4 | 577 | 6 | 30 |
| *Cucujus cinnaberinus* | CM | CcCMd | 49.43338 | 22.61107 | SE | Primeval | Primeval not protected | Unprotected | 1.7 | 1 | 2 | 425 | 11 | 10 |
| *Cucujus cinnaberinus* | CM | CcCMe | 49.71049 | 22.73197 | SE | Primeval | Primeval not protected | Unprotected | 3.6 | 3 | 2 | 391 | 10 | 10 |
| *Cucujus cinnaberinus* | CM | CcCMf | 49.68366 | 22.6059 | SE | Primeval | Primeval not protected | Unprotected | 17.4 | 2 | 5 | 407 | 5 | 10 |
| *Cucujus cinnaberinus* | CM | CcCMg | 49.57651 | 22.46964 | SE | Primeval | Primeval protected | Protected | 16.6 | 2 | 3 | 366 | 5 | 25 |
| *Cucujus cinnaberinus* | DB | CcDBa | 51.53947 | 17.33627 | SW | Managed | Reserve | Protected | 13 | 2 | 0 | 408 | 9 | 62 |
| *Cucujus cinnaberinus* | DB | CcDBb | 51.53287 | 17.34944 | SW | Managed | Reserve | Protected | 13 | 2 | 0 | 408 | 9 | 62 |
| *Cucujus cinnaberinus* | HM | CcHMa | 50.93119 | 20.89913 | SE | Primeval | Primeval protected | Protected | 7.7 | 1 | 0 | 520 | 5 | 75 |
| *Cucujus cinnaberinus* | HM | CcHMb | 50.9273 | 20.90871 | SE | Primeval | Primeval protected | Protected | 21.9 | 1 | 0 | 403 | 7 | 75 |
| *Cucujus cinnaberinus* | HM | CcHMc | 50.88528 | 21.11149 | SE | Primeval | Primeval protected | Protected | 6.4 | 1 | 2 | 392 | 4 | 75 |
| *Cucujus cinnaberinus* | HM | CcHMd | 51.06864 | 20.72184 | SE | Primeval | Primeval not protected | Unprotected | 9.7 | 3 | 1 | 322 | 6 | 5 |
| *Cucujus cinnaberinus* | HM | CcHMe | 50.96528 | 20.4835 | SE | Primeval | Primeval protected | Protected | 6.2 | 3 | 0 | 569 | 6 | 31 |
| *Cucujus cinnaberinus* | KF | CcKFa | 53.20318 | 23.57882 | NE | Managed | Reserve | Protected | 11.7 | 3 | 0 | 600 | 6 | 38 |
| *Cucujus cinnaberinus* | KF | CcKFb | 53.41062 | 23.38001 | NE | Managed | Reserve | Protected | 7.1 | 1 | 1 | 542 | 10 | 35 |
| *Cucujus cinnaberinus* | KF | CcKFc | 53.34254 | 23.30018 | NE | Managed | Reserve | Protected | 8.2 | 3 | 0 | 428 | 11 | 38 |
| *Cucujus cinnaberinus* | KF | CcKFd | 53.27662 | 23.38033 | NE | Managed | Reserve | Protected | 9.8 | 3 | 2 | 503 | 10 | 38 |
| *Cucujus cinnaberinus* | KF | CcKFe | 53.28903 | 23.13414 | NE | Managed | Logged | Unprotected | 2.2 | 2 | 0 | 456 | 8 | 5 |
| *Cucujus cinnaberinus* | KF | CcKFf | 53.2756 | 23.3047 | NE | Managed | Logged | Unprotected | 1.5 | 2 | 0 | 177 | 11 | 10 |
| *Cucujus cinnaberinus* | KF | CcKFg | 53.2641 | 23.21572 | NE | Managed | Logged | Unprotected | 7.1 | 1 | 0 | 125 | 1 | 0 |
| *Cucujus cinnaberinus* | KF | CcKFh | 53.11465 | 23.82827 | NE | Managed | Logged | Unprotected | 11.8 | 1 | 0 | 358 | 6 | 5 |
| *Cucujus cinnaberinus* | KF | CcKFi | 53.1481 | 23.6824 | NE | Managed | Logged | Unprotected | 5.2 | 3 | 0 | 36 | 6 | 0 |
| *Cucujus cinnaberinus* | KF | CcKFj | 53.19278 | 23.66817 | NE | Managed | Logged | Unprotected | 0.5 | 2 | 0 | 569 | 6 | 0 |
| *Cucujus cinnaberinus* | OF | CcOF a | 49.93805 | 18.33473 | SW | Managed | Logged | Unprotected | 7.5 | 1 | 2 | 130 | 5 | 10 |
| *Cucujus cinnaberinus* | OF | CcOF b | 50.01098 | 18.26592 | SW | Managed | Logged | Unprotected | 14.1 | 1 | 0 | 581 | 8 | 10 |
| *Cucujus cinnaberinus* | OF | CcOF c | 52.39222 | 14.54056 | SW | Managed | Reserve | Protected | 6.4 | 2 | 2 | 326 | 8 | 22 |
| *Cucujus cinnaberinus* | OF | CcOF d | 52.36735 | 14.56076 | SW | Managed | Reserve | Protected | 11.8 | 3 | 2 | 440 | 3 | 22 |
| *Elater ferrugineus* | BF | EfBFa | 52.84866 | 23.87789 | NE | Primeval | Primeval not protected | Unprotected | 2.1 | 2 | 5 | 417 | 9 | 10 |
| *Elater ferrugineus* | BF | EfBFb | 52.81759 | 23.75926 | NE | Primeval | Primeval protected | Protected | 1.8 | 2 | 2 | 364 | 10 | 22 |
| *Elater ferrugineus* | BF | EfBFc | 52.83307 | 23.68388 | NE | Primeval | Primeval not protected | Unprotected | 4.6 | 1 | 2 | 387 | 8 | 10 |
| *Elater ferrugineus* | BF | EfBFd | 52.65427 | 23.65124 | NE | Primeval | Primeval protected | Protected | 1 | 1 | 3 | 387 | 10 | 46 |
| *Elater ferrugineus* | BF | EfBFe | 52.6406 | 23.69784 | NE | Primeval | Primeval protected | Protected | 1.3 | 1 | 4 | 410 | 9 | 22 |
| *Elater ferrugineus* | BF | EfBFf | 52.6522 | 23.7734 | NE | Primeval | Primeval protected | Protected | 0 | 0 | 5 | 495 | 9 | 22 |
| *Elater ferrugineus* | BF | EfBFg | 52.7419 | 23.7762 | NE | Primeval | Primeval not protected | Unprotected | 3.6 | 2 | 2 | 45 | 6 | 5 |
| *Elater ferrugineus* | BF | EfBFh | 52.6932 | 23.8137 | NE | Primeval | Primeval protected | Protected | 3.5 | 2 | 2 | 338 | 8 | 56 |
| *Elater ferrugineus* | DB | EfDBa | 51.50814 | 17.02761 | SW | Managed | Reserve | Protected | 70 | 3 | 0 | 440 | 9 | 38 |
| *Elater ferrugineus* | DB | EfDBe | 51.51619 | 17.02687 | SW | Managed | Logged | Unprotected | 22.9 | 4 | 3 | 475 | 6 | 5 |
| *Elater ferrugineus* | DB | EfDBf | 51.47857 | 16.92432 | SW | Managed | Logged | Unprotected | 22.9 | 4 | 3 | 475 | 6 | 5 |
| *Elater ferrugineus* | DB | EfDBi | 51.47857 | 16.92432 | SW | Managed | Logged | Unprotected | 15.7 | 2 | 2 | 475 | 9 | 5 |
| *Elater ferrugineus* | DB | EfDBj | 51.59687 | 17.24105 | SW | Managed | Logged | Unprotected | 15.7 | 2 | 2 | 533 | 2 | 5 |
| *Elater ferrugineus* | HM | EfHMa | 51.05906 | 20.70155 | SE | Primeval | Primeval protected | Protected | 17.6 | 3 | 2 | 194 | 7 | 72 |
| *Elater ferrugineus* | HM | EfHMb | 50.92679 | 20.9198 | SE | Primeval | Primeval protected | Protected | 6.4 | 1 | 2 | 560 | 10 | 75 |
| *Elater ferrugineus* | OF | EfOFa | 50.94633 | 17.32814 | SW | Managed | Reserve | Protected | 63.8 | 4 | 2 | 555 | 11 | 29 |
| *Elater ferrugineus* | OF | EfOFb | 50.94215 | 17.3905 | SW | Managed | Reserve | Protected | 3.5 | 2 | 0 | 357 | 5 | 67 |
| *Elater ferrugineus* | OF | EfOFc | 50.94215 | 17.3905 | SW | Managed | Reserve | Protected | 2.2 | 1 | 1 | 447 | 3 | 67 |
| *Elater ferrugineus* | SF | EfSFf | 51.34233 | 15.43239 | SW | Managed | Logged | Unprotected | 14 | 1 | 4 | 277 | 11 | 10 |
| *Elater ferrugineus* | SF | EfSFg | 51.39802 | 15.46055 | SW | Managed | Logged | Unprotected | 9.4 | 1 | 1 | 429 | 5 | 10 |
| *Elater ferrugineus* | SF | EfSFh | 51.33381 | 15.42478 | SW | Managed | Logged | Unprotected | 0.5 | 1 | 1 | 508 | 16 | 10 |
| *Bolitophagus reticulatus* | AF | BrAFa | 54.04326 | 23.4698 | NE | Managed | Reserve | Protected | 2.6 | 3 | 0 | 221 | 5 | 42 |
| *Bolitophagus reticulatus* | AF | BrAFb | 54.02658 | 23.50595 | NE | Managed | Reserve | Protected | 1.6 | 2 | 0 | 268 | 7 | 42 |
| *Bolitophagus reticulatus* | AF | BrAFc | 53.88049 | 23.33836 | NE | Managed | Reserve | Protected | 2.8 | 2 | 0 | 382 | 5 | 37 |
| *Bolitophagus reticulatus* | AF | BrAFd | 54.06116 | 23.36167 | NE | Managed | Logged | Unprotected | 1.2 | 1 | 0 | 350 | 10 | 5 |
| *Bolitophagus reticulatus* | AF | BrAFe | 53.92646 | 23.42419 | NE | Managed | Logged | Unprotected | 3.2 | 2 | 0 | 241 | 6 | 5 |
| *Bolitophagus reticulatus* | AF | BrAFf | 53.9016 | 23.13949 | NE | Managed | Logged | Unprotected | 1.2 | 1 | 1 | 467 | 4 | 0 |
| *Bolitophagus reticulatus* | BF | BrBFa | 52.70405 | 23.6943 | NE | Primeval | Primeval protected | Protected | 11.7 | 3 | 0 | 309 | 8 | 56 |
| *Bolitophagus reticulatus* | BF | BrBFb | 52.78707 | 23.79641 | NE | Primeval | Primeval not protected | Unprotected | 1.8 | 2 | 2 | 320 | 7 | 10 |
| *Bolitophagus reticulatus* | BF | BrBFc | 52.87256 | 23.62713 | NE | Primeval | Primeval not protected | Unprotected | 6.1 | 2 | 0 | 281 | 8 | 10 |
| *Bolitophagus reticulatus* | BF | BrBFd | 52.63341 | 23.72864 | NE | Primeval | Primeval not protected | Unprotected | 5.9 | 1 | 0 | 376 | 9 | 10 |
| *Bolitophagus reticulatus* | BF | BrBFe | 52.7992 | 23.6435 | NE | Primeval | Primeval protected | Protected | 1.8 | 2 | 0 | 170 | 5 | 22 |
| *Bolitophagus reticulatus* | CM | BrCMa | 49.65622 | 22.49454 | SE | Primeval | Primeval protected | Protected | 28.7 | 2 | 2 | 446 | 7 | 29 |
| *Bolitophagus reticulatus* | CM | BrCMb | 49.57294 | 22.47413 | SE | Primeval | Primeval protected | Protected | 31.1 | 2 | 0 | 342 | 3 | 25 |
| *Bolitophagus reticulatus* | CM | BrCMc | 49.59552 | 22.63324 | SE | Primeval | Primeval protected | Protected | 14 | 2 | 4 | 627 | 5 | 30 |
| *Bolitophagus reticulatus* | CM | BrCMd | 49.53272 | 22.56863 | SE | Primeval | Primeval protected | Protected | 35.4 | 2 | 1 | 322 | 9 | 29 |
| *Bolitophagus reticulatus* | CM | BrCMe | 49.56705 | 22.58474 | SE | Primeval | Primeval protected | Protected | 22.8 | 3 | 0 | 524 | 3 | 29 |
| *Bolitophagus reticulatus* | CM | BrCMf | 49.61275 | 22.64283 | SE | Primeval | Primeval not protected | Unprotected | 12.7 | 1 | 2 | 552 | 3 | 10 |
| *Bolitophagus reticulatus* | CM | BrCMg | 49.58261 | 22.47182 | SE | Primeval | Primeval not protected | Unprotected | 13.7 | 1 | 4 | 529 | 3 | 10 |
| *Bolitophagus reticulatus* | CM | BrCMh | 49.55788 | 22.50051 | SE | Primeval | Primeval not protected | Unprotected | 12.3 | 1 | 0 | 442 | 5 | 5 |
| *Bolitophagus reticulatus* | DB | BrDBa | 51.53335 | 17.51282 | SW | Managed | Logged | Unprotected | 0.5 | 1 | 0 | 310 | 10 | 5 |
| *Bolitophagus reticulatus* | DB | BrDBb | 51.47372 | 17.41899 | SW | Managed | Logged | Unprotected | 0.2 | 1 | 1 | 249 | 6 | 5 |
| *Bolitophagus reticulatus* | DB | BrDBc | 51.47358 | 17.42319 | SW | Managed | Logged | Unprotected | 1.2 | 1 | 0 | 515 | 9 | 10 |
| *Bolitophagus reticulatus* | DB | BrDBd | 51.44053 | 17.24912 | SW | Managed | Logged | Unprotected | 4.1 | 1 | 0 | 19 | 4 | 0 |
| *Bolitophagus reticulatus* | DB | BrDBe | 51.43928 | 17.25553 | SW | Managed | Logged | Unprotected | 0.2 | 1 | 0 | 283 | 4 | 5 |
| *Bolitophagus reticulatus* | HM | BrHMa | 50.96621 | 20.47683 | SE | Primeval | Primeval protected | Protected | 6.2 | 3 | 0 | 578 | 5 | 31 |
| *Bolitophagus reticulatus* | HM | BrHMb | 50.88528 | 21.1053 | SE | Primeval | Primeval not protected | Unprotected | 21.9 | 1 | 0 | 415 | 4 | 75 |
| *Bolitophagus reticulatus* | HM | BrHMc | 50.88694 | 21.10548 | SE | Primeval | Primeval not protected | Unprotected | 18.5 | 1 | 1 | 394 | 5 | 75 |
| *Bolitophagus reticulatus* | HM | BrHMd | 51.05906 | 20.70155 | SE | Primeval | Primeval protected | Protected | 12.1 | 1 | 0 | 194 | 7 | 72 |
| *Bolitophagus reticulatus* | HM | BrHMe | 51.05809 | 20.70623 | SE | Primeval | Primeval protected | Protected | 17.6 | 3 | 2 | 317 | 10 | 72 |
| *Bolitophagus reticulatus* | KF | BrKFa | 53.27662 | 23.3823 | NE | Managed | Reserve | Protected | 1.4 | 3 | 0 | 367 | 9 | 38 |
| *Bolitophagus reticulatus* | KF | BrKFb | 53.20771 | 23.58145 | NE | Managed | Reserve | Protected | 6.4 | 3 | 0 | 505 | 7 | 38 |
| *Bolitophagus reticulatus* | KF | BrKFc | 53.33876 | 23.30416 | NE | Managed | Reserve | Protected | 2.6 | 3 | 0 | 387 | 10 | 38 |
| *Bolitophagus reticulatus* | KF | BrKFd | 53.28704 | 23.11028 | NE | Managed | Reserve | Protected | 1.8 | 1 | 0 | 375 | 9 | 38 |
| *Bolitophagus reticulatus* | KF | BrKFe | 53.33912 | 23.32745 | NE | Managed | Logged | Unprotected | 2.2 | 2 | 0 | 391 | 8 | 10 |
| *Bolitophagus reticulatus* | KF | BrKFf | 53.40152 | 23.31926 | NE | Managed | Logged | Unprotected | 2.5 | 1 | 3 | 540 | 8 | 5 |
| *Bolitophagus reticulatus* | KF | BrKFg | 53.3451 | 23.23897 | NE | Managed | Logged | Unprotected | 1.4 | 2 | 0 | 218 | 12 | 10 |
| *Bolitophagus reticulatus* | SF | BrSFa | 51.49745 | 15.69447 | SW | Managed | Reserve | Protected | 27 | 2 | 2 | 488 | 7 | 23 |
| *Bolitophagus reticulatus* | SF | BrSFb | 51.52112 | 15.77239 | SW | Managed | Reserve | Protected | 7.5 | 1 | 3 | 410 | 14 | 23 |
| *Stenurella melanura* | AF | SmAFa | 53.8964 | 23.3154 | NE | Managed | Reserve | Protected | 4.8 | 2 | 2 | 401 | 8 | 37 |
| *Stenurella melanura* | AF | SmAFb | 53.8727 | 23.3052 | NE | Managed | Reserve | Protected | 4.5 | 2 | 2 | 477 | 6 | 66 |
| *Stenurella melanura* | AF | SmAFc | 53.8741 | 23.3486 | NE | Managed | Reserve | Protected | 2.8 | 1 | 0 | 502 | 7 | 65 |
| *Stenurella melanura* | AF | SmAFd | 53.92574 | 23.23145 | NE | Managed | Logged | Unprotected | 0.3 | 1 | 0 | 400 | 6 | 0 |
| *Stenurella melanura* | AF | SmAFe | 53.90109 | 23.21815 | NE | Managed | Logged | Unprotected | 1.4 | 1 | 0 | 368 | 8 | 0 |
| *Stenurella melanura* | AF | SmAFf | 53.90756 | 23.3282 | NE | Managed | Logged | Unprotected | 1.6 | 2 | 0 | 192 | 3 | 0 |
| *Stenurella melanura* | AF | SmAFg | 54.01425 | 23.34953 | NE | Managed | Logged | Unprotected | 0.5 | 1 | 0 | 493 | 8 | 0 |
| *Stenurella melanura* | BF | SmBFa | 52.7406 | 23.7679 | NE | Primeval | Primeval not protected | Unprotected | 7.9 | 1 | 0 | 616 | 8 | 5 |
| *Stenurella melanura* | BF | SmBFb | 52.7457 | 23.772 | NE | Primeval | Primeval protected | Protected | 0.9 | 1 | 0 | 421 | 7 | 22 |
| *Stenurella melanura* | BF | SmBFc | 52.7153 | 23.6569 | NE | Primeval | Primeval protected | Protected | 6.8 | 2 | 1 | 401 | 7 | 56 |
| *Stenurella melanura* | BF | SmBFd | 52.6596 | 23.6211 | NE | Primeval | Primeval not protected | Unprotected | 0.1 | 1 | 0 | 328 | 6 | 10 |
| *Stenurella melanura* | BF | SmBFe | 52.8455 | 23.8437 | NE | Primeval | Primeval not protected | Unprotected | 2 | 2 | 0 | 503 | 8 | 5 |
| *Stenurella melanura* | CM | SmCMa | 49.79927 | 22.66289 | SE | Primeval | Primeval protected | Protected | 0.2 | 1 | 1 | 475 | 0 | 24 |
| *Stenurella melanura* | CM | SmCMb | 49.66089 | 22.50935 | SE | Primeval | Primeval protected | Protected | 17 | 2 | 4 | 499 | 7 | 34 |
| *Stenurella melanura* | CM | SmCMc | 49.69474 | 22.53812 | SE | Primeval | Primeval protected | Protected | 4.7 | 2 | 1 | 356 | 3 | 34 |
| *Stenurella melanura* | CM | SmCMd | 49.51807 | 22.41881 | SE | Primeval | Primeval protected | Protected | 12.1 | 1 | 0 | 673 | 5 | 29 |
| *Stenurella melanura* | CM | SmCMe | 49.45618 | 22.50602 | SE | Primeval | Primeval not protected | Unprotected | 1.8 | 2 | 1 | 263 | 5 | 0 |
| *Stenurella melanura* | DB | SmDBa | 51.51957 | 15.76484 | SW | Managed | Reserve | Protected | 0.1 | 1 | 0 | 387 | 14 | 23 |
| *Stenurella melanura* | DB | SmDBb | 51.5029 | 16.99712 | SW | Managed | Reserve | Protected | 4.2 | 2 | 0 | 478 | 12 | 62 |
| *Stenurella melanura* | DB | SmDBc | 51.50797 | 17.02786 | SW | Managed | Reserve | Protected | 35.9 | 2 | 2 | 440 | 9 | 62 |
| *Stenurella melanura* | DB | SmDBd | 51.50117 | 17.34964 | SW | Managed | Logged | Unprotected | 5 | 2 | 1 | 336 | 11 | 5 |
| *Stenurella melanura* | DB | SmDBe | 51.56313 | 16.84588 | SW | Managed | Logged | Unprotected | 2.9 | 1 | 0 | 262 | 13 | 5 |
| *Stenurella melanura* | HM | SmHMa | 51.05906 | 20.70155 | SE | Primeval | Primeval protected | Protected | 12.1 | 1 | 0 | 194 | 7 | 72 |
| *Stenurella melanura* | HM | SmHMb | 50.96202 | 20.47607 | SE | Primeval | Primeval protected | Protected | 1.1 | 2 | 0 | 452 | 8 | 31 |
| *Stenurella melanura* | HM | SmHMc | 50.9197 | 20.9003 | SE | Primeval | Primeval protected | Protected | 4 | 1 | 2 | 279 | 7 | 75 |
| *Stenurella melanura* | HM | SmHMd | 50.92719 | 20.91948 | SE | Primeval | Primeval protected | Protected | 5.6 | 1 | 2 | 560 | 10 | 75 |
| *Stenurella melanura* | HM | SmHMe | 51.06074 | 20.69682 | SE | Primeval | Primeval not protected | Unprotected | 17.6 | 3 | 2 | 134 | 5 | 0 |
| *Stenurella melanura* | KF | SmKFa | 53.3337 | 23.0622 | NE | Managed | Reserve | Protected | 4.2 | 2 | 1 | 636 | 8 | 35 |
| *Stenurella melanura* | KF | SmKFb | 53.3001 | 23.1263 | NE | Managed | Reserve | Protected | 13.3 | 2 | 3 | 446 | 8 | 38 |
| *Stenurella melanura* | KF | SmKFc | 53.291 | 23.1089 | NE | Managed | Reserve | Protected | 2.2 | 1 | 1 | 451 | 9 | 38 |
| *Stenurella melanura* | KF | SmKFd | 53.1682 | 23.83917 | NE | Managed | Logged | Unprotected | 4 | 3 | 0 | 268 | 7 | 0 |
| *Stenurella melanura* | KF | SmKFe | 53.24049 | 23.7451 | NE | Managed | Logged | Unprotected | 0.2 | 1 | 0 | 233 | 7 | 0 |
| *Stenurella melanura* | KF | SmKFf | 53.17341 | 23.78975 | NE | Managed | Logged | Unprotected | 1.2 | 1 | 0 | 53 | 7 | 0 |
| *Stenurella melanura* | KF | SmKFg | 53.498 | 23.31904 | NE | Managed | Logged | Unprotected | 1.6 | 1 | 2 | 492 | 10 | 5 |
| *Stenurella melanura* | OF | SmOFa | 50.9327 | 17.40988 | SW | Managed | Logged | Unprotected | 2.2 | 1 | 1 | 155 | 7 | 5 |
| *Stenurella melanura* | OF | SmOFb | 50.94335 | 17.39298 | SW | Managed | Reserve | Protected | 17.4 | 2 | 3 | 447 | 4 | 67 |
| *Stenurella melanura* | OF | SmOFc | 50.9275 | 17.40745 | SW | Managed | Logged | Unprotected | 10 | 1 | 1 | 463 | 8 | 5 |
| *Stenurella melanura* | OF | SmOFd | 50.93692 | 17.39472 | SW | Managed | Logged | Unprotected | 0.7 | 2 | 1 | 445 | 8 | 5 |
| *Stenurella melanura* | SF | SmSFa | 51.50753 | 15.70049 | SW | Managed | Reserve | Protected | 27 | 3 | 2 | 298 | 8 | 23 |
| *Stenurella melanura* | SF | SmSFb | 51.34055 | 15.43079 | SW | Managed | Logged | Unprotected | 14 | 2 | 4 | 277 | 11 | 5 |
| *Stenurella melanura* | SF | SmSFc | 51.38291 | 15.41588 | SW | Managed | Logged | Unprotected | 2.5 | 2 | 0 | 202 | 4 | 5 |
| *Stenurella melanura* | SF | SmSFd | 51.50246 | 15.73339 | SW | Managed | Logged | Unprotected | 0.1 | 1 | 0 | 113 | 7 | 5 |
| *Schizotus pectinicornis* | AF | SpAFa | 54.04326 | 23.4698 | NE | Managed | Reserve | Protected | 2.6 | 3 | 0 | 221 | 5 | 42 |
| *Schizotus pectinicornis* | AF | SpAFc | 53.83878 | 23.40427 | NE | Managed | Logged | Unprotected | 0.4 | 2 | 0 | 329 | 8 | 0 |
| *Schizotus pectinicornis* | AF | SpAFd | 54.06116 | 23.36167 | NE | Managed | Logged | Unprotected | 1.2 | 1 | 0 | 350 | 10 | 5 |
| *Schizotus pectinicornis* | AF | SpAFe | 54.06004 | 23.35966 | NE | Managed | Reserve | Protected | 1.2 | 1 | 0 | 505 | 6 | 42 |
| *Schizotus pectinicornis* | BF | SpBFa | 52.67616 | 23.84657 | NE | Primeval | Primeval not protected | Unprotected | 2.1 | 1 | 0 | 41 | 6 | 5 |
| *Schizotus pectinicornis* | BF | SpBFb | 52.6804 | 23.86476 | NE | Primeval | Primeval not protected | Unprotected | 2.7 | 3 | 0 | 422 | 7 | 10 |
| *Schizotus pectinicornis* | BF | SpBFc | 52.68725 | 23.8612 | NE | Primeval | Primeval not protected | Unprotected | 8.8 | 3 | 0 | 422 | 10 | 10 |
| *Schizotus pectinicornis* | BF | SpBFd | 52.70284 | 23.88965 | NE | Primeval | Primeval not protected | Unprotected | 2.4 | 1 | 0 | 549 | 12 | 5 |
| *Schizotus pectinicornis* | BF | SpBFe | 52.72171 | 23.9162 | NE | Primeval | Primeval not protected | Unprotected | 8.5 | 1 | 4 | 478 | 10 | 10 |
| *Schizotus pectinicornis* | CM | SpCMa | 49.69614 | 22.53888 | SE | Primeval | Primeval protected | Protected | 4.7 | 2 | 1 | 499 | 7 | 34 |
| *Schizotus pectinicornis* | CM | SpCMb | 49.53822 | 22.55883 | SE | Primeval | Primeval protected | Protected | 11.2 | 2 | 3 | 468 | 5 | 29 |
| *Schizotus pectinicornis* | CM | SpCMc | 49.56467 | 22.59458 | SE | Primeval | Primeval protected | Protected | 18.9 | 2 | 2 | 695 | 4 | 29 |
| *Schizotus pectinicornis* | CM | SpCMd | 49.591 | 22.63954 | SE | Primeval | Primeval protected | Protected | 14 | 2 | 4 | 577 | 6 | 30 |
| *Schizotus pectinicornis* | CM | SpCMe | 49.70632 | 22.69328 | SE | Primeval | Primeval not protected | Unprotected | 14.5 | 1 | 4 | 391 | 10 | 5 |
| *Schizotus pectinicornis* | DB | SpDBa | 51.53544 | 17.39263 | SW | Managed | Reserve | Protected | 36.8 | 3 | 9 | 389 | 9 | 62 |
| *Schizotus pectinicornis* | DB | SpDBb | 51.53381 | 17.51372 | SW | Managed | Logged | Unprotected | 0.5 | 1 | 0 | 310 | 10 | 5 |
| *Schizotus pectinicornis* | DB | SpDBd | 51.50425 | 16.93835 | SW | Managed | Reserve | Protected | 10.4 | 3 | 1 | 647 | 3 | 62 |
| *Schizotus pectinicornis* | DB | SpDBe | 51.47221 | 17.37708 | SW | Managed | Logged | Unprotected | 0.9 | 1 | 4 | 688 | 5 | 5 |
| *Schizotus pectinicornis* | HM | SpHMa | 50.88485 | 21.11595 | SE | Primeval | Primeval protected | Protected | 18.5 | 1 | 1 | 552 | 6 | 42 |
| *Schizotus pectinicornis* | HM | SpHMb | 51.07638 | 20.71713 | SE | Primeval | Primeval not protected | Unprotected | 7.5 | 1 | 1 | 298 | 11 | 5 |
| *Schizotus pectinicornis* | HM | SpHMc | 51.07723 | 20.76637 | SE | Primeval | Primeval not protected | Unprotected | 3.6 | 3 | 0 | 209 | 5 | 0 |
| *Schizotus pectinicornis* | HM | SpHMd | 51.059 | 20.70225 | SE | Primeval | Primeval protected | Protected | 17.6 | 3 | 2 | 194 | 7 | 72 |
| *Schizotus pectinicornis* | HM | SpHMe | 51.0529 | 20.7027 | SE | Primeval | Primeval protected | Protected | 8 | 2 | 2 | 441 | 6 | 72 |
| *Schizotus pectinicornis* | KF | SpKFa | 53.41062 | 23.38001 | NE | Managed | Reserve | Protected | 7.1 | 1 | 1 | 542 | 10 | 35 |
| *Schizotus pectinicornis* | KF | SpKFb | 53.4156 | 23.37417 | NE | Managed | Reserve | Protected | 4.3 | 3 | 2 | 484 | 8 | 35 |
| *Schizotus pectinicornis* | KF | SpKFc | 53.34254 | 23.30018 | NE | Managed | Reserve | Protected | 15.3 | 2 | 5 | 428 | 11 | 38 |
| *Schizotus pectinicornis* | KF | SpKFd | 53.27662 | 23.38033 | NE | Managed | Reserve | Protected | 9.8 | 2 | 2 | 503 | 10 | 38 |
| *Schizotus pectinicornis* | KF | SpKFe | 53.2807 | 23.37435 | NE | Managed | Reserve | Protected | 7.5 | 3 | 7 | 485 | 11 | 38 |
| *Schizotus pectinicornis* | KF | SpKFf | 53.11465 | 23.82827 | NE | Managed | Logged | Unprotected | 11.8 | 1 | 0 | 358 | 6 | 5 |
| *Schizotus pectinicornis* | KF | SpKFg | 53.11185 | 23.80233 | NE | Managed | Logged | Unprotected | 3.1 | 2 | 2 | 256 | 6 | 5 |
| *Schizotus pectinicornis* | OF | SpOFa | 49.93805 | 18.33473 | SW | Managed | Logged | Unprotected | 7.5 | 1 | 2 | 130 | 5 | 10 |
| *Schizotus pectinicornis* | OF | SpOFb | 50.01342 | 18.2726 | SW | Managed | Logged | Unprotected | 14.1 | 1 | 0 | 389 | 6 | 10 |
| *Schizotus pectinicornis* | OF | SpOFc | 51.04001 | 17.20826 | SW | Managed | Logged | Unprotected | 17.4 | 2 | 3 | 409 | 8 | 5 |
| *Schizotus pectinicornis* | SF | SpSFa | 51.50322 | 15.69162 | SW | Managed | Reserve | Protected | 27 | 2 | 2 | 435 | 6 | 23 |
| *Schizotus pectinicornis* | SF | SpSFb | 51.50553 | 15.68472 | SW | Managed | Reserve | Protected | 8.7 | 1 | 2 | 322 | 6 | 23 |
| *Thanasimus formicarius* | AF | TfAFa | 53.87063 | 23.29963 | NE | Managed | Reserve | Protected | 14.6 | 2 | 1 | 391 | 7 | 66 |
| *Thanasimus formicarius* | AF | TfAFb | 53.87185 | 23.29804 | NE | Managed | Reserve | Protected | 8.9 | 2 | 0 | 398 | 5 | 66 |
| *Thanasimus formicarius* | AF | TfAFc | 53.87063 | 23.29963 | NE | Managed | Reserve | Protected | 7.6 | 2 | 0 | 391 | 7 | 66 |
| *Thanasimus formicarius* | AF | TfAFd | 54.0403 | 23.4705 | NE | Managed | Reserve | Protected | 2.6 | 3 | 0 | 681 | 5 | 42 |
| *Thanasimus formicarius* | AF | TfAFe | 54.0403 | 23.4705 | NE | Managed | Reserve | Protected | 9 | 3 | 0 | 681 | 5 | 42 |
| *Thanasimus formicarius* | AF | TfAFf | 53.84909 | 23.36436 | NE | Managed | Logged | Unprotected | 3.6 | 0 | 0 | 0 | 4 | 5 |
| *Thanasimus formicarius* | AF | TfAFg | 53.90557 | 23.3279 | NE | Managed | Logged | Unprotected | 0 | 0 | 0 | 530 | 7 | 0 |
| *Thanasimus formicarius* | AF | TfAFh | 53.96092 | 23.48048 | NE | Managed | Logged | Unprotected | 1 | 0 | 0 | 329 | 6 | 5 |
| *Thanasimus formicarius* | BF | TfBFa | 52.68286 | 23.72911 | NE | Primeval | Primeval not protected | Unprotected | 5.9 | 2 | 0 | 449 | 6 | 10 |
| *Thanasimus formicarius* | BF | TfBFb | 52.63758 | 23.54996 | NE | Primeval | Primeval not protected | Unprotected | 7.2 | 2 | 1 | 468 | 7 | 10 |
| *Thanasimus formicarius* | BF | TfBFc | 52.7239 | 23.6729 | NE | Primeval | Primeval protected | Protected | 6.9 | 2 | 2 | 513 | 9 | 22 |
| *Thanasimus formicarius* | BF | TfBFd | 52.6663 | 23.7131 | NE | Primeval | Primeval not protected | Unprotected | 5 | 2 | 0 | 278 | 3 | 10 |
| *Thanasimus formicarius* | BF | TfBFe | 52.69684 | 23.88644 | NE | Primeval | Primeval not protected | Unprotected | 0.8 | 1 | 1 | 439 | 15 | 10 |
| *Thanasimus formicarius* | BF | TfBFf | 52.68229 | 23.72703 | NE | Primeval | Primeval not protected | Unprotected | 4.4 | 2 | 0 | 479 | 7 | 5 |
| *Thanasimus formicarius* | CM | TfCMa | 49.42282 | 22.6525 | SE | Primeval | Primeval not protected | Unprotected | 5.8 | 1 | 4 | 337 | 6 | 5 |
| *Thanasimus formicarius* | CM | TfCMb | 49.43037 | 22.64883 | SE | Primeval | Primeval not protected | Unprotected | 6.9 | 3 | 0 | 167 | 9 | 10 |
| *Thanasimus formicarius* | CM | TfCMd | 49.67364 | 22.45568 | SE | Primeval | Primeval not protected | Unprotected | 7.1 | 1 | 0 | 394 | 5 | 5 |
| *Thanasimus formicarius* | CM | TfCMe | 49.64156 | 22.4831 | SE | Primeval | Primeval not protected | Unprotected | 7.1 | 1 | 0 | 291 | 5 | 5 |
| *Thanasimus formicarius* | DB | TfDBa | 51.52531 | 17.33845 | SW | Managed | Logged | Unprotected | 4.2 | 2 | 0 | 272 | 10 | 5 |
| *Thanasimus formicarius* | DB | TfDBb | 51.44725 | 17.26112 | SW | Managed | Logged | Unprotected | 0.2 | 1 | 1 | 212 | 7 | 5 |
| *Thanasimus formicarius* | HM | TfHMa | 51.07723 | 20.76637 | SE | Primeval | Primeval not protected | Unprotected | 3.6 | 3 | 0 | 209 | 5 | 0 |
| *Thanasimus formicarius* | HM | TfHMb | 51.07627 | 20.76533 | SE | Primeval | Primeval not protected | Unprotected | 3.8 | 1 | 0 | 238 | 6 | 0 |
| *Thanasimus formicarius* | HM | TfHMc | 51.04973 | 20.68072 | SE | Primeval | Primeval not protected | Unprotected | 0 | 0 | 0 | 383 | 4 | 5 |
| *Thanasimus formicarius* | HM | TfHMd | 51.05337 | 20.70374 | SE | Primeval | Primeval protected | Protected | 8 | 2 | 2 | 441 | 6 | 72 |
| *Thanasimus formicarius* | HM | TfHMe | 51.05274 | 20.70792 | SE | Primeval | Primeval not protected | Unprotected | 7.5 | 1 | 1 | 492 | 6 | 5 |
| *Thanasimus formicarius* | KF | TfKFa | 53.1481 | 23.6824 | NE | Managed | Logged | Unprotected | 5.2 | 3 | 0 | 36 | 6 | 0 |
| *Thanasimus formicarius* | KF | TfKFb | 53.15397 | 23.88885 | NE | Managed | Logged | Unprotected | 0 | 0 | 0 | 162 | 7 | 10 |
| *Thanasimus formicarius* | KF | TfKFc | 53.15143 | 23.82178 | NE | Managed | Logged | Unprotected | 1.5 | 1 | 0 | 208 | 8 | 0 |
| *Thanasimus formicarius* | KF | TfKFd | 53.12797 | 23.82293 | NE | Managed | Logged | Unprotected | 0 | 0 | 0 | 314 | 8 | 5 |
| *Thanasimus formicarius* | KF | TfKFe | 53.11717 | 23.6134 | NE | Managed | Logged | Unprotected | 0 | 0 | 0 | 409 | 9 | 5 |
| *Thanasimus formicarius* | KF | TfKFf | 53.28903 | 23.13414 | NE | Managed | Logged | Unprotected | 2.2 | 3 | 0 | 456 | 8 | 5 |
| *Thanasimus formicarius* | OF | TfOFa | 51.3129 | 16.42287 | SW | Managed | Logged | Unprotected | 0.5 | 1 | 0 | 186 | 5 | 5 |
| *Thanasimus formicarius* | SF | TfSFa | 51.50107 | 15.69765 | SW | Managed | Reserve | Protected | 3.6 | 3 | 0 | 468 | 8 | 23 |
| *Thanasimus formicarius* | SF | TfSFe | 51.51092 | 15.75758 | SW | Managed | Logged | Unprotected | 6.3 | 1 | 2 | 348 | 8 | 5 |
| *Thanasimus formicarius* | SF | TfSFb | 51.52051 | 15.76959 | SW | Managed | Logged | Unprotected | 1.5 | 2 | 0 | 358 | 9 | 5 |
| *Thanasimus formicarius* | SF | TfSFc | 51.32744 | 15.44299 | SW | Managed | Logged | Unprotected | 0 | 0 | 0 | 286 | 3 | 5 |
| *Thanasimus formicarius* | SF | TfSFd | 51.49886 | 15.90851 | SW | Managed | Logged | Unprotected | 0 | 0 | 1 | 223 | 5 | 5 |
| *Ips acuminatus* | AF | IaAFa | 53.9007 | 23.2199 | NE | Managed | Logged | Unprotected | 1.4 | 1 | 0 | 368 | 8 | 0 |
| *Ips acuminatus* | AF | IaAFb | 53.9255 | 23.2311 | NE | Managed | Logged | Unprotected | 0.3 | 0 | 0 | 400 | 6 | 0 |
| *Ips acuminatus* | AF | IaAFc | 53.9195 | 23.235 | NE | Managed | Logged | Unprotected | 3.5 | 1 | 1 | 563 | 5 | 0 |
| *Ips acuminatus* | AF | IaAFd | 53.8966 | 23.3318 | NE | Managed | Logged | Unprotected | 3.1 | 1 | 0 | 483 | 5 | 0 |
| *Ips acuminatus* | AF | IaAFe | 53.9333 | 23.3149 | NE | Managed | Logged | Unprotected | 0 | 0 | 0 | 288 | 4 | 5 |
| *Ips acuminatus* | AF | IaAFf | 53.9598 | 23.5108 | NE | Managed | Logged | Unprotected | 0.5 | 1 | 0 | 345 | 7 | 0 |
| *Ips acuminatus* | AF | IaAFg | 53.8866 | 23.2276 | NE | Managed | Logged | Unprotected | 1.9 | 2 | 0 | 354 | 4 | 0 |
| *Ips acuminatus* | BF | IaBFa | 52.6383 | 23.5499 | NE | Primeval | Primeval not protected | Unprotected | 7.2 | 2 | 1 | 468 | 7 | 10 |
| *Ips acuminatus* | BF | IaBFb | 52.723 | 23.6073 | NE | Primeval | Primeval protected | Protected | 5 | 2 | 0 | 589 | 6 | 56 |
| *Ips acuminatus* | BF | IaBFc | 52.6865 | 23.7162 | NE | Primeval | Primeval not protected | Unprotected | 0.4 | 2 | 0 | 169 | 6 | 10 |
| *Ips acuminatus* | BF | IaBFd | 52.6775 | 23.8097 | NE | Primeval | Primeval not protected | Unprotected | 1.6 | 2 | 0 | 422 | 7 | 5 |
| *Ips acuminatus* | HM | IaHMa | 51.07565 | 20.76524 | SE | Primeval | Primeval not protected | Unprotected | 3.6 | 3 | 0 | 238 | 6 | 0 |
| *Ips acuminatus* | HM | IaHMb | 50.88561 | 21.07888 | SE | Primeval | Primeval protected | Protected | 2.6 | 2 | 0 | 542 | 11 | 75 |
| *Ips acuminatus* | HM | IaHMc | 50.88581 | 21.07165 | SE | Primeval | Primeval protected | Protected | 5.1 | 2 | 0 | 330 | 12 | 75 |
| *Ips acuminatus* | HM | IaHMd | 50.86765 | 21.06348 | SE | Primeval | Primeval protected | Protected | 1.8 | 2 | 0 | 340 | 10 | 75 |
| *Ips acuminatus* | KF | IaKFa | 53.1451 | 23.797 | NE | Managed | Logged | Unprotected | 4.9 | 2 | 0 | 443 | 7 | 5 |
| *Ips acuminatus* | KF | IaKFb | 53.1489 | 23.7933 | NE | Managed | Logged | Unprotected | 2.5 | 2 | 0 | 51 | 5 | 5 |
| *Ips acuminatus* | KF | IaKFc | 53.1274 | 23.3386 | NE | Managed | Logged | Unprotected | 0 | 0 | 0 | 427 | 6 | 0 |
| *Ips acuminatus* | KF | IaKFd | 53.2802 | 23.3564 | NE | Managed | Logged | Unprotected | 0 | 0 | 0 | 451 | 10 | 5 |
| *Ips acuminatus* | KF | IaKFe | 53.1481 | 23.6824 | NE | Managed | Logged | Unprotected | 5.2 | 3 | 0 | 36 | 6 | 0 |
| *Ips acuminatus* | KF | IaKFf | 53.15143 | 23.82178 | NE | Managed | Logged | Unprotected | 1.5 | 1 | 0 | 208 | 8 | 0 |
| *Ips acuminatus* | OF | IaOFa | 51.00032 | 17.44799 | SW | Managed | Logged | Unprotected | 0.7 | 2 | 0 | 266 | 8 | 0 |
| *Ips acuminatus* | OF | IaOFb | 50.98329 | 17.37798 | SW | Managed | Logged | Unprotected | 10.8 | 3 | 0 | 211 | 4 | 5 |
| *Ips acuminatus* | OF | IaOFc | 51.0107 | 17.39934 | SW | Managed | Logged | Unprotected | 7.8 | 2 | 1 | 144 | 6 | 5 |
| *Ips acuminatus* | OF | IaOFd | 50.78999 | 18.44108 | SW | Managed | Logged | Unprotected | 0.5 | 1 | 0 | 349 | 8 | 5 |
| *Ips acuminatus* | OF | IaOFe | 51.01195 | 17.39893 | SW | Managed | Logged | Unprotected | 1.2 | 2 | 0 | 229 | 9 | 5 |
| *Ips typographus* | DB | ItDBa | 51.48954 | 17.11837 | SW | Managed | Logged | Unprotected | 2.2 | 2 | 0 | 374 | 5 | 5 |
| *Ips typographus* | DB | ItDBb | 51.48791 | 17.11825 | SW | Managed | Logged | Unprotected | 2.2 | 2 | 0 | 278 | 6 | 5 |
| *Ips typographus* | OF | ItOFa | 50.97671 | 17.38246 | SW | Managed | Logged | Unprotected | 10.8 | 3 | 0 | 426 | 6 | 5 |
| *Ips typographus* | OF | ItOFb | 51.39577 | 16.46351 | SW | Managed | Logged | Unprotected | 0 | 0 | 0 | 179 | 10 | 5 |
| *Ips typographus* | OF | ItOFc | 51.28554 | 16.87641 | SW | Managed | Logged | Unprotected | 0.5 | 1 | 0 | 328 | 8 | 5 |
| *Ips typographus* | OF | ItOFd | 51.00996 | 17.3982 | SW | Managed | Logged | Unprotected | 7.8 | 2 | 1 | 320 | 5 | 5 |
| *Ips typographus* | SF | ItSFa | 51.33914 | 15.26158 | SW | Managed | Logged | Unprotected | 0.067858 | 1 | 0 | 278 | 6 | 5 |
| *Ips typographus* | AF | ItAFa | 53.87063 | 23.29963 | NE | Managed | Reserve | Protected | 4.5 | 2 | 2 | 391 | 7 | 66 |
| *Ips typographus* | AF | ItAFb | 53.87004 | 23.30383 | NE | Managed | Reserve | Protected | 4.5 | 2 | 2 | 391 | 7 | 66 |
| *Ips typographus* | AF | ItAFc | 53.87185 | 23.29804 | NE | Managed | Reserve | Protected | 8.9 | 2 | 0 | 398 | 5 | 66 |
| *Ips typographus* | AF | ItAFd | 53.84909 | 23.36436 | NE | Managed | Logged | Unprotected | 3.6 | 1 | 0 | 0 | 4 | 5 |
| *Ips typographus* | AF | ItAFe | 53.99144 | 23.47705 | NE | Managed | Logged | Unprotected | 16.9 | 2 | 0 | 433 | 6 | 5 |
| *Ips typographus* | AF | ItAFf | 53.90557 | 23.3279 | NE | Managed | Logged | Unprotected | 1.6 | 2 | 0 | 530 | 7 | 0 |
| *Ips typographus* | AF | ItAFg | 53.93221 | 23.43924 | NE | Managed | Logged | Unprotected | 0.6 | 2 | 0 | 390 | 7 | 0 |
| *Ips typographus* | AF | ItAFh | 53.95998 | 23.51073 | NE | Managed | Logged | Unprotected | 0.5 | 1 | 0 | 345 | 7 | 0 |
| *Ips typographus* | AF | ItAFi | 53.82715 | 23.48394 | NE | Managed | Logged | Unprotected | 1.4 | 1 | 0 | 368 | 8 | 0 |
| *Ips typographus* | AF | ItAFj | 53.90109 | 23.21815 | NE | Managed | Logged | Unprotected | 1.4 | 1 | 0 | 368 | 8 | 0 |
| *Ips typographus* | BF | ItBFa | 52.71124 | 23.92365 | NE | Primeval | Primeval not protected | Unprotected | 1.1 | 2 | 0 | 308 | 8 | 5 |
| *Ips typographus* | BF | ItBFb | 52.6152 | 23.63888 | NE | Primeval | Primeval protected | Protected | 5.9 | 3 | 2 | 449 | 8 | 46 |
| *Ips typographus* | BF | ItBFc | 52.79247 | 23.79715 | NE | Primeval | Primeval not protected | Unprotected | 1.8 | 2 | 2 | 375 | 5 | 10 |
| *Ips typographus* | BF | ItBFd | 52.67866 | 23.68303 | NE | Primeval | Primeval protected | Protected | 5 | 2 | 0 | 302 | 6 | 22 |
| *Ips typographus* | BF | ItBFe | 52.7239 | 23.6729 | NE | Primeval | Primeval protected | Protected | 6.9 | 2 | 2 | 513 | 9 | 22 |
| *Ips typographus* | BF | ItBFf | 52.7216 | 23.7469 | NE | Primeval | Primeval not protected | Unprotected | 8.2 | 1 | 0 | 406 | 11 | 10 |
| *Ips typographus* | BF | ItBFg | 52.71968 | 23.74731 | NE | Primeval | Primeval protected | Protected | 6.9 | 2 | 2 | 513 | 9 | 22 |
| *Ips typographus* | BF | ItBFh | 52.63758 | 23.54996 | NE | Primeval | Primeval not protected | Unprotected | 7.2 | 2 | 1 | 468 | 7 | 10 |
| *Ips typographus* | CM | ItCMa | 49.28495 | 22.68341 | SE | Primeval | Primeval not protected | Unprotected | 12.4 | 1 | 4 | 237 | 7 | 5 |
| *Ips typographus* | CM | ItCMb | 49.42312 | 22.65213 | SE | Primeval | Primeval not protected | Unprotected | 4.5 | 1 | 2 | 337 | 6 | 5 |
| *Ips typographus* | CM | ItCMc | 49.43045 | 22.64807 | SE | Primeval | Primeval not protected | Unprotected | 1.6 | 2 | 3 | 193 | 4 | 0 |
| *Ips typographus* | CM | ItCMd | 49.43045 | 22.64807 | SE | Primeval | Primeval not protected | Unprotected | 17.7 | 1 | 0 | 167 | 9 | 0 |
| *Ips typographus* | CM | ItCMe | 49.43045 | 22.64807 | SE | Primeval | Primeval not protected | Unprotected | 9.8 | 2 | 3 | 389 | 3 | 0 |
| *Ips typographus* | HM | ItHMb | 51.04654 | 20.66986 | SE | Primeval | Primeval not protected | Unprotected | 1.8 | 3 | 0 | 475 | 7 | 5 |
| *Ips typographus* | HM | ItHMc | 51.0867 | 20.72645 | SE | Primeval | Primeval not protected | Unprotected | 0 | 0 | 0 | 280 | 6 | 0 |
| *Ips typographus* | KF | ItKFb | 53.20596 | 23.58029 | NE | Managed | Reserve | Protected | 5.8 | 2 | 0 | 505 | 7 | 38 |
| *Ips typographus* | KF | ItKFc | 53.34563 | 23.31341 | NE | Managed | Reserve | Protected | 2.5 | 1 | 2 | 266 | 8 | 38 |
| *Ips typographus* | KF | ItKFd | 53.2807 | 23.37435 | NE | Managed | Reserve | Protected | 7.5 | 3 | 7 | 485 | 11 | 38 |
| *Ips typographus* | KF | ItKFe | 53.27746 | 23.11569 | NE | Managed | Reserve | Protected | 8.4 | 1 | 0 | 368 | 6 | 38 |
| *Ips typographus* | KF | ItKFf | 53.30004 | 23.12651 | NE | Managed | Reserve | Protected | 13.3 | 4 | 3 | 446 | 8 | 68 |
| *Ips typographus* | KF | ItKFg | 53.13739 | 23.71744 | NE | Managed | Logged | Unprotected | 0.4 | 1 | 0 | 474 | 8 | 5 |
| *Ips typographus* | KF | ItKFi | 53.1473 | 23.67949 | NE | Managed | Logged | Unprotected | 5.2 | 3 | 0 | 36 | 6 | 0 |
| *Ips typographus* | KF | ItKFj | 53.12163 | 23.86057 | NE | Managed | Logged | Unprotected | 0.5 | 2 | 0 | 242 | 8 | 0 |
| *Ips typographus* | KF | ItKFk | 53.28903 | 23.13414 | NE | Managed | Logged | Unprotected | 2.2 | 3 | 0 | 456 | 8 | 5 |
| *Rhagium inquisitor* | AF | RiAFa | 53.87254 | 23.30556 | NE | Managed | Reserve | Protected | 4.5 | 2 | 2 | 477 | 6 | 66 |
| *Rhagium inquisitor* | AF | RiAFb | 53.87159 | 23.34875 | NE | Managed | Reserve | Protected | 24.2 | 3 | 1 | 466 | 10 | 65 |
| *Rhagium inquisitor* | AF | RiAFc | 54.02699 | 23.48057 | NE | Managed | Reserve | Protected | 12.8 | 2 | 0 | 113 | 6 | 42 |
| *Rhagium inquisitor* | AF | RiAFd | 54.04048 | 23.47056 | NE | Managed | Reserve | Protected | 9 | 3 | 0 | 681 | 5 | 42 |
| *Rhagium inquisitor* | AF | RiAFe | 53.91163 | 23.23372 | NE | Managed | Logged | Unprotected | 3.6 | 1 | 0 | 0 | 3 | 0 |
| *Rhagium inquisitor* | AF | RiAFf | 53.88132 | 23.37915 | NE | Managed | Logged | Unprotected | 1.2 | 1 | 2 | 438 | 5 | 5 |
| *Rhagium inquisitor* | AF | RiAFg | 53.82127 | 23.33982 | NE | Managed | Logged | Unprotected | 4.1 | 1 | 0 | 314 | 5 | 5 |
| *Rhagium inquisitor* | AF | RiAFh | 53.99144 | 23.47705 | NE | Managed | Logged | Unprotected | 16.9 | 2 | 0 | 433 | 6 | 5 |
| *Rhagium inquisitor* | BF | RiBFa | 52.6646 | 23.7307 | NE | Primeval | Primeval not protected | Unprotected | 10.3 | 1 | 0 | 451 | 11 | 10 |
| *Rhagium inquisitor* | BF | RiBFb | 52.71124 | 23.92365 | NE | Primeval | Primeval not protected | Unprotected | 1.1 | 1 | 0 | 308 | 8 | 5 |
| *Rhagium inquisitor* | BF | RiBFc | 52.6292 | 23.7131 | NE | Primeval | Primeval protected | Protected | 0.9 | 1 | 0 | 458 | 6 | 22 |
| *Rhagium inquisitor* | BF | RiBFd | 52.68902 | 23.79492 | NE | Primeval | Primeval not protected | Unprotected | 5.7 | 2 | 0 | 262 | 5 | 5 |
| *Rhagium inquisitor* | BF | RiBFe | 52.85456 | 23.66052 | NE | Primeval | Primeval not protected | Unprotected | 5.1 | 2 | 0 | 480 | 6 | 5 |
| *Rhagium inquisitor* | CM | RiCMa | 49.52831 | 22.57142 | SE | Primeval | Primeval protected | Protected | 35.4 | 2 | 1 | 417 | 6 | 0 |
| *Rhagium inquisitor* | CM | RiCMb | 49.68393 | 22.60343 | SE | Primeval | Primeval not protected | Unprotected | 17.4 | 2 | 5 | 407 | 5 | 10 |
| *Rhagium inquisitor* | CM | RiCMc | 49.43335 | 22.61115 | SE | Primeval | Primeval not protected | Unprotected | 1.7 | 1 | 2 | 425 | 11 | 10 |
| *Rhagium inquisitor* | CM | RiCMd | 49.67153 | 22.47975 | SE | Primeval | Primeval not protected | Unprotected | 2.1 | 2 | 0 | 220 | 6 | 5 |
| *Rhagium inquisitor* | CM | RiCMe | 49.70699 | 22.69346 | SE | Primeval | Primeval not protected | Unprotected | 14.5 | 2 | 4 | 138 | 6 | 5 |
| *Rhagium inquisitor* | DB | RiDBa | 51.5332 | 17.5613 | SW | Managed | Logged | Unprotected | 0.4 | 1 | 1 | 144 | 4 | 0 |
| *Rhagium inquisitor* | DB | RiDBb | 51.43847 | 17.22055 | SW | Managed | Logged | Unprotected | 0.3 | 1 | 0 | 355 | 5 | 5 |
| *Rhagium inquisitor* | DB | RiDBc | 51.53243 | 17.09377 | SW | Managed | Reserve | Protected | 2.9 | 1 | 0 | 402 | 7 | 62 |
| *Rhagium inquisitor* | HM | RiHMa | 51.04342 | 20.75519 | SE | Primeval | Primeval not protected | Unprotected | 2.4 | 2 | 0 | 191 | 9 | 5 |
| *Rhagium inquisitor* | HM | RiHMb | 50.88674 | 21.07608 | SE | Primeval | Primeval protected | Protected | 5.6 | 1 | 2 | 542 | 11 | 75 |
| *Rhagium inquisitor* | HM | RiHMc | 51.08176 | 20.77203 | SE | Primeval | Primeval not protected | Unprotected | 1.1 | 1 | 0 | 273 | 3 | 0 |
| *Rhagium inquisitor* | HM | RiHMd | 50.96656 | 20.47914 | SE | Primeval | Primeval protected | Protected | 6.2 | 1 | 0 | 578 | 5 | 31 |
| *Rhagium inquisitor* | HM | RiHMe | 50.82551 | 21.11321 | SE | Primeval | Primeval not protected | Unprotected | 0.5 | 1 | 0 | 280 | 3 | 0 |
| *Rhagium inquisitor* | KF | RiKFa | 53.20771 | 23.58145 | NE | Managed | Reserve | Protected | 6.4 | 2 | 0 | 505 | 7 | 38 |
| *Rhagium inquisitor* | KF | RiKFb | 53.41062 | 23.38001 | NE | Managed | Reserve | Protected | 7.1 | 1 | 1 | 542 | 10 | 35 |
| *Rhagium inquisitor* | KF | RiKFc | 53.34254 | 23.30018 | NE | Managed | Reserve | Protected | 8.2 | 2 | 0 | 428 | 11 | 38 |
| *Rhagium inquisitor* | KF | RiKFd | 53.30004 | 23.12651 | NE | Managed | Reserve | Protected | 13.3 | 2 | 3 | 446 | 8 | 68 |
| *Rhagium inquisitor* | KF | RiKFe | 53.14203 | 23.67624 | NE | Managed | Logged | Unprotected | 1.9 | 2 | 0 | 13 | 7 | 0 |
| *Rhagium inquisitor* | KF | RiKFf | 53.19278 | 23.66817 | NE | Managed | Logged | Unprotected | 0.5 | 2 | 0 | 569 | 6 | 0 |
| *Rhagium inquisitor* | SF | RiSFa | 51.50253 | 15.73197 | SW | Managed | Logged | Unprotected | 0.5 | 2 | 0 | 330 | 3 | 5 |
| *Rhagium inquisitor* | SF | RiSFb | 51.49848 | 15.71398 | SW | Managed | Logged | Unprotected | 2.5 | 2 | 0 | 344 | 8 | 5 |
| *Phaenops cyanea* | AF | PcAFa | 53.90557 | 23.3279 | NE | Managed | Logged | Unprotected | 1.6 | 2 | 0 | 530 | 7 | 0 |
| *Phaenops cyanea* | AF | PcAFb | 53.82715 | 23.48394 | NE | Managed | Logged | Unprotected | 0.1 | 1 | 0 | 280 | 5 | 0 |
| *Phaenops cyanea* | AF | PcAFc | 53.82868 | 23.48511 | NE | Managed | Logged | Unprotected | 0.1 | 1 | 0 | 280 | 5 | 0 |
| *Phaenops cyanea* | AF | PcAFd | 53.85855 | 23.32973 | NE | Managed | Logged | Unprotected | 0 | 0 | 0 | 0 | 3 | 0 |
| *Phaenops cyanea* | AF | PcAFe | 54.0117 | 23.4922 | NE | Managed | Logged | Unprotected | 0.3 | 1 | 0 | 203 | 4 | 0 |
| *Phaenops cyanea* | BF | PcBFa | 52.60347 | 23.58641 | NE | Primeval | Primeval not protected | Unprotected | 1.5 | 1 | 0 | 324 | 5 | 0 |
| *Phaenops cyanea* | BF | PcBFb | 52.61541 | 23.63855 | NE | Primeval | Primeval protected | Protected | 5.9 | 3 | 2 | 449 | 8 | 46 |
| *Phaenops cyanea* | BF | PcBFc | 52.64549 | 23.70082 | NE | Primeval | Primeval protected | Protected | 3 | 1 | 0 | 241 | 6 | 30 |
| *Phaenops cyanea* | BF | PcBFd | 52.64553 | 23.69587 | NE | Primeval | Primeval not protected | Unprotected | 1.3 | 1 | 4 | 266 | 3 | 10 |
| *Phaenops cyanea* | CM | PcCMa | 49.64591 | 22.50589 | SE | Primeval | Primeval not protected | Unprotected | 2.1 | 2 | 0 | 389 | 5 | 5 |
| *Phaenops cyanea* | CM | PcCMb | 49.67407 | 22.4502 | SE | Primeval | Primeval not protected | Unprotected | 7.1 | 1 | 0 | 394 | 5 | 5 |
| *Phaenops cyanea* | CM | PcCMc | 49.6414 | 22.48374 | SE | Primeval | Primeval not protected | Unprotected | 4.1 | 2 | 2 | 291 | 5 | 10 |
| *Phaenops cyanea* | CM | PcCMd | 49.60468 | 22.58671 | SE | Primeval | Primeval not protected | Unprotected | 2.8 | 2 | 0 | 174 | 8 | 5 |
| *Phaenops cyanea* | CM | PcCMe | 49.68911 | 22.40972 | SE | Primeval | Primeval not protected | Unprotected | 4.5 | 1 | 2 | 325 | 5 | 5 |
| *Phaenops cyanea* | CM | PcCMf | 49.7232 | 22.60576 | SE | Primeval | Primeval not protected | Unprotected | 1.4 | 2 | 2 | 188 | 7 | 5 |
| *Phaenops cyanea* | DB | PcDBa | 51.54507 | 17.02616 | SW | Managed | Logged | Unprotected | 4.2 | 2 | 0 | 274 | 5 | 5 |
| *Phaenops cyanea* | HM | PcHMa | 50.8553 | 20.5904 | SE | Primeval | Primeval not protected | Unprotected | 0.2 | 1 | 0 | 196 | 4 | 0 |
| *Phaenops cyanea* | HM | PcHMb | 51.07217 | 20.75963 | SE | Primeval | Primeval not protected | Unprotected | 3.8 | 1 | 0 | 270 | 3 | 0 |
| *Phaenops cyanea* | HM | PcHMc | 51.07723 | 20.76637 | SE | Primeval | Primeval not protected | Unprotected | 0.6 | 1 | 0 | 238 | 6 | 0 |
| *Phaenops cyanea* | KF | PcKFa | 53.1481 | 23.6824 | NE | Managed | Logged | Unprotected | 5.2 | 2 | 0 | 36 | 6 | 0 |
| *Phaenops cyanea* | KF | PcKFb | 53.13965 | 23.74009 | NE | Managed | Logged | Unprotected | 0.6 | 2 | 0 | 277 | 7 | 5 |
| *Phaenops cyanea* | KF | PcKFc | 53.13806 | 23.72906 | NE | Managed | Logged | Unprotected | 0.5 | 1 | 0 | 377 | 8 | 5 |
| *Phaenops cyanea* | KF | PcKFd | 53.1382 | 23.4458 | NE | Managed | Logged | Unprotected | 0.2 | 1 | 0 | 25 | 7 | 5 |
| *Phaenops cyanea* | KF | PcKFe | 53.13961 | 23.66718 | NE | Managed | Logged | Unprotected | 0 | 0 | 0 | 403 | 4 | 5 |
| *Phaenops cyanea* | SF | PcSFa | 51.39124 | 15.5394 | SW | Managed | Logged | Unprotected | 0.1 | 1 | 0 | 224 | 3 | 5 |
| *Phaenops cyanea* | SF | PcSFb | 51.51323 | 15.75438 | SW | Managed | Logged | Unprotected | 6.3 | 1 | 2 | 363 | 6 | 5 |
| *Phaenops cyanea* | SF | PcSFc | 51.50063 | 15.7214 | SW | Managed | Logged | Unprotected | 3.6 | 3 | 0 | 369 | 5 | 0 |
| *Phaenops cyanea* | SF | PcSFd | 51.4178 | 15.85139 | SW | Managed | Logged | Unprotected | 5.3 | 2 | 0 | 316 | 5 | 5 |

**Table S2.** Characteristics of the 12 saproxylic beetle species included in the genetic analyses, including species-level ecological and conservation attributes.

| **Species** | **Family** | **Occurance** | **Conservation_status** | **Trophy** | **Dispersal** |
| --- | --- | --- | --- | --- | --- |
| *Neomida haemorrhoidalis* | Tenebrionidae | rare | unprotected | fungivorous | moderate |
| *Boros schneideri* | Boridae | rare | protected | invertivorous | moderate |
| *Cucujus cinnaberinus* | Cucujidae | rare | protected | invertivorous | moderate |
| *Elater ferrugineus* | Elateridae | rare | protected | invertivorous | high |
| *Bolitophagus reticulatus* | Tenebrionidae | common | unprotected | fungivorous | moderate |
| *Schizotus pectinicornis* | Pyrochroidae | common | unprotected | invertivorous | moderate |
| *Stenurella melanura* | Cerambycidae | common | unprotected | xylovorous | high |
| *Thanasimus formicarius* | Cleridae | common | unprotected | invertivorous | high |
| *Ips acuminatus* | Curculionidae | common | unprotected | xylovorous | high |
| *Ips typographus* | Curculionidae | common | unprotected | xylovorous | high |
| *Rhagium inquisitor* | Cerambycidae | common | unprotected | xylovorous | moderate |
| *Phaenops cyanea* | Buprestidae | common | unprotected | xylovorous | high |

**Table S3.** **Species-level summary of observed microhaplotype diversity and analytical expectations under linkage equilibrium across 12 saproxylic beetle species.** Mean H_d(obs)_ represents the mean observed microhaplotype diversity across sampling populations and multi-SNP RAD loci, whereas mean H_d(LE)_ represents the corresponding analytical expectation calculated from the marginal allele frequencies of constituent SNP positions under the assumption of independence among SNPs. Mean ΔH_d(LE)_ is the difference between observed and expected diversity (H_d(obs)_ - H_d(LE)_). Only population-locus combinations with data from five diploid individuals were included in the calculations. N_sites indicates the number of sampling sites represented after applying this minimum sample-size criterion. N_loci indicates the total number of population-by-locus observations included in the calculations.

| Species | N_sites | N_loci | Mean H_d(obs)_ | Mean H_d(LE)_ | Mean Δ H_d(LE)_ |
| --- | --- | --- | --- | --- | --- |
| *Neomida haemorrhoidalis* | 40 | 92500 | 0.398 | 0.458 | -0.060 |
| *Bolitophagus reticulatus* | 33 | 22884 | 0.397 | 0.462 | -0.066 |
| *Boros schneideri* | 27 | 32231 | 0.398 | 0.465 | -0.067 |
| *Cucujus cinnaberinus* | 34 | 27658 | 0.388 | 0.453 | -0.065 |
| *Elater ferrugineus* | 13 | 38341 | 0.403 | 0.558 | -0.155 |
| *Bolitophagus reticulatus* | 33 | 22884 | 0.397 | 0.462 | -0.066 |
| *Stenurella melanura* | 31 | 49180 | 0.456 | 0.524 | -0.068 |
| *Schizotus pectinicornis* | 26 | 13144 | 0.393 | 0.464 | -0.071 |
| *Thanasimus formicarius* | 30 | 5550 | 0.400 | 0.496 | -0.096 |
| *Ips acuminatus* | 23 | 14060 | 0.410 | 0.478 | -0.067 |
| *Ips typographus* | 38 | 9582 | 0.403 | 0.466 | -0.063 |
| *Rhagium inquisitor* | 25 | 10267 | 0.421 | 0.485 | -0.064 |
| *Phaenops cyanea* | 28 | 153652 | 0.417 | 0.531 | -0.114 |

**Table S4.** Number of sampling sites per species across protection and forest management categories within each geographic region.

| Species | Geography | Protected sites | Unprotected sites | Primeval sites | Managed sites |
| --- | --- | --- | --- | --- | --- |
| *Neomida haemorrhoidalis* | NE | 7 | 9 | 6 | 10 |
| *Neomida haemorrhoidalis* | SE | 12 | 0 | 12 | 0 |
| *Neomida haemorrhoidalis* | SW | 7 | 5 | 0 | 12 |
| *Boros schneideri* | NE | 5 | 15 | 6 | 14 |
| *Boros schneideri* | SE | 6 | 5 | 11 | 0 |
| *Cucujus cinnaberinus* | NE | 6 | 15 | 5 | 16 |
| *Cucujus cinnaberinus* | SE | 8 | 4 | 12 | 0 |
| *Cucujus cinnaberinus* | SW | 4 | 2 | 0 | 6 |
| *Elater ferrugineus* | NE | 5 | 3 | 8 | 0 |
| *Elater ferrugineus* | SE | 2 | 0 | 2 | 0 |
| *Elater ferrugineus* | SW | 4 | 7 | 0 | 11 |
| *Bolitophagus reticulatus* | NE | 9 | 9 | 5 | 13 |
| *Bolitophagus reticulatus* | SE | 8 | 5 | 13 | 0 |
| *Bolitophagus reticulatus* | SW | 2 | 5 | 0 | 7 |
| *Stenurella melanura* | NE | 8 | 11 | 5 | 14 |
| *Stenurella melanura* | SE | 8 | 2 | 10 | 0 |
| *Stenurella melanura* | SW | 5 | 8 | 0 | 13 |
| *Schizotus pectinicornis* | NE | 7 | 9 | 5 | 11 |
| *Schizotus pectinicornis* | SE | 7 | 3 | 10 | 0 |
| *Schizotus pectinicornis* | SW | 4 | 5 | 0 | 9 |
| *Thanasimus formicarius* | NE | 6 | 14 | 6 | 14 |
| *Thanasimus formicarius* | SE | 1 | 8 | 9 | 0 |
| *Thanasimus formicarius* | SW | 1 | 7 | 0 | 8 |
| *Ips acuminatus* | NE | 1 | 16 | 4 | 13 |
| *Ips acuminatus* | SE | 3 | 1 | 4 | 0 |
| *Ips acuminatus* | SW | 0 | 5 | 0 | 5 |
| *Ips typographus* | NE | 12 | 15 | 8 | 19 |
| *Ips typographus* | SE | 0 | 7 | 7 | 0 |
| *Ips typographus* | SW | 0 | 7 | 0 | 7 |
| *Rhagium inquisitor* | NE | 9 | 10 | 5 | 14 |
| *Rhagium inquisitor* | SE | 3 | 7 | 10 | 0 |
| *Rhagium inquisitor* | SW | 1 | 4 | 0 | 5 |
| *Phaenops cyanea* | NE | 2 | 12 | 4 | 10 |
| *Phaenops cyanea* | SE | 0 | 9 | 9 | 0 |
| *Phaenops cyanea* | SW | 0 | 5 | 0 | 5 |

**Table S5.** Total number of sampling sites per region across geographic regions (NE, SE and SW).

| **Geography** | **Primeval sites** | **Managed sites** | **Protected sites** | **Unprotected sites** | **Total sites** |
| --- | --- | --- | --- | --- | --- |
| **NE** | 67 | 148 | 78 | 137 | 215 |
| **SE** | 109 | 0 | 63 | 46 | 109 |
| **SW** | 0 | 88 | 28 | 60 | 88 |
| **Total** | 176 | 236 | 169 | 243 | 412 |

**Table S6.** Species-specific beta-binomial model results for the **NE-only** sensitivity analysis of forest **management effects** on F_HAP_. E. ferrugineus was excluded because no managed sites were represented in the NE region.

| **Species** | **N inds** | **N sites** | **estimate** | **SE** | **statistic** | **p.value** | **conf.low** | **conf.high** | **AIC** | **P_FDR** |
| --- | --- | --- | --- | --- | --- | --- | --- | --- | --- | --- |
| *Neomida haemorrhoidalis* | 80 | 16 | 0.004 | 0.109 | 0.036 | 0.972 | -0.209 | 0.217 | 1090.671 | 0.972 |
| *Boros schneideri* | 97 | 20 | 0.018 | 0.050 | 0.363 | 0.717 | -0.080 | 0.116 | 1104.423 | 0.867 |
| *Cucujus cinnaberinus* | 103 | 21 | 0.023 | 0.074 | 0.306 | 0.759 | -0.122 | 0.168 | 1180.803 | 0.867 |
| *Bolitophagus reticulatus* | 90 | 18 | -0.156 | 0.172 | -0.909 | 0.363 | -0.493 | 0.181 | 1054.301 | 0.867 |
| *Schizotus pectinicornis* | 93 | 16 | -0.026 | 0.026 | -1.001 | 0.317 | -0.076 | 0.025 | 1025.973 | 0.867 |
| *Stenurella melanura* | 77 | 19 | -0.058 | 0.123 | -0.473 | 0.636 | -0.300 | 0.183 | 825.295 | 0.867 |
| *Thanasimus formicarius* | 99 | 20 | 0.031 | 0.114 | 0.269 | 0.788 | -0.193 | 0.255 | 983.139 | 0.867 |
| *Ips acuminatus* | 84 | 17 | 0.065 | 0.073 | 0.886 | 0.376 | -0.079 | 0.209 | 924.568 | 0.867 |
| *Ips typographus* | 132 | 27 | -0.010 | 0.028 | -0.374 | 0.708 | -0.065 | 0.044 | 1117.155 | 0.867 |
| *Rhagium inquisitor* | 90 | 19 | -0.072 | 0.035 | -2.047 | 0.041 | -0.142 | -0.003 | 814.978 | 0.448 |
| *Phaenops cyanea* | 70 | 14 | -0.087 | 0.172 | -0.507 | 0.612 | -0.424 | 0.249 | 1066.821 | 0.867 |

**Table S7.** Sensitivity analysis of species-specific associations between forest structural and deadwood-related variables and haplotype diversity (H_d_), using models **without geographic region** as a covariate. Estimates, standard errors, test statistics, P-values, 95% confidence intervals, number of sampling sites, model R^2^, and FDR-adjusted P-values are reported. *I. acuminatus* has 23 sites with 0 veterans and 3 sites with only 1 veteran, so “veterans” was excluded.

| Species | Predictor | Estimate | SE | Statistic | P value | conf.low | conf.high | N sites | R^2^ | P_FDR |
| --- | --- | --- | --- | --- | --- | --- | --- | --- | --- | --- |
| *Neomida haemorrhoidalis* | Latest_Logging | 0.000 | 0.000 | -0.409 | 0.685 | -0.001 | 0.000 | 34 | 0.005 | 0.822 |
|  | Dw_diversity | 0.016 | 0.007 | 2.121 | 0.042 | 0.001 | 0.031 | 34 | 0.123 | 0.167 |
|  | V_total | 0.000 | 0.001 | -0.579 | 0.566 | -0.002 | 0.001 | 34 | 0.010 | 0.971 |
|  | Tree_diversity | -0.001 | 0.003 | -0.516 | 0.610 | -0.007 | 0.004 | 34 | 0.008 | 0.732 |
|  | Wood_volume | 0.000 | 0.000 | 0.294 | 0.770 | 0.000 | 0.000 | 34 | 0.003 | 0.925 |
|  | veterans | 0.001 | 0.006 | 0.215 | 0.831 | -0.011 | 0.013 | 34 | 0.001 | 0.910 |
| *Boros schneideri* | Latest_Logging | 0.000 | 0.000 | -2.077 | 0.047 | -0.001 | 0.000 | 31 | 0.129 | 0.100 |
|  | Dw_diversity | -0.001 | 0.005 | -0.184 | 0.855 | -0.011 | 0.009 | 31 | 0.001 | 0.967 |
|  | V_total | 0.000 | 0.001 | 0.007 | 0.994 | -0.002 | 0.002 | 31 | 0.000 | 0.994 |
|  | Tree_diversity | -0.001 | 0.002 | -0.405 | 0.688 | -0.005 | 0.004 | 31 | 0.006 | 0.751 |
|  | Wood_volume | 0.000 | 0.000 | -0.536 | 0.596 | 0.000 | 0.000 | 31 | 0.010 | 0.806 |
|  | veterans | -0.001 | 0.004 | -0.303 | 0.764 | -0.009 | 0.007 | 31 | 0.003 | 0.910 |
| *Cucujus cinnaberinus* | Latest_Logging | 0.000 | 0.000 | -0.879 | 0.386 | -0.001 | 0.000 | 37 | 0.022 | 0.514 |
|  | Dw_diversity | 0.008 | 0.007 | 1.065 | 0.294 | -0.007 | 0.022 | 37 | 0.031 | 0.504 |
|  | V_total | -0.001 | 0.001 | -0.640 | 0.526 | -0.002 | 0.001 | 37 | 0.012 | 0.971 |
|  | Tree_diversity | 0.003 | 0.002 | 1.581 | 0.123 | -0.001 | 0.008 | 37 | 0.067 | 0.319 |
|  | Wood_volume | 0.000 | 0.000 | 0.017 | 0.986 | 0.000 | 0.000 | 37 | 0.000 | 0.986 |
|  | veterans | -0.002 | 0.005 | -0.326 | 0.746 | -0.011 | 0.008 | 37 | 0.003 | 0.910 |
| *Elater ferrugineus* | Latest_Logging | 0.000 | 0.001 | 0.135 | 0.895 | -0.002 | 0.002 | 15 | 0.001 | 0.900 |
|  | Dw_diversity | -0.022 | 0.015 | -1.426 | 0.177 | -0.055 | 0.011 | 15 | 0.135 | 0.355 |
|  | V_total | -0.001 | 0.001 | -0.920 | 0.375 | -0.003 | 0.001 | 15 | 0.061 | 0.899 |
|  | Tree_diversity | -0.003 | 0.009 | -0.306 | 0.765 | -0.021 | 0.016 | 15 | 0.007 | 0.765 |
|  | Wood_volume | 0.000 | 0.000 | -1.272 | 0.226 | 0.000 | 0.000 | 15 | 0.111 | 0.612 |
|  | veterans | 0.002 | 0.014 | 0.115 | 0.910 | -0.029 | 0.033 | 15 | 0.001 | 0.910 |
| *Bolitophagus reticulatus* | Latest_Logging | 0.001 | 0.001 | 2.458 | 0.019 | 0.000 | 0.003 | 38 | 0.144 | 0.100 |
|  | Dw_diversity | 0.050 | 0.016 | 3.101 | 0.004 | 0.017 | 0.082 | 38 | 0.211 | 0.031 |
|  | V_total | 0.002 | 0.001 | 1.787 | 0.082 | 0.000 | 0.005 | 38 | 0.081 | 0.672 |
|  | Tree_diversity | -0.009 | 0.005 | -1.813 | 0.078 | -0.020 | 0.001 | 38 | 0.084 | 0.319 |
|  | Wood_volume | 0.000 | 0.000 | 0.748 | 0.459 | 0.000 | 0.000 | 38 | 0.015 | 0.787 |
|  | veterans | -0.004 | 0.012 | -0.338 | 0.737 | -0.028 | 0.020 | 38 | 0.003 | 0.910 |
| *Stenurella melanura* | Latest_Logging | 0.000 | 0.000 | 1.107 | 0.275 | 0.000 | 0.001 | 42 | 0.030 | 0.412 |
|  | Dw_diversity | -0.006 | 0.008 | -0.843 | 0.404 | -0.022 | 0.009 | 42 | 0.017 | 0.607 |
|  | V_total | -0.001 | 0.001 | -1.170 | 0.249 | -0.002 | 0.001 | 42 | 0.033 | 0.746 |
|  | Tree_diversity | -0.003 | 0.002 | -1.654 | 0.106 | -0.007 | 0.001 | 42 | 0.064 | 0.319 |
|  | Wood_volume | 0.000 | 0.000 | 1.155 | 0.255 | 0.000 | 0.000 | 42 | 0.032 | 0.612 |
|  | veterans | -0.004 | 0.004 | -0.964 | 0.341 | -0.012 | 0.004 | 42 | 0.023 | 0.910 |
| *Schizotus pectinicornis* | Latest_Logging | 0.001 | 0.000 | 2.065 | 0.048 | 0.000 | 0.002 | 31 | 0.128 | 0.100 |
|  | Dw_diversity | 0.015 | 0.010 | 1.427 | 0.164 | -0.006 | 0.036 | 31 | 0.066 | 0.355 |
|  | V_total | 0.002 | 0.001 | 1.561 | 0.129 | -0.001 | 0.004 | 31 | 0.078 | 0.672 |
|  | Tree_diversity | -0.006 | 0.003 | -1.586 | 0.124 | -0.013 | 0.002 | 31 | 0.080 | 0.319 |
|  | Wood_volume | 0.000 | 0.000 | 0.523 | 0.605 | 0.000 | 0.000 | 31 | 0.009 | 0.806 |
|  | veterans | 0.006 | 0.004 | 1.570 | 0.127 | -0.002 | 0.014 | 31 | 0.078 | 0.910 |
| *Thanasimus formicarius* | Latest_Logging | 0.008 | 0.003 | 3.180 | 0.003 | 0.003 | 0.014 | 36 | 0.229 | 0.038 |
|  | Dw_diversity | 0.091 | 0.057 | 1.580 | 0.123 | -0.026 | 0.207 | 36 | 0.068 | 0.355 |
|  | V_total | 0.025 | 0.018 | 1.409 | 0.168 | -0.011 | 0.061 | 36 | 0.055 | 0.672 |
|  | Tree_diversity | -0.031 | 0.028 | -1.096 | 0.281 | -0.088 | 0.026 | 36 | 0.034 | 0.481 |
|  | Wood_volume | 0.001 | 0.000 | 2.938 | 0.006 | 0.000 | 0.002 | 36 | 0.202 | 0.071 |
|  | veterans | -0.086 | 0.075 | -1.144 | 0.261 | -0.239 | 0.067 | 36 | 0.037 | 0.910 |
| *Ips acuminatus* | Latest_Logging | 0.001 | 0.001 | 1.290 | 0.209 | 0.000 | 0.002 | 26 | 0.065 | 0.359 |
|  | Dw_diversity | -0.007 | 0.017 | -0.421 | 0.677 | -0.042 | 0.028 | 26 | 0.007 | 0.903 |
|  | V_total | 0.000 | 0.006 | 0.065 | 0.949 | -0.011 | 0.012 | 26 | 0.000 | 0.994 |
|  | Tree_diversity | 0.011 | 0.007 | 1.555 | 0.133 | -0.003 | 0.025 | 26 | 0.092 | 0.319 |
|  | Wood_volume | 0.000 | 0.000 | 2.348 | 0.027 | 0.000 | 0.000 | 26 | 0.187 | 0.165 |
| *Ips typographus* | Latest_Logging | 0.000 | 0.000 | 2.022 | 0.050 | 0.000 | 0.001 | 41 | 0.095 | 0.100 |
|  | Dw_diversity | 0.010 | 0.003 | 2.965 | 0.005 | 0.003 | 0.016 | 41 | 0.184 | 0.031 |
|  | V_total | 0.000 | 0.001 | -0.221 | 0.826 | -0.002 | 0.001 | 41 | 0.001 | 0.991 |
|  | Tree_diversity | 0.002 | 0.002 | 1.319 | 0.195 | -0.001 | 0.006 | 41 | 0.043 | 0.390 |
|  | Wood_volume | 0.000 | 0.000 | 1.730 | 0.092 | 0.000 | 0.000 | 41 | 0.071 | 0.366 |
|  | veterans | 0.001 | 0.002 | 0.386 | 0.701 | -0.003 | 0.005 | 41 | 0.004 | 0.910 |
| *Rhagium inquisitor* | Latest_Logging | 0.000 | 0.000 | -0.127 | 0.900 | 0.000 | 0.000 | 34 | 0.001 | 0.900 |
|  | Dw_diversity | -0.001 | 0.008 | -0.068 | 0.946 | -0.017 | 0.016 | 34 | 0.000 | 0.967 |
|  | V_total | 0.000 | 0.001 | -0.371 | 0.713 | -0.001 | 0.001 | 34 | 0.004 | 0.991 |
|  | Tree_diversity | -0.002 | 0.002 | -0.776 | 0.444 | -0.006 | 0.003 | 34 | 0.018 | 0.666 |
|  | Wood_volume | 0.000 | 0.000 | 0.782 | 0.440 | 0.000 | 0.000 | 34 | 0.019 | 0.787 |
|  | veterans | 0.001 | 0.004 | 0.368 | 0.715 | -0.006 | 0.009 | 34 | 0.004 | 0.910 |
| *Phaenops cyanea* | Latest_Logging | -0.001 | 0.001 | -2.285 | 0.031 | -0.003 | 0.000 | 28 | 0.167 | 0.100 |
|  | Dw_diversity | 0.000 | 0.008 | 0.042 | 0.967 | -0.017 | 0.018 | 28 | 0.000 | 0.967 |
|  | V_total | -0.001 | 0.003 | -0.245 | 0.808 | -0.006 | 0.005 | 28 | 0.002 | 0.991 |
|  | Tree_diversity | 0.002 | 0.004 | 0.525 | 0.604 | -0.006 | 0.011 | 28 | 0.010 | 0.732 |
|  | Wood_volume | 0.000 | 0.000 | -0.179 | 0.860 | 0.000 | 0.000 | 28 | 0.001 | 0.938 |
|  | veterans | 0.004 | 0.006 | 0.706 | 0.486 | -0.008 | 0.016 | 28 | 0.019 | 0.910 |

**Table S8.** Sensitivity analysis comparing mean haplotype diversity (H_d_) between managed and primeval forest sites in the **NE region** for each study species (*Elater ferrugineus* is absent as it was sampled only in primeval forest in NE region). The table reports the number of sampling sites in each management category, mean (H_d_), the difference in mean (H_d_) (managed - primeval), nominal P-values, and FDR-adjusted P-values.

| Species | N sites NE | N managed NE | N primeval NE | Mean H_d_ managed NE | Mean H_d_ primeval NE | Δ H_d_ NE | P value | P FDR |
| --- | --- | --- | --- | --- | --- | --- | --- | --- |
| *Neomida haemorrhoidalis* | 16 | 10 | 6 | 0.507 | 0.506 | 0.001 | 0.871 | 0.967 |
| *Boros schneideri* | 20 | 14 | 6 | 0.496 | 0.513 | -0.016 | 0.091 | 0.333 |
| *Cucujus cinnaberinus* | 21 | 16 | 5 | 0.469 | 0.464 | 0.005 | 0.967 | 0.967 |
| *Bolitophagus reticulatus* | 18 | 13 | 5 | 0.499 | 0.481 | 0.018 | 0.490 | 0.851 |
| *Stenurella melanura* | 19 | 14 | 5 | 0.431 | 0.425 | 0.006 | 0.497 | 0.851 |
| *Schizotus pectinicornis* | 16 | 11 | 5 | 0.739 | 0.731 | 0.007 | 0.287 | 0.789 |
| *Thanasimus formicarius* | 20 | 14 | 6 | 0.861 | 0.604 | 0.256 | 0.902 | 0.967 |
| *Ips acuminatus* | 17 | 13 | 4 | 0.472 | 0.435 | 0.037 | 0.777 | 0.967 |
| *Ips typographus* | 27 | 19 | 8 | 0.545 | 0.551 | -0.007 | 0.541 | 0.851 |
| *Rhagium inquisitor* | 19 | 14 | 5 | 0.627 | 0.599 | 0.029 | 0.058 | 0.317 |
| *Phaenops cyanea* | 14 | 10 | 4 | 0.751 | 0.709 | 0.042 | 0.040 | 0.317 |

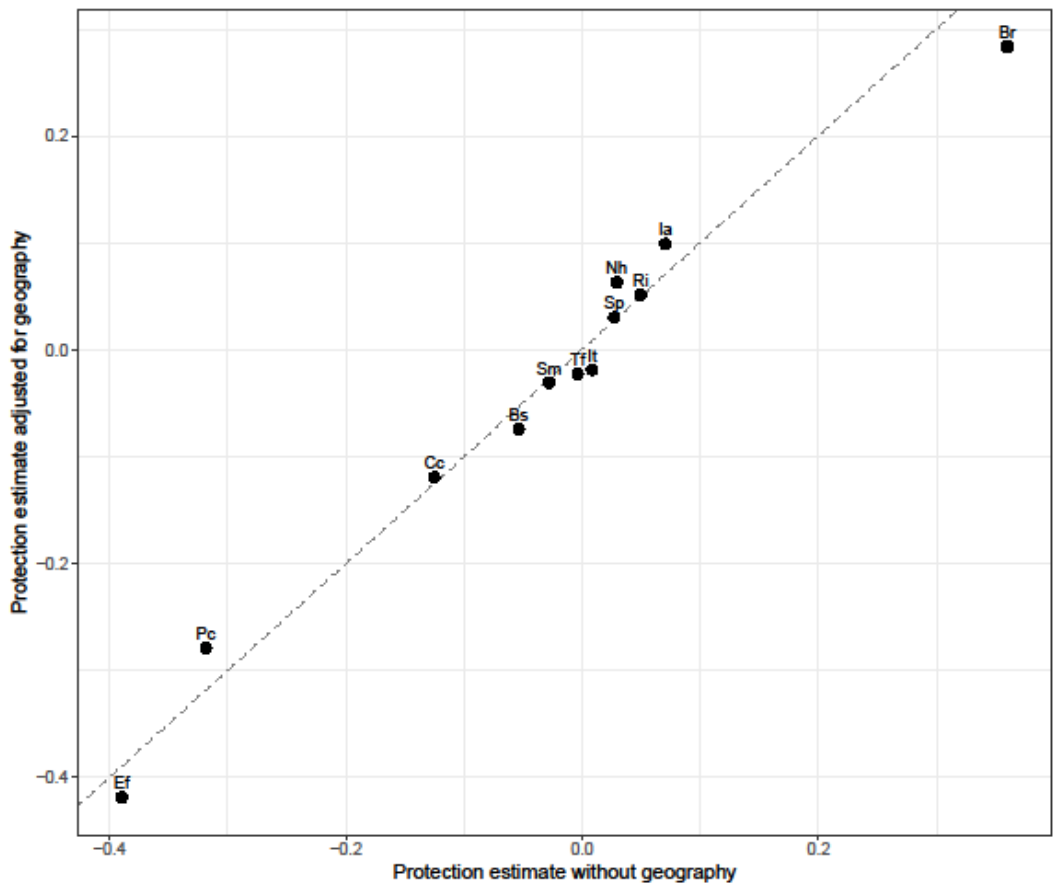

**Figure S1.** Sensitivity of species-specific **protection effects** on individual microhaplotype homozygosity (F_HAP_) to geographic adjustment. Points represent effect estimates from models with and without geographic region adjustment; the dashed line indicates the 1:1 relationship.

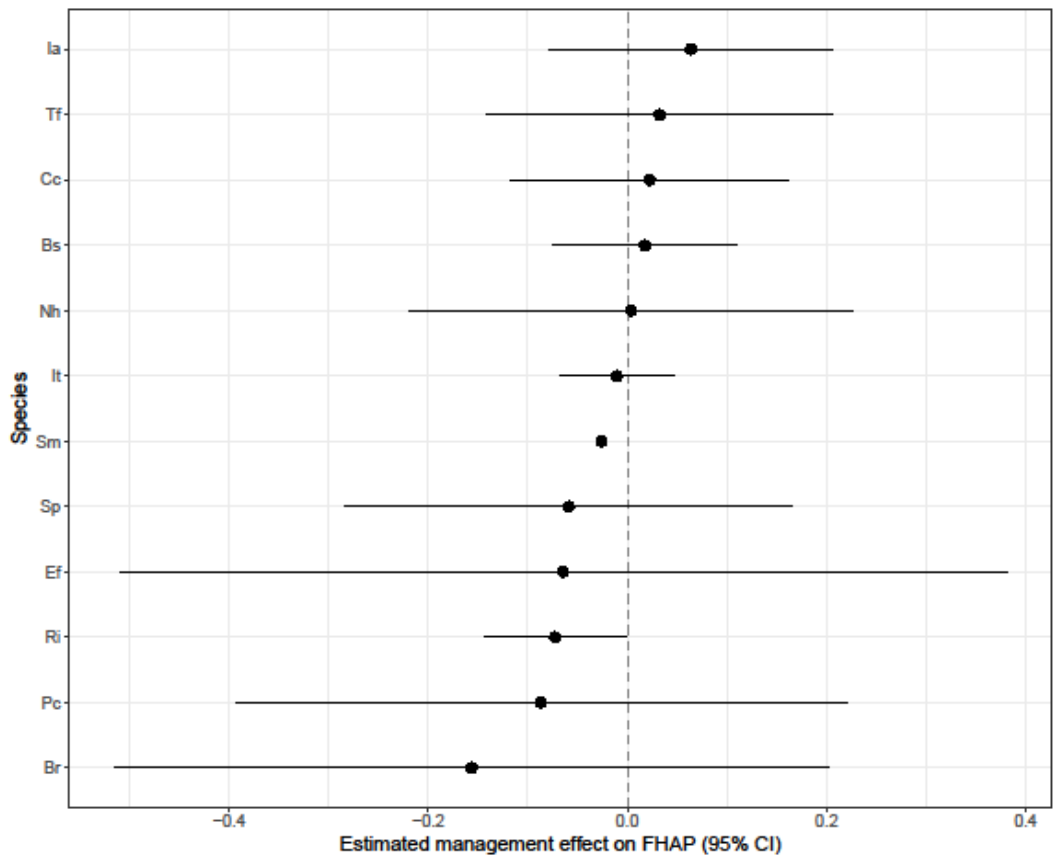

**Figure S2.** Species-specific effects of forest **management type** on individual microhaplotype homozygosity (F_HAP_), adjusted for geographic region. Points represent estimated management effects, and horizontal bars indicate 95% confidence intervals. The vertical dashed line indicates no effect.

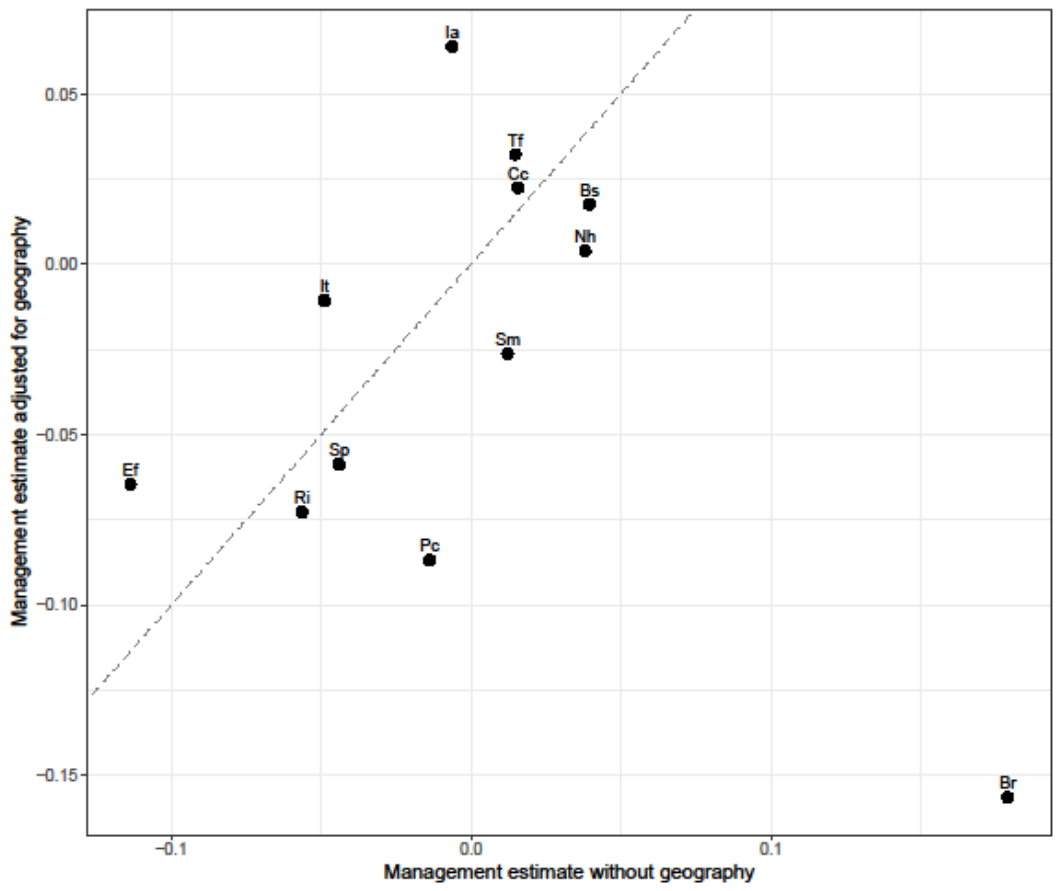

**Figure S3.** Sensitivity of species-specific forest **management effects** on individual microhaplotype homozygosity (F_HAP_) to geographic adjustment. Points represent estimated management effects from models with and without geographic adjustment; the dashed line indicates the 1:1 relationship. Deviations from this line indicate differences in estimated effects between models.
